# A multilocus perspective on the evolutionary dynamics of multistrain pathogens

**DOI:** 10.1101/2023.10.27.564465

**Authors:** David V. McLeod, Claudia Bank, Sylvain Gandon

## Abstract

Many human pathogens, including malaria, dengue, influenza, *Streptococcus pneumoniae*, and cytomegalovirus, coexist as multiple genetically distinct strains. Understanding how these multistrain pathogens evolve is of critical importance for forecasting epidemics and predicting the consequences of vaccination. One factor believed to play an important role is naturally acquired immunity. Consequently, a large body of research has sought to predict how acquired immunity molds the genomics of pathogen populations (i.e., what shapes pathogen strain structure). The diversity of existing models has resulted in conflicting evolutionary predictions, and has sparked an ongoing debate about which predictions are most broadly applicable. Here, we adopt a multilocus population genetics perspective that unifies the predictions of existing models. We identify three key factors that determine the role of naturally acquired immunity in the evolution of pathogen strain structure: (i) the strength and specificity of immune protections, (ii) the dynamic immunological landscape, and (iii) the number of loci coding for the antigens of the pathogen. Isolating and discussing these three factors clarifies the relationship among previous models of multistrain dynamics, and establishes a solid theoretical foundation for the study of the evolutionary epidemiology of multistrain pathogens.

## 1 Introduction

Genomic data of many pathogens of importance for human health, including malaria [1], dengue [2], *Streptococcus pneumoniae* [3], and cytomegalovirus [4], reveal species genetically subdivided into distinct genotypes, or strains, characterized by statistical associations between alleles at different loci [5–8]. Disentangling what produces these genomic patterns (henceforth termed pathogen strain structure or PSS) is a major focus of evolutionary epidemiology, owing to its relevance for a variety of human health concerns, including forecasting epidemics [9] and predicting the epidemiological consequences of vaccination campaigns [10, 11]. One factor believed to play an important role is naturally acquired host immunity, and a large body of work has sought to discern how host immunity affects the population genomics of pathogens [6, 9, 12–20].

The aspect of naturally acquired immunity that has received the most attention is the role of cross-protection, broadly defined as any immune protection an infection provides against genetically similar strains [e.g., 6, 9, 12, 14–16, 21–25]. Roughly speaking, the predictions are as follows [15, 26]: if cross-protection is weak, there is no PSS; if it is of intermediate strength, evolutionary oscillations are common; if it is strong, a set of strains sharing no antigens in common is overrepresented in the population [6, 12]. Yet, each of these predictions comes with exceptions, which has caused debate about the generality of the results. For example, it is not well-understood why only some models produce evolutionary oscillations [27], how predictions of PSS scale with an increasing number of immunogenic loci and alleles [18–20], or what patterns of PSS are either possible or likely [28]. One reason for this debate is that models which combine naturally acquired immunity with pathogen genetic diversity require a large number of variables [29]. Consequently, existing work has taken different approaches to simplify the dynamics, with poorly understood consequences [27, 29, 30].

Here we take a population genetics perspective to review, generalize, and provide evolutionary intuition about when, and why, existing predictions hold (or not). We do so in three parts, each of which emphasizes biological intuition [see e.g., 27, 30, for a more quantitative perspective]. In Section 2, we define PSS. In Section 3, we use this definition to identify the evolutionary forces producing PSS. In Section 4, we show how the identified evolutionary forces are shaped by the epidemiology of naturally acquired immunity. This results in the identification of three crucial factors that determine whether and which type of PSS occurs: the strength and specificity of the immune protections, the dynamic immunological landscape, and the amount of pathogen genetic diversity. We conclude by discussing future directions. Notation and acronyms used are detailed in Table 1.

**Table 1:** Key notation used in main text.

| Notation | Description |
| --- | --- |
| $p_i(t), p_{ij}(t)$ | Frequency of infections by strains carrying allele(s) $i$ and $i, j$, respectively, at time $t$. |
| $D(t)$ | Pairwise linkage disequilibrium (LD) between alleles $A$ and $B$. |
| PSS | Pathogen strain structure; indicates the presence of LD of some order, that is, some strains are overrepresented. |
| DSS | Discordant strain structure; is a specific form of PSS in which a set of strains sharing no alleles is overrepresented. |
| $w_k$ | Additive selection coefficient for allele $k$. Indicates whether allele $k$ is favoured ($w_k > 0$) or disfavoured ($w_k < 0$). |
| $w_{ij}, w_{ijk}$ | Epistasis between alleles $i, j$ and $i, j, k$. If positive (resp. negative), the combination of alleles are more (resp. less) favoured than expected from adding the independent effects of each allele. |
| $\sigma_n, \sigma_h$ | Probability of infection of a host with immunity to $n = 0, 1, 2, \dots$ antigens, or homologous immunity $h$, of the transmitted strain. |
| $\sigma_A$ | Controls the rate of increase of cross-protection as the number of antigens recognized by the immune system increases. If positive (resp. negative), cross protections are saturating (resp. accelerating). |
| $\sigma_S$ | Controls whether homologous immunity is equivalent to (i.e., $\sigma_S = 0$), or more protective than (i.e., $\sigma_S > 0$), full antigenic immunity. |
| $\mathcal{A}_\Delta(t)$ | Indicates whether more uninfected hosts have full antigenic immunity to strains belonging to the set $\{ab, AB\}$ (i.e., $\mathcal{A}_\Delta(t) > 0$) or to the set $\{aB, Ab\}$ (i.e., $\mathcal{A}_\Delta(t) < 0$). |
| $\mathcal{S}_\Delta(t)$ | Indicates whether more uninfected hosts have homologous immunity to strains belonging to the set $\{ab, AB\}$ (i.e., $\mathcal{S}_\Delta(t) > 0$) or to the set $\{aB, Ab\}$ (i.e., $\mathcal{S}_\Delta(t) < 0$). |
| $f_n$ | Proportional to the effective reproductive number of an infection in a host with immunity to $n$ of its antigens. |
| $\epsilon_A$ | Indicates whether an infection in a host with immunity to one of its antigens has a smaller ($\epsilon_A > 0$) or larger ($\epsilon_A < 0$) effective reproductive number than expected. |
| $R_0$ | Basic reproductive number. |
| $\theta$ | Ratio of infectious period to host lifespan. If small, infections are acute; if large, infections are chronic. |

## 2 What is pathogen strain structure (PSS)?

Pathogen strain structure (PSS) refers to the statistical association between alleles at different loci which are neither explainable by chance or by selection on individual alleles. Thus, PSS requires some strains (i.e., some multilocus combination of alleles) to be *overrepresented* relative to their expectation under a random shuffling of alleles, that is, some strains are more frequent than the product of their allele frequencies. Hence, identification of PSS necessitates specifying the multilocus pathogen genetics. For a pathogen genome consisting of two diallelic loci, {*a, A*} and {*b, B*}, corresponding to four strains {*ab, Ab, aB, AB*}, PSS requires

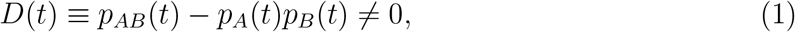

where *p*_*i*_(*t*) and *p*_*ij*_(*t*) denote the frequency of infections by strains carrying allele(s) *i* and *ij*, respectively, at time *t*. In classical population genetics, the quantity *D*(*t*) is pairwise linkage disequilibrium (LD) [31], and is expected to be removed by recombination. The concept of LD generalizes to multiple loci and alleles [31], and since LD captures any nonrandom associations between alleles, the presence of LD is necessary for some strains to be overrepresented. Hence, the minimal definition of PSS is the presence of LD (pairwise or higher-order).

A more stringent form of PSS, *discordant strain structure* (DSS), was first proposed by Gupta et al. [6]. DSS occurs when a set of strains whose members share no alleles (e.g., {*ab, AB*} or {*Ab, aB*}) is overrepresented in the population [6, 12, 32]. The presence of pairwise LD is a necessary but not sufficient condition for DSS [32, 33].

Finally, although PSS requires the presence of LD, in the long term, this could correspond to either LD reaching a stable equilibrium with unchanging strain frequencies or to LD undergoing sustained (potentially chaotic) oscillations, where strain frequencies temporally fluctuate. If LD reaches a non-zero stable equilibrium, we will refer to this as *stable PSS*, whereas if LD undergoes sustained oscillations, we will refer to this as *oscillatory PSS*.

## 3 Which evolutionary forces produce strain structure?

As PSS requires the presence of LD, the dynamics of LD are key to understanding its evolution. To describe these dynamics, we initially focus on two diallelic loci. Moreover, we initially assume that each allele has the same intrinsic fitness, that is, each allele induces the same degree of immune protections. For multistrain models of adaptive immunity without intrinsic fitness differences between alleles, two timescales often emerge [2 7, 34]. On the fast timescale, strong negative frequency-dependent selection (NFDS) operates through the additive selection coefficients at each locus, *w*_*A*_ ≡ *r*_*A b*_ − *r*_*a b*_ and *w*_*B*_ ≡ *r*_*a B*_ − *r*_*a b*_, to push the alleles to an equal frequency, *p*_*A*_(*t*) ≈ *p*_*a*_(*t*) ≈ *p*_*B*_(*t*) ≈ *p*_*b*_(*t*) (here, *r*_*ij*_ is the per-capita growth rate of strain *ij*; see Box 1). At near-equal allele frequencies, the strains necessarily self-organize into two sets {*ab, AB*} and {*Ab, aB*} with *p*_*ab*_(*t*) ≈ *p*_*AB*_(*t*) and *p*_*Ab*_(*t*) ≈ *p*_*aB*_(*t*). Following this self-organization, strains belonging to the same set behave similarly, that is, they have the same per-capita growth rates (i.e., *r*_*AB*_ ≈ *r*_*ab*_ and *r*_*Ab*_ ≈ *r*_*aB*_). As a consequence of *r*_*Ab*_ ≈ *r*_*aB*_, it follows that *w*_*A*_ ≈ *w*_*B*_. Moreover, if we let *w*_*AB*_ denote epistasis in fitness between alleles *A* and *B*, that is,

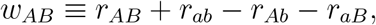

then, using the relations between the per-capita growth rates, we obtain

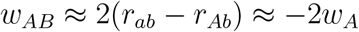

and so *w*_*A*_ ≈ *w*_*B*_ ≈ −*w*_*AB*_*/*2. Consequently, while the allele frequencies are in quasi-equilibrium, in the long term, LD evolves according to the equation

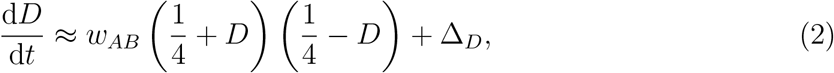

where Δ_*D*_ is the net effect of recombination on LD (Sup. Info. S3 and equations (S22)). Equation (1) and (2) reveal the evolutionary forces that generate or erode PSS.

First, equation (1) implies that the presence of PSS requires both loci to be polymorphic. Maintaining genetic polymorphisms is rarely a problem in models of adaptive immunity, as NFDS will tend to maintain allelic diversity. In fact, most models tend to maximize this diversity; for example, in the case of two diallelic loci without intrinsic fitness differences between alleles, allele frequencies tend to 1*/*2. As our interest is what favours or disfavours some strains given allelic variation observable in genomic data, we will not consider further what favours or disfavours allelic diversity [e.g., see 8, 35, 36].

Second, assuming that both loci are polymorphic, the long-term presence of PSS requires a force that generates LD. Equation (2) yields two possibilities: epistasis and (biased) recombination. Because virtually all multistrain models implicitly revolve around epistasis [but see 37, 38], we will assume that recombination is weak and unbiased, i.e., Δ_*D*_ ≈ −*ρD*, where *ρ* is a (small) positive constant. Whenever the rate of recombination is small relative to the magnitude of epistasis, that is, *ρ* ≪ |*w*_*AB*_|, the dynamics of (2) are driven by epistasis, and its sign indicates whether the combination of alleles *A* and *B* is favored, i.e., *w*_*AB*_ *>* 0, or disfavoured, i.e., *w*_*AB*_ *<* 0.

## 4 Epistasis in an epidemiological context

How is epistasis in fitness shaped by epidemiology? To answer this requires an epidemiological model. When possible, we will use the model of Box 1. Briefly, in this model, we assume that all strains have the same transmission rate *β* and exposure to an allele (or antigen) induces an adaptive immune response in the host that protects against future infections. In particular, the probability of infection of a host with immunity to *n* = 0, 1, 2 antigens of the transmitted strain is *σ*_*n*_, and the probability a host is re-infected by the same strain is *σ*_*h*_ (homologous immunity). Then epistasis is

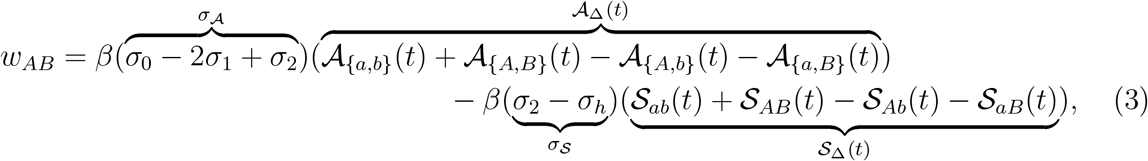

The quantity 𝒜_{*i,j*}_(*t*) denotes the density of uninfected hosts with immunity to *at least* the set of antigens *i* and *j* (𝒜 for *antigen*). In contrast, the quantity 𝒮_*ij*_(*t*) denotes the density of uninfected hosts that have been previously infected by strain *ij* (𝒮 for *strain*; Box 1). Equation (3) shows that two factors contribute to the dynamics of epistasis. The first is the strength and specificity of the immune protection, captured by the composite parameters *σ*_𝒜_ and *σ*_𝒮_ (Fig. 2). The second is the dynamic immunological landscape, captured by the composite variables 𝒜_Δ_(*t*) and 𝒮_Δ_(*t*).

### Box 1

**A population model of naturally acquired immunity**

Models of adaptive immunity largely differ regarding three types of assumptions (Sup. Info. S2): the type of immune protection considered (e.g., does immunity reduce infection or transmissibility?); how immunity is gained and lost; and whether hosts can be simultaneously infected by multiple strains (co-infection). Typically, the choice of assumptions is made to simplify the mathematics. The model we focus on here prioritizes conceptual clarity; extensions and links to other models can be found in the Supplementary Information.

We consider a pathogen whose genome consists of some number of multiallelic loci; each allele, or antigen, induces a (host) adaptive immune response. Transmission occurs exclusively to uninfected hosts (so we do not permit co-infection) via mass-action with the same rate constant *β* for all strains, whereas the probability of successful infection, 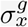, depends upon the host’s immune history, *x*, and the genotype, *g*, of the pathogen strain. In principle, *x* could be any combination of antigens and/or strains and could also account for the temporal order of infections (e.g., more recent infections induce a more protective response). However, here we assume that *x* is an unordered set of prior infections. Thus for *n* pathogen strains, there are 2^*n*^ possible infection histories. For a host with immunity to *n* = 0, 1, … antigens of strain *g* but no prior infection by that strain, 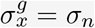, whereas for a host with prior infection by strain *g*, 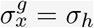 (homologous immunity). For example, for two diallelic loci, a host with immune history *x*^*′*^ = {*ab, Ab*} has 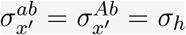 and 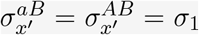. For simplicity, we will assume that *σ*_0_ ≥ *σ*_1_ ≥ … ≥ *σ*_*n*_ ≥ *σ*_*h*_. Hosts clear infections and die from natural and virulence-related mortality at per-capita rates *µ, d*, and *α*, respectively. Immunologically naive hosts (*x* = {∅}) enter the population at rate *b*; immunity is life-long and acquired at clearance. For hosts with immune history *x*, let *S*_*x*_(*t*) denote the density of uninfected individuals and 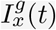 denote the density of individuals infected by strain *g* at time *t*. Then

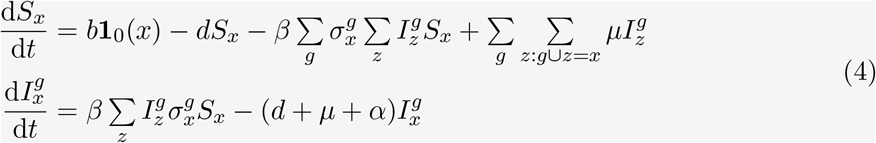

where **1**_0_(*x*) = 1 if *x* = { Ø } and 0 otherwise.

The per-capita growth rates of the different strains link the epidemiological and evolutionary dynamics through the (additive) selection coefficients and epistasis. To extract the selection coefficients and epistasis from model (4), we focus upon two diallelic loci (i.e., *g* = *ij*, where *i* ∈ {*a, A*} and *j* ∈ {*b, B*}). Let 𝒜_{*i,j*}_(*t*) denote the density of uninfected hosts with immunity to *at least* the set of antigens {*i, j*} and 𝒮_*ij*_(*t*) denote the density of uninfected hosts with prior infection by strain *ij* (Fig. 1). For example, uninfected hosts with infection history {*ab, AB*} would contribute to all four of the variables 𝒜_{*a,b*}_(*t*), 𝒜_{*A,b*}_(*t*), 𝒜_{*a,B*}_(*t*) and 𝒜_{*A,B*}_(*t*) but only to the two variables 𝒮_*ab*_(*t*) and 𝒮_*AB*_(*t*). Using this notation, the per-capita growth rate of strain *ij* is

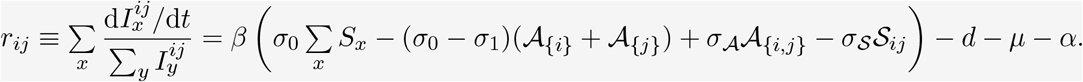

where *σ*_𝒜_ ≡ *σ*_0_ − 2*σ*_1_ + *σ*_2_ and *σ*_𝒮_ ≡ *σ*_2_ − *σ*_*h*_.

**Figure 1:**
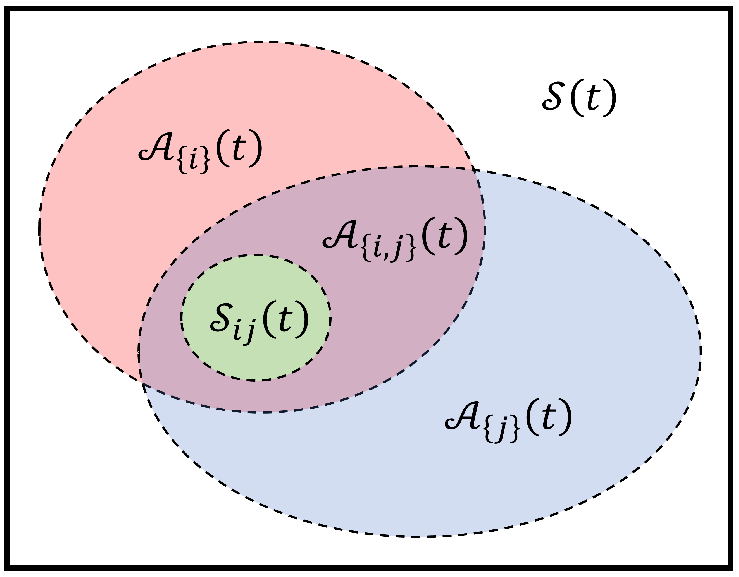
Relationship between variables describing the distribution of immunity among uninfected hosts, where 𝒮 ≡ ∑_*x*_ *S*_*x*_ is the total density of uninfected hosts.

**Figure 2:**
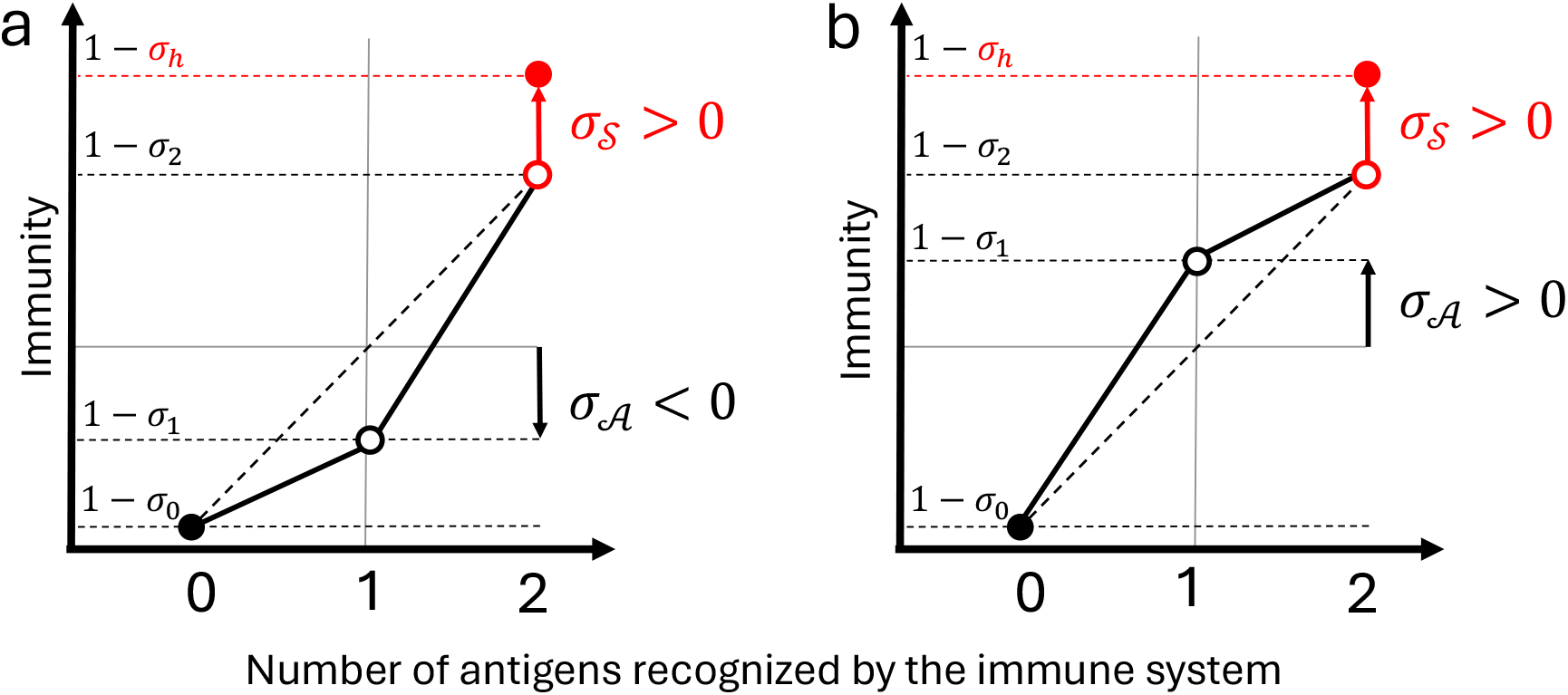
Schematic of cross-protection. Specification of cross-protective immunity translates to two composite parameters controlling how cross-protection affects epistasis. The first composite parameter, *σ*_𝒜_, captures the degree to which the increase in cross-protection is accelerating (*σ*_𝒜_ *<* 0, panel **a**) or saturating (*σ*_𝒜_ *>* 0, panel **b**) as the number of antigens recognized by the immune system increases. The second composite parameter, *σ*_𝒮_, captures whether homologous immunity provides equivalent (*σ*_𝒮_ = 0) or more (*σ*_𝒮_ *>* 0) protection than full antigenic immunity.

### 4.1 The strength and specificity of immune protections

Existing models can be partitioned into two groups based on the specificity and strength of immune protections. Hosts acquire *antigen-specific* immunity in the first group [e.g., 18–20, 39]: here, immunity to strains *Ab* and *aB* translates to immunity to the set of antigens {*a, A, b, B*} and so full protection against strain *AB*. In this case, *σ*_2_ = *σ*_*h*_ and so *σ*_𝒮_ = 0, that is, homologous immunity is no more protective than full antigenic immunity. Consequently, the evolutionary dynamics hinge on *σ*_𝒜_, which controls whether the rate of increase of antigen-specific cross-protection is accelerating (*σ*_𝒜_ *<* 0) or saturating (*σ*_𝒜_ *>* 0) as the number of antigens recognized by the immune system of the host increases (Fig. 2). Models of antigen-specific immunity may apply to, for example, pathogens that temporally express different antigens to which the host develops an immune response (e.g., malaria [18–20], pathogenic *Neisseria* [40]).

Hosts acquire *strain-specific* immunity in the second group [e.g., 6, 9, 12, 14–16, 26, 41]: the level of cross-protection against a particular strain is determined by the most antigenically similar strain that has infected the host, but this is less protective than homologous immunity, that is, *σ*_2_ *> σ*_*h*_ and so *σ*_𝒮_ *>* 0. Thus *σ*_𝒮_ controls the degree to which homologous immunity is more protective than full antigenic immunity (Fig. 2). For example, although ‘strains’ might be defined based on the antigenic variation at a few immunodominant epitopes, strains might also vary at other weakly antigenic sites. If this variation is more similar within a strain than between strains, homologous immunity may be more protective than full antigenic immunity.

More generally, if the adaptive immune system recognizes, and targets, the pathogen at the level of its constitutive antigens, rather than as a unit (i.e., strain), then *σ*_𝒮_ = 0. If, instead, the pathogen is targeted as a unit and cross-protection arises through single-strain similarity, then *σ*_𝒮_ *>* 0. These two perspectives yield very different evolutionary dynamics, as we will see below.

#### Rapidly increasing antigen-specific cross-protection produces stable PSS

When full antigenic immunity is as protective as homologous immunity (i.e. *σ*_2_ = *σ*_*h*_ and *σ*_𝒮_ = 0), epistasis reduces to 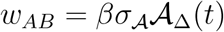, where the variable 𝒜_Δ_(*t*) indicates whether more uninfected hosts have full antigenic immunity to strains belonging to the set {*ab, AB*} (if positive) or to the set {*Ab, aB*} (if negative). There are three possible outcomes, determined by the sign of *σ*_𝒜_. First, if *σ*_𝒜_ *<* 0, all strains stably coexist at equal frequencies. Second, if *σ*_𝒜_ = 0, epistasis vanishes. Hence, any observed PSS is neutral and contingent upon the initial conditions. Third, if *σ*_𝒜_ *>* 0, then stable PSS occurs; in this case, because there is a single epidemiological feedback, 𝒜_Δ_(*t*), one set of strains is competitively excluded. Thus, the sign of *σ*_𝒜_ drives the formation of PSS. In the Supplementary Information S4, we support these predictions with extensive simulations over a range of *R*_0_ ∈ [1.2, 20] and *θ* ∈ [5 *×* 10^−5^, 0.5], where

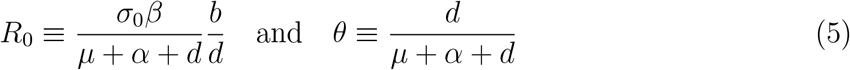

are the basic reproductive number and the ratio of the infectious period to host lifespan [27, 42], respectively. In addition, we analytically show that *σ*_𝒜_ *>* 0 is the necessary condition for stable PSS.

To understand the impact of the sign of *σ*_𝒜_, suppose that, starting from a state in which each strain is equally abundant (*D*(*t*) = 0 and 𝒜_Δ_(*t*) = 0), strains belonging to the set {*ab, AB*} (say) slightly increase in frequency by infecting more hosts (so *D*(*t*) *>* 0, that is, these strains become overrepresented). As these infected hosts clear the infections, this will produce an excess of uninfected hosts with immunity to strains belonging to the overrepresented set of strains {*ab, AB*}, that is, 𝒜_Δ_(*t*) *>* 0. These uninfected hosts have immunity to one of the antigens of the underrepresented strains, {*Ab, aB*} and immunity to zero or two antigens of the overrepresented strains. If *σ*_𝒜_ *<* 0, then *σ*_1_ *>* (*σ*_0_ + *σ*_2_)*/*2, and so these uninfected hosts are more susceptible to infection by a randomly chosen strain from the set of underrepresented strains than from the set of overrepresented strains. Consequently, the underrepresented strains will increase in frequency, returning *D*(*t*) to 0. If *σ*_𝒜_ = 0, then *σ*_1_ = (*σ*_0_ + *σ*_2_)*/*2, and so these uninfected hosts are equally likely to be infected by any strain, and therefore *D*(*t*) will not change any further. Finally, if *σ*_𝒜_ *>* 0, then *σ*_1_ *<* (*σ*_0_ + *σ*_2_)*/*2, and so these uninfected hosts are more susceptible to infection by a randomly chosen overrepresented strain. Consequently, the overrepresented strains will further increase in frequency, creating a positive feedback loop that will push *D*(*t*) → 1*/*4.

This analysis assumes that there are no intrinsic fitness differences between alleles or strains. However, some intrinsic variation may be expected in the ability of the host immune system to target specific antigens or strains [43, 44]. If we suppose alleles and/or combinations of alleles vary in the strength of induced immune protections (i.e., *σ*_1_ and *σ*_2_ depend on the identity of the allele(s)), then each allelic combination {*i, j*} will have a specific value of *σ*_𝒜_, i.e., 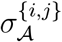. If we define 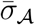 as the average of 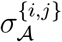, numerical simulations indicate that 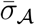 remains a strong predictor of whether PSS occurs or not, provided allelic variation is not too large (Sup. Info. S7). Unsurprisingly, the accuracy of 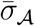 as an evolutionary predictor varies with its value: 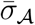 is most accurate when strongly positive (predicting stable PSS), and is least accurate when 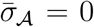. The reason why the prediction breaks down when 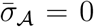 is because epistasis is the sum of the product of the allele combination specific 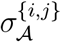 parameters and the dynamic immunological landscape, 𝒜_{*i,j*}_ (equation (S49)). Thus epistasis need not be zero even if 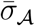 is.

#### Strongly protective homologous immunity generates oscillatory PSS and maintains strain diversity

Importantly, the sign of *σ*_𝒜_ alone cannot sustain evolutionary oscillations (oscillatory PSS), nor can it stably maintain an intermediate level of stable PSS (i.e., 0 *<* |*D*(*t*)| *<* 1*/*4); two common predictions of multistrain models. This is not surprising: when *σ*_𝒮_ = 0, there is a single epidemiological feedback on epistasis, whether more hosts have *antigenic immunity* to the combination of antigens of strains belonging to one set of strains or the other, 𝒜_Δ_(*t*). Consequently, there is no reason to expect the sign of epistasis to change based on the amount of LD. However, if homologous immunity is more protective than full antigenic immunity, then *σ*_𝒮_ *>* 0. This creates a second epidemiological feedback, whether more hosts have *homologous immunity* to strains belonging to one set or the other, 𝒮_Δ_(*t*). The second feedback promotes strain diversity because overrepresented strains will have caused more infections, putting them at a competitive disadvantage. If *σ*_𝒜_ *<* 0, both feedbacks prevent strains from becoming overrepresented, leading to a stable coexistence of all strains at an equal frequency (Fig. 3). However, if *σ*_𝒜_ *>* 0, in principle promoting PSS, this will be counteracted by the feedback created by *σ*_𝒮_ *>* 0, which inhibits PSS (Fig. 3).

**Figure 3:**
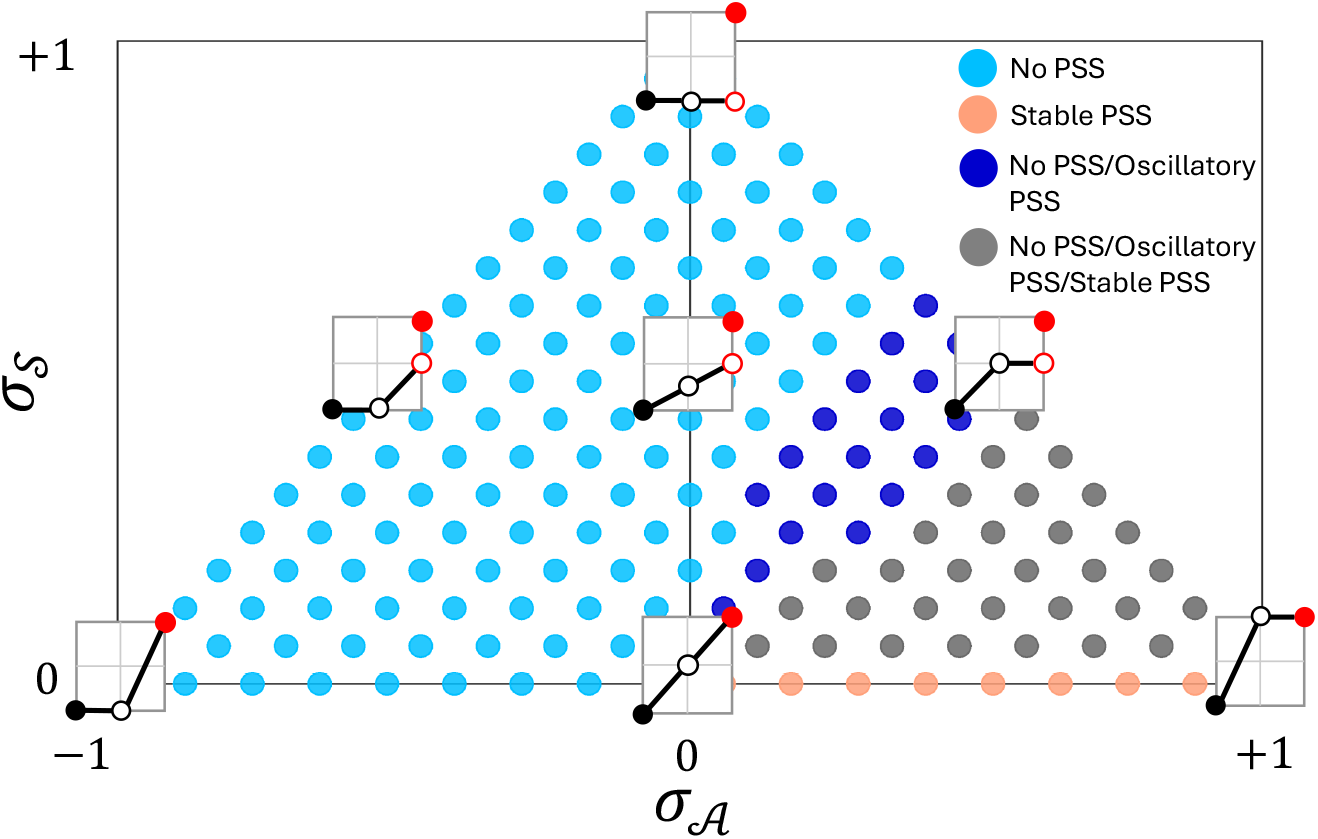
The strength and specificity of cross-protection determines when PSS can form. The strength and specificity of cross-protection is controlled by *σ*_𝒮_ and *σ*_𝒜_ (see Fig. 2); as they vary, this permits the long-term existence of stable or oscillatory PSS or some combination of both. If *σ*_𝒜_ *<* 0, no PSS is expected, regardless of the value of *σ*_𝒮_ (light blue dots). If *σ*_𝒮_ = 0 and *σ*_𝒮_ *>* 0, stable PSS is expected (light red dots). In between these extremes, different outcomes are possible depending upon the epidemiological parameters (i.e., *R*_0_ and *θ*). In particular, if 0 *< σ*_𝒜_ *< σ*_𝒮_, both oscillatory PSS and no PSS are possible, but not stable PSS (dark blue dots), whereas if 0 *< σ*_𝒮_ *< σ*_𝒜_, no PSS, stable PSS, and oscillatory PSS are possible (gray dots). Each circle represents the long-term outcome of 100 simulations of equation (4), where we loop over 10 evenly-spaced values of *R*_0_ ∈ [1.2, 18] and 10 evenly-spaced values (on log-scale) of *θ* ∈ [10^−4^, 0.39]. Each square inset shows how immunity varies in (*σ*_𝒜_, *σ*_𝒮_) space using the same format as Figure 2. Note that the values of *σ*_𝒜_ and *σ*_𝒮_ are constrained due to the assumption that *σ*_0_ ≥ *σ*_1_ ≥ *σ*_2_ ≥ *σ*_*h*_. Representative dynamical behaviour through time for the different cases is shown in Figure S2. All simulations use parameters *b* = *d* = 0.01, *α* = 0, *σ*_0_ = 1, *σ*_*h*_ = 0, and *ρ* = 10^−9^.

The counteracting forces of *σ*_𝒜_ *>* 0 and *σ*_𝒮_ *>* 0 can produce oscillatory PSS, where the amount of PSS undergoes sustained oscillations (potentially chaotic), and numerical simulations indicate *σ*_𝒮_ *>* 0 remains a necessary condition for oscillations even when there is intrinsic variation in fitness of alleles (Sup. Info. S7). Indeed, the divergence between homologous immunity and full antigenic immunity is not only the source of the complex dynamics seen in many models [6, 9, 12, 16, 45], it also explains why oscillations are absent from so-called ‘status-based’ models but not from ‘history-based’ models [13, 27, 30, 34]. History-based models, such as the model in Box 1, assume that following infection, hosts may be partially protected against genetically similar strains (i.e., they have a reduced, but non-zero probability of infection). On the other hand, status-based models assume that following infection, hosts become either completely protected or unprotected against each strain, with the probabilities dependent on immune history (Sup. Info. S2). This is sometimes referred to as *polarized immunity*, as for each strain, the population is divided into either naive hosts or hosts with sterilizing immunity (i.e., if the host recognizes *any* antigens, it cannot be infected, so *σ*_*n*_ = 0 for *n* ≥ 1 and *σ*_*h*_ = 0). Consequently, for status-based models, there is no difference between full antigenic or homologous immunity (so *σ*_𝒮_ = 0), and so no potential for sustained evolutionary oscillations.

#### Strain structure depends on the life-history effects of immune protection

Of course, immunity need not only protect against infection. In the model of Box 1, any of transmissibility, *β*, clearance rate, *µ*, and virulence mortality, *α*, could depend upon the immune history of the host. Although in this case epistasis can still be written as a time-varying quantity as in equation (3), the effective reproductive number of an infection will depend on its current environment (i.e., the immune history of the host). Consequently, the per-capita growth rate of strain *ij* is the weighted average across possible environments, where the weights are the frequency of strain *ij* infections by environment (Sup. Info. S5). As such, time-varying epistasis is less amenable to interpretation, and so here we focus on epistasis at equilibrium. Suppose homologous immunity is no more protective than full antigenic immunity. Then, the population will evolve to a stable equilibrium, whether it be stable PSS or no strain structure. At the stable PSS equilibrium, epistasis and LD have the same sign if

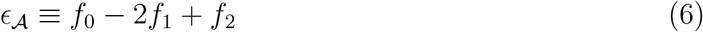

is positive (Sup. Info. S5). In equation (6), 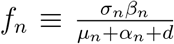 is proportional to the effective reproductive number of an infection in a host with immunity to *n* of its antigens (Sup. Info. S5). In other words, the quantity *ϵ*_𝒜_ measures how cross-immunity increases with the number of antigens recognised by the immune system when immunity affects multiple traits of the pathogen (*ϵ*_𝒜_ is proportional to *σ*_𝒜_ when immunity affects only *σ*). Moreover, the sign of *ϵ*_𝒜_ captures whether the increase in cross-protection is accelerating (*ϵ*_𝒜_ *<* 0, Fig. 2**a**) or saturating (*ϵ*_𝒜_ *>* 0, Fig. 2**b**) as the number of antigens recognized by the immune system increases.

Equation (6) reveals that the life-history effects of immune protection alter the evolutionary predictions in two ways. First, although increasing the protection offered by immunity to one antigen promotes PSS if immunity reduces susceptibility and/or transmissibility [6, 12, 15, 46] or increases clearance rate [18–20, 39], it inhibits PSS if immunity reduces virulence mortality [47, 48]. Note, however, that immune protections against virulence mortality alone are insufficient to maintain genetic polymorphisms (Sup. Info. S5). Second, if each antigen produces an additive increase in the strength of immunity [8], and and we suppose only one type of immune protection is present, then if immunity reduces susceptibility to infection or transmissibility, *ϵ*_𝒜_ = 0, whereas if immunity increases clearance rate (a common assumption for malaria models [18–20]), *ϵ*_𝒜_ *>* 0. Thus, immunity that increases the clearance rate is more likely to produce PSS. This is particularly true when pathogens produce acute infections (smaller *θ*) and/or do not cause host death (smaller *α* for fixed *θ*) (Sup. Info. S5).

### 4.2 The dynamic immunological landscape

In addition to the strength and specificity of immune protections, epistasis also depends upon the dynamic immunological landscape controlled by 𝒜_Δ_(*t*) and 𝒮_Δ_(*t*). When models become more complex than that of Box 1, this landscape plays an increasingly prominent role. However, because these complexities typically alter the dynamics of uninfected hosts, the analysis of equation (3) helps understand the influence of the immunological landscape on the presence of PSS. In particular, this analysis illustrates the impact of re-infections and how immunity is gained and lost.

#### The frequency of re-infections shapes the evolutionary dynamics

When both epidemiological feedbacks, *βσ*_𝒜_𝒜_Δ_(*t*) and *βσ*_𝒮_ 𝒮_Δ_(*t*), are acting on (3), the sign and dynamical behaviour of epistasis depend upon the difference between the distribution of immunity to different antigens, 𝒜_Δ_(*t*), and different strains, 𝒮_Δ_(*t*). We show in Sup. Info. S3.2 that only hosts with immunity to the antigens defining a single strain contribute to 𝒜_Δ_(*t*), whereas 𝒮_Δ_(*t*) depends upon hosts with immunity to more than one strain. Thus, the ‘length’ of the immune history, that is, the number of strains to which a host currently has immunity, plays an important role. If most hosts have immunity to only one strain, then 𝒜_Δ_(*t*) ≈ 𝒮_Δ_(*t*). In this case, equation (3) shows that stable PSS occurs if *σ*_𝒜_ *> σ*_𝒮_ ; otherwise no PSS occurs. If almost all hosts have immunity to more than one strain, then 𝒜_Δ_(*t*) → 0 much faster than 𝒮_Δ_(*t*) → 0. In this case, the epidemiological feedback *σ*_𝒮_ 𝒮_Δ_(*t*) dominates, and as this favours rarer strains, no PSS occurs. In between these extremes, stable PSS, no PSS, or oscillatory PSS are all possible outcomes. Thus, a necessary but not sufficient condition for evolutionary oscillations is that reinfection is possible but not too frequent.

In addition to the strength and specificity of immune protections, two generic pathogen life-history quantities affect the relationship between 𝒜_Δ_(*t*) and 𝒮_Δ_(*t*). The first is the basic reproductive number, *R*_0_, and the second is the ratio of the infectious period to host lifespan, *θ* [27, 42] (see equation (5)). Generally speaking, the ‘length’ of immune history is expected to increase with *R*_0_ and decrease with *θ* (see equations (S5) and (S6)). Thus, when *R*_0_ is small and/or infections are chronic (*θ* ≈ 1) and co-infection is not possible, hosts are unlikely to be infected more than once during their lifespan, so 𝒜_Δ_(*t*) ≈ 𝒮_Δ_(*t*). Consequently, if *σ*_𝒜_ *> σ*_𝒮_, stable PSS will occur (Fig. 4**a**), otherwise no PSS is expected. As *R*_0_ becomes large for a given *θ*, hosts are commonly infected multiple times during their lifespan, thus 𝒜_Δ_(*t*) → 0 faster than 𝒮_Δ_(*t*) → 0, and so no PSS is expected (Fig. 4**a**). Finally, although *σ*_𝒮_ *>* 0 and an ‘intermediate’ *R*_0_ are necessary conditions for evolutionary oscillations, these conditions are not sufficient. In addition, infections need to be highly acute (small *θ*; Fig. 4). This is because oscillations arise through a rapid divergence between 𝒮_Δ_(*t*) and 𝒜_Δ_(*t*) (Fig. 4**c**). As both of these quantities involve uninfected hosts, it is not sufficient for reinfections to occur in order for evolutionary oscillations, the infections must also be quickly cleared.

**Figure 4:**
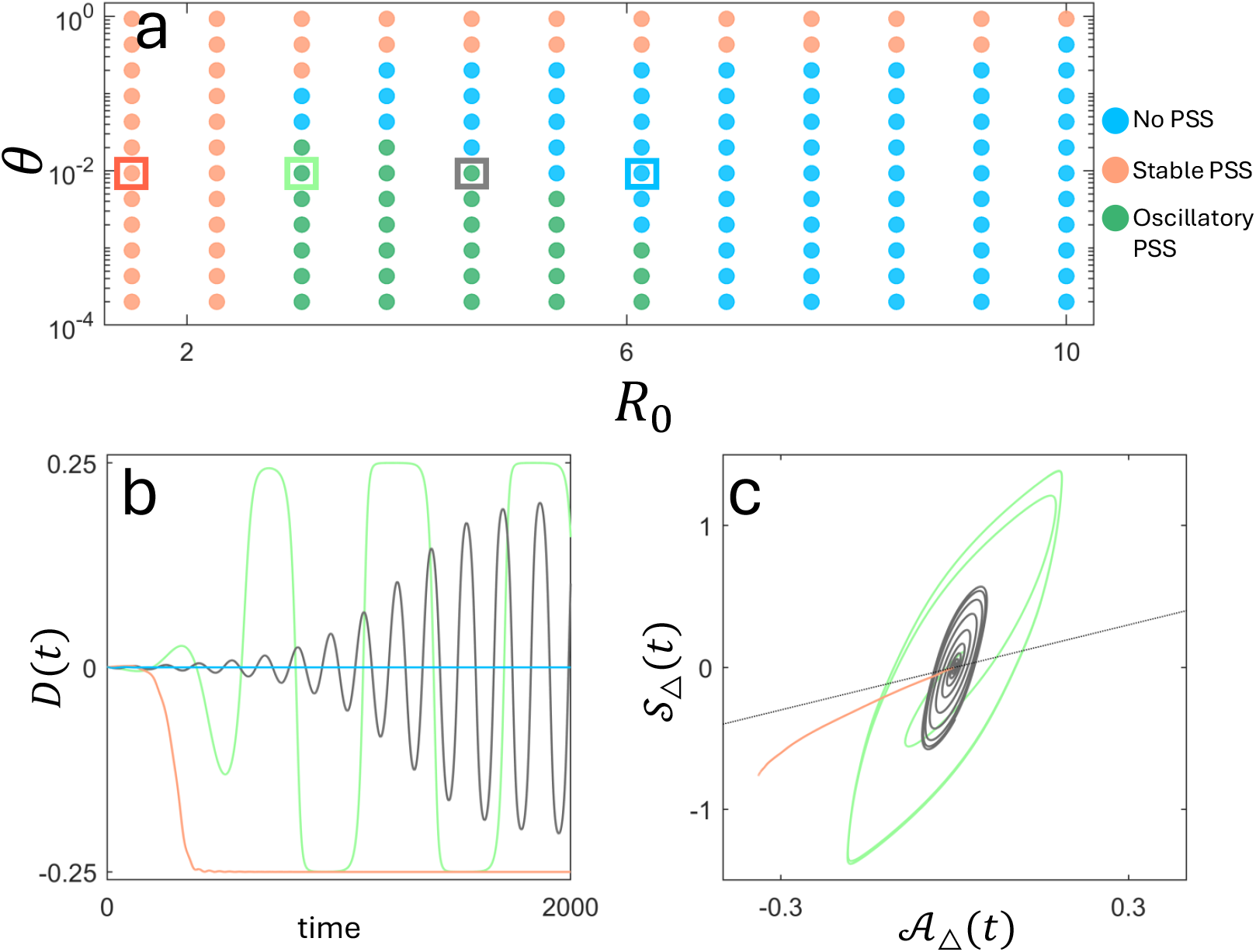
The dynamic immunological landscape. Panel **a** shows the long-term evolutionary outcome as *R*_0_ and the ratio of infectious period to host lifespan, *θ*, vary for fixed *σ*_0_ = 1, *σ*_1_ = 0.35, *σ*_2_ = 0.25, *σ*_*h*_ = 0.15 (so *σ*_𝒜_ = 0.55 and *σ*_𝒮_ = 0.1). When *R*_0_ is small and/or infections are increasingly chronic (large *θ*), 𝒜_Δ_(*t*) ≈ 𝒮_Δ_(*t*), and since (for this example) *σ*_𝒜_ *> σ*_𝒮_, stable PSS occurs (red circles). For a given *θ*, as *R*_0_ increases, it becomes likely that hosts are infected multiple times during their lifespan; here, 𝒮_Δ_(*t*) becomes larger than 𝒜_Δ_(*t*), which destabilizes stable PSS. For acute infections (small *θ*), this can lead to oscillatory PSS (green circles) before sufficiently high *R*_0_ leads to no PSS (blue circles), whereas for more chronic infections, the increase in *R*_0_ leads to a transition from stable PSS to no PSS without oscillations. Panels **b** and **c** show example dynamical behavior for four different values of *R*_0_ and fixed *θ* (indicated by the coloured squares in panel **a**). Note that in panel **c** the initial condition is 𝒜_Δ_(0) = 𝒮_Δ_(0) = 0. All simulations used parameters *b* = *d* = 0.01, *α* = 0, *ρ* = 10^−8^, with *β* and *µ* set by the values of *R*_0_ and *θ*.

Additional factors may affect the ‘length’ of the immune history and may thus act on the dynamic immunological landscape. For example, if immunity wanes over time [49] the length of the immune history will be shortened as hosts ‘forget’ previous infections. Likewise, if there is short-term strain-transcending immunity [17, 41, 50–54] (analogous to no co-infection), the length of immune history will be shortened as the likelihood of becoming re-infected is reduced. If *R*_0_ is very high, shortening the immune history will increase the density of hosts contributing to 𝒜_Δ_(*t*). This increases the likelihood of both oscillatory and stable PSS. If *R*_0_ is intermediate, the likelihood of oscillations will be reduced. This dependence upon *R*_0_ explains why shortening immune history through waning immunity [49], conserved immune responses [55], or strain-transcending immunity [41] has been observed to both decrease [49, 55] and increase [41] the likelihood of oscillations.

#### The impact of cross-protection depends on the gain and loss of immunity

The predicted consequences of the sign of *σ*_𝒜_ hinge upon the expectation that the distribution of infections is coupled with the immunological landscape. That is, when hosts acquire immunity, they acquire immunity to all the antigens of the infecting strain, and when hosts lose immunity, they lose immunity to all the antigens of the previously infecting strain. This tends to create a positive correlation between the sign of *D*(*t*) and the immunological landscape, 𝒜_Δ_(*t*) (Fig. S3). However, this relationship can be disrupted if a host infected by strain *ij* does not always simultaneously acquire and/or lose immunity to the combination of antigens {*i, j*}. For example, suppose host heterogeneity in antigen-presenting cells means that hosts are more likely to acquire immunity to only a single, randomly chosen antigen of an infecting strain rather than both. Moreover, suppose that the set of overrepresented strains is {*ab, AB*}, that is, *D*(*t*) *>* 0. Naive hosts are then most likely to become infected by one of the overrepresented strains (say *AB*), after which they are most likely to become infected by the other overrepresented strain, *ab*, as it is more abundant. But if each infection provides immunity to a single antigen, following the second infection, the host will have immunity to either the set of antigens {*A, b*} or {*a, B*}. Therefore, the host will have full antigenic immunity to one of the underrepresented strains, {*Ab, aB*}. Consequently, this will produce an excess of hosts with immunity to the antigens of one of the underrepresented strains, that is, 𝒜_Δ_(*t*) will have the opposite sign to *D*(*t*). Thus, from equation (3), PSS can only occur if *σ*_𝒜_ *<* 0, i.e., if the increase in cross-protection is accelerating (Sup. Info. S7).

Indeed, there exist a variety of immunological reasons why hosts may not simultaneously gain and/or lose immunity to the combination of antigens {*i, j*}. For example, host variation in the adaptive immune response might mean that different hosts target and/or lose immunity to different antigens, based upon host genetics (e.g., MHC alleles [8]) or randomly (e.g., which *B*-cell encounters which epitope first; see [56] and Sup. Info. S7 for a longer discussion). Importantly, this heterogeneity can disrupt the positive correlation between the distribution of infections and the immunological landscape, reversing predictions about the consequences of the sign of *σ*_𝒜_.

### 4.3 The number of loci and the possible patterns of PSS

Many of the key predictions from our model of two diallelic loci hold as the number of alleles and number of loci increase: PSS is promoted if the increase in cross-protection saturates as the number of antigens recognized by the host increases; homologous immunity being more protective than full antigenic immunity promotes strain diversity and is necessary for evolutionary oscillations; and oscillations are more likely when hosts tend to be infected sequentially by more than one strain during their lifespan. What changes is that as the number of loci increases, the population need not self-organize into sets of strains whose members have no antigens in common (DSS). As a consequence, different types of PSS are possible [9, 16, 28, 41]. This is because LD is no longer restricted to non-random associations spanning two loci. Instead, LD can involve as many loci as are segregating, with each order of LD corresponding to a different type of PSS. Here, order of LD refers to the number of loci across which the non-random allelic association is measured; see Sup. Info. S9. For example, triple LD is an overrepresentation of strains whose members share at most one antigen (e.g., in the case of three segregating loci, one such set of strains is {*abC, aBc, Abc, ABC*}). Generally, the number of overrepresented strains increases with the order of LD. Although the evolutionary dynamics are more complicated, in the long-term, at most one order of LD will be nonzero and evolve. This evolution is still dictated by epistasis, whether it be pairwise or higher-order (Sup. Info. S9).

When full antigenic immunity provides as much protection as homologous immunity, each order of epistasis hinges upon the rate of increase of cross-protection. For *N* loci, this rate is captured by the *N* − 1 composite parameters (Fig. 5a),

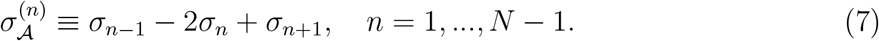

As before, the sign of 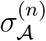 determines whether the rate of increase of cross-protection against *n* antigens is accelerating, 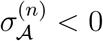, or saturating, 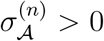. Thus, these parameters control how the strength of competition between strains changes based on genetic similarity, shaping epistasis (Fig. 5). For example, for three diallelic loci, triple epistasis in fitness, *w*_*ABC*_, can maintain triple LD if 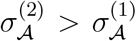. In contrast, pairwise LD between alleles *i* and *j* can be maintained if pairwise LD and *w*_*ij*_ + *w*_*ABC*_ */*2 share sign, which can be approximated as 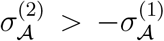 (Sup. Info. S9). One important consequence of multiple possible types of PSS is that the conditions permitting different orders of LD to evolve can be simultaneously satisfied. When this occurs, an evolutionary multistability emerges such that the type of PSS present depends on the initial conditions (Fig. 5; [see also 28]).

**Figure 5:**
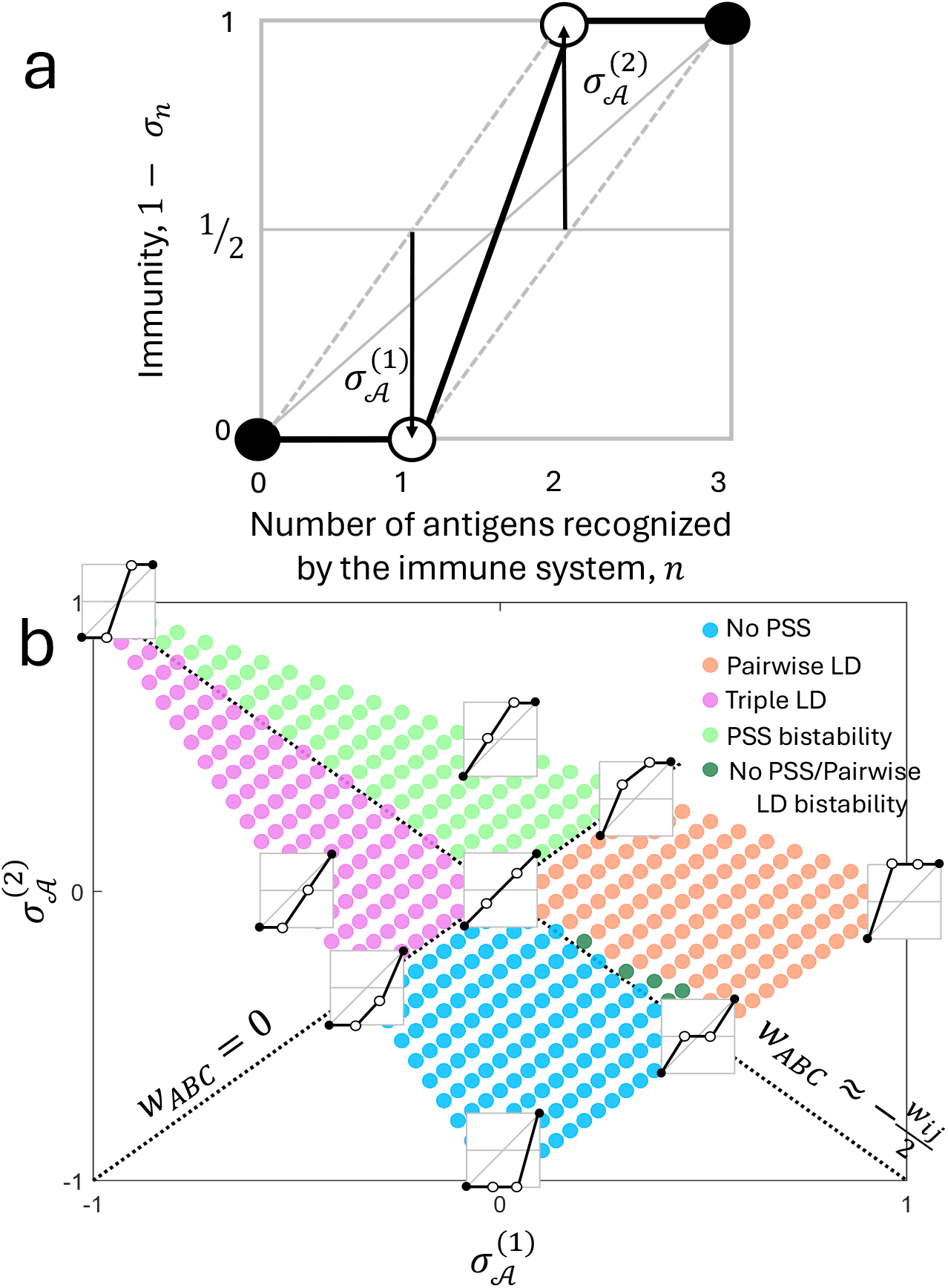
As the number of loci increases, so do the possible types of PSS. The number of loci determines the number of parameters capturing the shape of the function that determines how cross-protections increase with the number of antigens recognized by the immune system (panel **a**). As these parameters vary, different long-term outcomes are possible (panel **b**; see Fig. S11 for the representative dynamics through time). When 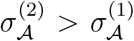, triple LD has the same sign as *w*_*ABC*_ and so can be stably maintained. When 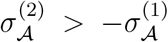, pairwise LD between alleles *i* and *j* will generally have the same sign as *w*_*ABC*_ + *w*_*ij*_*/*2 (Sup. Info. S9) and so can be stably maintained. As both conditions can be simultaneously met, evolutionary bistability can occur, where the type of PSS that evolves depends upon the initial conditions. Our assumption of *σ*_0_ ≥ *σ*_1_ ≥ *σ*_2_ ≥ *σ*_3_ constrains the possible values that 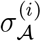 can take in panel **b**. Because *σ*_*S*_ = 0, when PSS occurs, it is stable. Parameter values used in panel **b**: *β* = 1, *d* = *b* = 0.01, *α* = 0, *µ* = 0.4, *ρ* = 10^−7^, *σ*_0_ = 1, *σ*_3_ = *σ*_*h*_ = 0, whereas *σ*_1_ and *σ*_2_ vary.

Given *N >* 2 loci and the possibility of multiple types of PSS, what genomic patterns are expected? This hinges upon the number of antigens sufficient to induce a strongly protective response. For example, if immunity to two antigens is strongly protective, 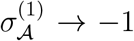 and 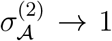, triple LD is more likely than pairwise LD (Fig. 5). If instead immunity to one antigen is strongly protective, 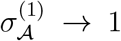 and 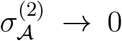, then pairwise LD is more likely (Fig. 5). This explains why models assuming that immunity to one antigen is strongly cross-protective (i.e., *σ*_1_ = *σ*_2_ = … = *σ*_*n*_, and *σ*_*n*_ → *σ*_*h*_) predict DSS and pairwise LD [6, 12].

## 5 Discussion

Knowledge of when, and why, naturally acquired immunity produces non-random genomic patterns in pathogen populations (pathogen strain structure or PSS) is critical for understanding and predicting pathogen dynamics. Here we review and generalize predictions for the evolution of PSS by focusing on linkage disequilibrium (LD) and the forces that drive its dynamics. In doing so, we establish the evolutionary connections between the multitude of existing models. In particular, our analysis illustrates the factors which determine (i) the long-term presence of PSS as a stable equilibrium (stable PSS) and (ii) the long-term presence of PSS as evolutionary oscillations (oscillatory PSS).

The long-term presence of stable PSS hinges upon two factors. First, stable PSS is more likely when the increase in cross-protection is saturating (*σ*_𝒜_ *>* 0) and prior exposure to *all* antigens of a given strain provides similar protection to homologous immunity (*σ*_𝒮_ ≈ 0). Second, stable PSS is more likely if hosts have immunity to only a few strains. This occurs for pathogens with a low-to-intermediate *R*_0_, pathogens causing chronic infections with limited co-infection, or when hosts display either waning or strain-transcending immunity. The most likely type of stable PSS, expressed through the non-zero order of LD, depends on the number of antigens required to induce a strongly protective immune response. As this number increases, so will the expected non-zero order of LD, and thus the fraction of possible strains present in the population. This implies that, although each type of PSS is in principle detectable from genomic data, it is necessary to identify the appropriate order of LD. For example, a highly structured population dominated by triple LD could appear unstructured based on the assessment of pairwise LD.

We showed that the shape of the function specifying cross-protection (see Fig. 2 and Fig. 5**a**) controls the amount of genotypic diversity which provides testable predictions. For example, vaccine challenge studies of influenza A virus (IAV) in humans and equine influenza virus (EIV) in horses indicate that cross-protection increases roughly linearly with antigenic distance for IAV, but the increase is saturating for EIV (see Fig. 1a in [57]). Consequently, our model would predict that, all else being equal, EIV should be less *genotypically* diverse than IAV. Consistent with our prediction, EIV can be roughly divided into three antigenic clusters [58], whereas IAV consists of 11 antigenic clusters [59] (see also [60]). Notably, EIV and IAV differ with respect to other properties (e.g., IAV undergoes more rapid antigenic drift [60] and the human hosts of IAV have a longer lifespan than the equine hosts of EIV). Quantitatively evaluating how the shape of the function specifying cross-protection limits genotypic diversity thus warrants further work.

Evolutionary oscillations, or oscillatory PSS, have been a focal point of research for multistrain pathogens [e.g., 6, 9, 12, 16, 45]. Our multilocus perspective reveals that oscillatory PSS is most likely when three conditions are met (Fig. 3). First, immunity to each additional antigen provides a larger-than-additive gain in protection (*σ*_𝒜_ is positive), but only homologous immunity triggers a strongly protective immune response (so *σ*_𝒮_ is positive). Second, hosts are likely to be infected by, and so have homologous immunity to, multiple strains over their lifespan. This is most likely for pathogens with either an intermediate *R*_0_, or those with a high *R*_0_ in the presence of waning or strain-transcending immunity (i.e., something shortening the ‘length’ of immune history). Third, infections are acute (*θ* is small), creating a rapid divergence between the distribution of hosts with immunity to certain antigens and the distribution of hosts with immunity to certain strains. Whether these conditions are met or not will depend upon the pathogen species. We propose that identifying these key biological quantities may have greater utility and be more empirically feasible than identifying regions of parameter space for a given model that permit oscillations. For example, Recker et al. [9] hypothesized that rather than being a mutation-driven process [13, 14, 52], influenza evolution may be better explained by the periodic recycling (or oscillations) of a finite set of antigens (but see [36]). This argument was supported by oscillations occurring over a wide range of parameter space. Future work could evaluate whether influenza meets the biological conditions for displaying such behaviour.

### Co-infection and within-host interactions

The model of Box 1 assumes that transmission only occurs to uninfected hosts. It thus ignores within-host interactions. Multistrain models frequently assume that hosts can be simultaneously infected by any number of *different* strains (co-infection; see Table S2), each of which is unaffected by the presence of other co-infecting strains [e.g., 6, 12, 15, 41, 54, 61, 62]. Under these assumptions, and by redefining certain variables in the model of Box 1, all our key predictions hold (Sup. Info. S2.3). However, assumptions maintain very high levels of genetic diversity, including otherwise selectively disadvantageous trait variation [e.g., hypervirulence 22]. Although a previous simulation study found limited effects of within-host interactions [18], this may not hold generally. For example, co-infection in *Streptococcus pneumonia* can lead to the simultaneous clearance of all infecting strains [8, 63, 64]. This clearance will disproportionately (negatively) affect the more abundant strains, potentially removing PSS which may partially explain why *S. pneumoniae* genomic data show low amounts of strain structure [65]. Thus, the impact of within-host interactions on PSS warrants further work.

### Demographic stochasticity

Our analysis is based on a deterministic model that neglects demographic stochasticity, which was previously shown to affect multistrain dynamics in two ways. First, demographic stochasticity can produce random fluctuations in allele and strain frequencies not seen in a deterministic model [18, 66]. Demographic stochasticity can generate LD, cause LD to drift in a quasi-neutral fashion, and/or trigger infrequent oscillations between strains, thus affecting PSS, especially when epistasis is weak. Indeed, demographic stochasticity coupled with weak epistasis is likely the source of the quasi-stable PSS observed in some stochastic models, in which the identity of the overrepresented strains infrequently changes [18]. For two diallelic loci, epistasis is weak if: *σ*_*𝒮*_ = 0 and hosts are likely to be infected multiple times (*R*_0_ is large); *σ*_*𝒮*_ *≈* 0 and *σ*_*𝒜*_ *≈* 0; and/or hosts are likely to be infected once and *σ*_*𝒮*_ *≈ σ*_*𝒜*_.

Second, demographic stochasticity causes the loss of strains or antigens by chance [21, 23, 50, 54, 67, 68]. This is especially relevant if the segregating loci have an unequal number of alleles. Here, antigens at the more diverse locus are predicted to be more easily lost in finite populations, particularly when cross-protection is high [23]. This can be explained as follows (see also Sup. Info. S8). If one locus is more antigenically diverse than the others, e.g., if two loci have alleles *{a, A}* and *{B*_1_, *B*_2_, *B*_3_*}*, it is impossible to maintain each antigen at its expected frequency, *p*_*A*_(*t*) = 1*/*2 and *p*_*Bi*_ (*t*) = 1*/*3, and simultaneously maximize LD between each *{A, B*_*i*_*}*. This creates tension between selection on antigens through the additive selection coefficients and selection on strains through epistasis in fitness. This tension is modulated by the relative strength of *σ*_*𝒜*_ and *σ*_*𝒮*_. As cross-protection increases, selection maintaining antigens at their expected frequencies becomes weaker (i.e., the strength of *σ*_*𝒮*_ decreases). At the same time, epistasis becomes stronger, favouring some strains (i.e., *σ*_*𝒜*_ becomes increasingly positive). Consequently, some antigens at the more diverse loci are forced to lower frequencies and are thus more likely to be stochastically lost.

### The multilocus perspective clarifies the source of PSS and the drivers of antigenic variation

The multilocus perspective identifies epistasis as the source of PSS, which explains the previously contentious role played by negative frequency-dependent selection (NFDS) for PSS[18–20, 69]. Although NFDS is required to maintain genetic polymorphism, it need not yield epistasis (e.g., if *σ*_*𝒜*_ = *σ*_*𝒮*_ = 0). Our multilocus perspective demonstrates that epistasis is necessary to produce strain overrepresentation. Indeed, past work has compared time series of the genomic patterns produced by models of PSS against a null model in which antigenic variation is neutral [19, 20]. The multilocus perspective suggests an intermediate null model, in which there is NFDS but no epistasis (e.g., *σ*_*𝒜*_ = 0, *σ*_*𝒮*_ = 0). In such a model, NFDS would maintain antigenic variation but no directional force would maintain distinct allelic combinations, such that demographic stochasticity would be the only cause of LD. Moreover, the multilocus perspective synthesizes different explanations for observed patterns of antigenic diversity, summarized in a recent review [8]. In addition to host heterogeneity and functional constraints, the authors discussed three hypotheses to explain patterns of antigenic diversity. First, each antigen contributes to host immunity additively (so *σ*_*𝒜*_ = 0); second, immunity to a single antigen provides strong cross-protection [i.e., *σ*_1_ = *σ*_*i*+1_, *i >* 0; 6, 12]; and third, there is an interaction between antigens at different loci. The multilocus perspective reveals considerable overlap between these hypotheses; for example, although the first and second predict, respectively, no PSS and stable pairwise LD (DSS), the third simply means that there is epistasis (e.g., *σ*_*A*_*≠* 0), which can lead to either no PSS or stable PSS, as well as PSS consisting of higher-order LD. Indeed, as the number of antigens required to induce a strongly protective immune response determines the probable order of LD, the likelihood a novel antigen can evade population immunity increases with the dominant order of LD. Thus, we would predict a positive correlation between the dominant order of LD and the amount of allelic variation at each locus.

### Multilocus models and vaccination

Our analysis provides new insight into ‘evolutionarily optimal’ vaccine design, complementing a body of work trying to understand how genetically diverse pathogen populations respond to vaccination [70–73]. Our work shows that the dominant order LD is generally indicative of how many antigens are required to generate a strongly protective immune response. However, our analysis also emphasizes challenges associated with predicting the consequences of vaccination. First, the input of vaccinal immunity to the population may disrupt the relationship between strain frequency and population immunity with complex dynamical consequences [e.g., 10, 11, 74, 75]. Second, the predicted optimal vaccine type depends on whether homologous immunity is more protective than full antigenic immunity (*σ*_*𝒮*_ *>* 0), and if so, whether vaccination provides equivalent protection to full antigenic or homologous immunity. Third, the pathogen population response will hinge upon whether evolutionary bistability is possible, as the influx of vaccinal immunity may alter the dominant order of PSS, increasing or decreasing genotypic diversity. Fourth, vaccines are often chosen to target a subset of strains, such as those linked to virulence factors [76]. The inclusion of additional pathogen life-history variation, such as virulence factors, will induce fitness asymmetries among strains that are difficult to address with existing models. However, by emphasizing key evolutionary quantities, the multilocus perspective provides a useful frame of reference to tackle these questions in the future.

## Conclusion

Many key human pathogen species, including malaria, dengue, *S. pneumoniae, N. meningitidis* and cytomegalovirus, consist of multiple coexisting strains that make medical interventions, such as vaccination, challenging. Thus, understanding the dynamics of multistrain pathogens is important not only to unravel what maintains and structures genetic diversity, but as an urgent question of public health. Yet, combining models of pathogen genetic diversity with realistic epidemiological feedbacks has remained a daunting task. One way forward is by adopting a population genetics perspective, and focusing upon evolutionary quantities like linkage disequilibrium. As we have shown here, taking such an approach can disentangle the factors driving the evolutionary dynamics of pathogen populations. It illustrates when, and why, pathogen strain structure occurs, and carries the promise to predict pathogen dynamics and identify pathogen-specific optimal interventions.

## Acknowledgements

SG thanks Julia Gog and Sunetra Gupta for many inspiring discussions.

## Supplementary Information

### S1 Epidemiological model

#### S1.1 Basic model without adaptive immunity

In the absence of immunity, the host-pathogen model considered here is straightforward: the pathogen is transmitted to uninfected hosts by mass-action with rate constant *β*. If transmission occurs, an infection is successfully established with probability *σ*, while infections are naturally cleared by the host at a per-capita rate *µ*. Hosts enter the population at a fixed rate *b* and are removed from the population due to natural- and virulence-related mortality (if infected) at per-capita rates *d* and *α*, respectively. Hosts cannot be simultaneously infected by multiple strains (no co-infection); we later discuss how co-infection affects the dynamics. At time *t*, let *S*(*t*) denote the density of uninfected hosts and and *I*(*t*) denote the total density of infected hosts. Then, the epidemiological dynamics are

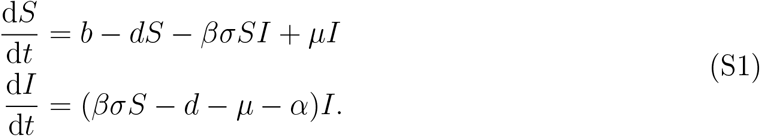

The equilibrium density of uninfected hosts in the absence of the pathogen is *b/d*. Therefore, the basic reproduction number of the pathogen, *R*_0_, is

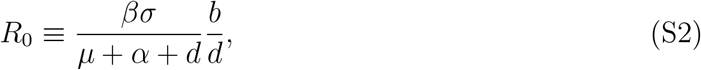

and the endemic equilibrium is

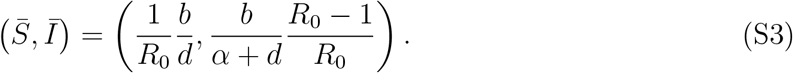

The endemic equilibrium is globally asymptotically stable whenever *R*_0_ *>* 1. We denote the ratio between the expected duration of infection, 1*/*(*µ* + *α* + *d*), and the expected lifespan of the host, 1*/d*, as

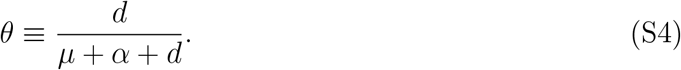

For small *θ*, infections are acute, whereas as *θ →* 1, infections become increasingly chronic.

##### S1.1.1 ‘Length’ of immune history

Both *R*_0_ and *θ* drive the dynamics of the pathogen and affect t he ‘ length’ o f t he immune history of an individual host. In particular, they determine (i) the expected number of infections a host experiences through its lifespan, and (ii) the number of prior infections of a randomly selected uninfected host.

To calculate the expected number of infections a host experiences through its lifespan, suppose we are at the endemic equilibrium, that *b* = *d*, and, for simplicity, *α* = 0. The probability, *P*_*n*_, that a host is infected *n >* 0 times during its life, is the product of the probability that a host is infected, and clears, *n −* 1 infections

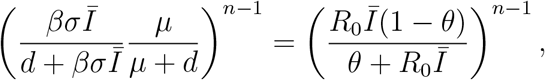

and the probability a host is infected an *n*-th time, after which the host dies before becoming re-infected:

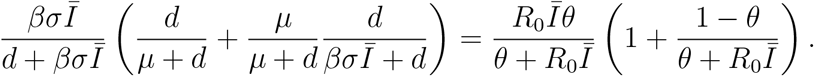

The expected number of infections during the lifespan of an average host, which we denote *T*, is

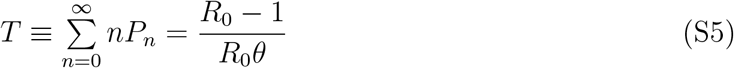

where to compute the sum we have used the relation 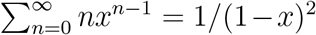 for 0 *< x <* 1. The expected number of prior infections of a randomly selected uninfected host also depends upon *R*_0_ and *θ*. To calculate this, let 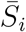 denote the equilibrium density of uninfected hosts with *i* prior infections, and 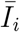 the density of infected hosts with a history of *i* infections (including the current infection). Assuming *α* = 0 and *b* = *d* as before, at the endemic equilibrium

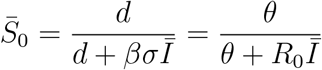

and

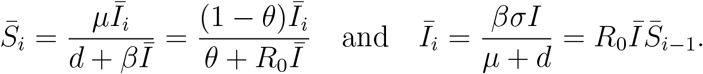

Therefore,

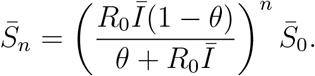

Hence, the expected number of prior infections of a randomly selected uninfected host, which we denote *M*, is

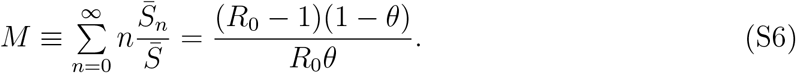

Notice that both *M* and *T* increase with *R*_0_ and decrease with *θ* (i.e., infections become increasingly chronic). Although these calculations do not account for immune protections, immunity will generally act to reduce *M* and *T*, and not change their qualitative dependence upon *R*_0_ and *θ*. Thus, the ‘length’ of immune history, which is discussed in the main text, is expected to depend on *M* and *T*, and therefore to increase with *R*_0_ and decrease with *θ*.

#### S1.2 Specification of adaptive immunity

Adaptive immunity is built into the basic model of equations (S1) as follows. We consider a pathogen whose genotype consists of some number of multiallelic loci. The host can develop an adaptive immune response to the alleles (i.e., antigens) at each of these loci. This immune response can potentially reduce:

i. The susceptibility to infection; we let 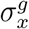 denote the probability of infection of a host with immune history *x* by strain (genotype) *g*.
ii. The transmissibility of an infection; we let 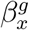 denote the transmissibility of an infection by strain *g* from a host with immune history *x*.
iii. The duration of infection; we let 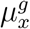 denote the per-capita rate at which a host with immune history *x* clears an infection by strain *g*.
iv. The virulence-related mortality; we let 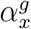 denote the per-capita rate at which a host with immune history *x* dies due to an infection by strain *g*.

For each of these protections, we distinguish between immunity to antigens and immunity to previously-infecting strains (i.e., homologous immunity). In particular, for a host with immunity to *n* = 0, 1, 2, … antigens of strain *g*, the immune protections are

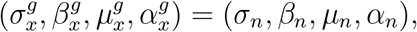

whereas for a host with homologous immunity to strain *g*, the immune protections are

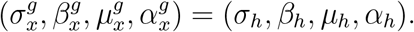

We assume that homologous immunity is at least as strong as full antigenic immunity, and that the host immune response is always the most protective available to the host, based upon its immune history. For example, for a pathogen genome with two diallelic loci, *{a, A}* and *{b, B}*, a host with immune history *x*^*′*^ = *{ab, Ab}* has immune protections

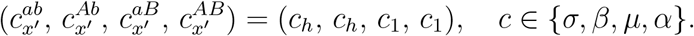

#### S1.3 Epidemiological dynamics and per-capita growth rates

At time *t*, let *S*_*x*_(*t*) denote the density of uninfected hosts with prior infection by strains belonging to the set *x* (so *x* is the immune history), and let 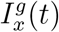 denote the density of strain *g* infections in hosts with immune history *x*. The resulting epidemiological dynamics are

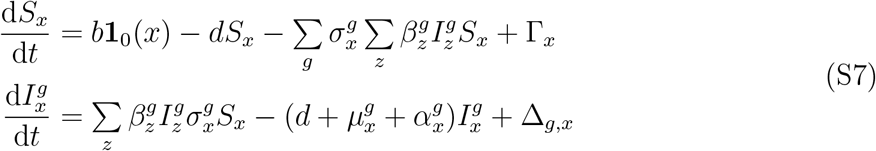

where **1**_0_(*x*) = 1 if *x* = 0 and 0 otherwise, Δ_*g,x*_ is the net change due to recombination of strain *g* infections in hosts with immune history *x* (for now, left arbitrary), and Γ_*x*_ controls the rate at which hosts gain (or lose) the immune history *x*. If immunity is life-long and is updated when infections are cleared,

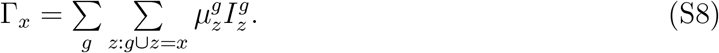

Next, let *r*_*g*_ denote the per-capita growth rate of strain *g*; this is the growth rate of strain *g* averaged across the different immune environments *x*. It follows that

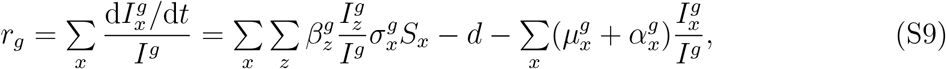

where 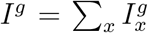 is the total density of strain *g*. In Box 1 of the main text we present a version of this model where we have restricted immune protections to reducing 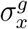 That is, immunity only reduces host susceptibility to infection.

### S2 Biological connections between existing models

Next, we discuss the relationship between model (S7) and existing models. Multistrain models typically differ from one another with respect to three biological properties:

i. the immune protections, and how these are affected by antigenic distance;
ii. the specification of how hosts acquire immunity, that is, the specification of Γ_*x*_; and
iii. whether co-infection is permitted or not.

Here, we discuss each of these assumptions in more detail, and in Table S2 we summarize how the model (S7) relates to existing models. It is important to note that the combination of assumptions employed by previous models is typically made because they reduce the number of equations. Doing so makes the problem more tractable for mathematical analysis and simulation. As our purpose here is to understand and, where possible, generalize the biological predictions of these models, we will simply state the assumptions made by each model without showing the mathematical benefits of these assumptions. We refer interested readers to Kucharski & Gog [30], who provide a detailed treatment of the mathematics behind some of the commonly used models.

#### S2.1 Immune protections

Multistrain models vary in what type of protection immunity provides and how prior infections combine to determine the strength of immunity.

First, most models assume that immunity reduces either transmissibility or susceptibility; this choice is made typically for model-specific mathematical reasons. For example, ‘status’-based models often assume reduced susceptibility [13, 27, 34], whereas ‘history’-based models often assume reduced transmissibility [6, 9, 12, 16] (but not always, [14, 42]). More recently, simulation models have assumed that immune protections reduce the duration of carriage, as this is more biologically realistic for malaria [18–20]. Finally, some models of antibody-dependent enhancement have allowed immunity to reduce virulence-related mortality [47, 48]. Although immune protections reducing virulence-related mortality can have a qualitatively different effect [e.g., 47], it is generally believed that there is no qualitative difference between protections reducing susceptibility, transmissibility, or clearance [15, 26]. At least one consequence of the differences in immune protections is that if immunity reduces susceptibility, hosts are less likely to acquire immunity to different strains, provided that immunity is only acquired after successful infection [15].

Second, models differ in how multiple prior infections combine to determine the strength of immunity. We will illustrate these differences using the example immune history *x*^*′′*^ = *{Ab, aB}*. Thus, we focus on a pathogen genome consisting of two diallelic loci. Moreover, we focus on the case in which immune protections reduce susceptibility to infection; the other cases follow similarly. The most common approach is to assume that immunity against a particular strain is determined by the *single* most antigenically similar previously infecting strain [6, 9, 12, 16, 25]. For the immune history *x*^*′′*^, we obtain

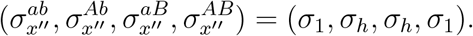

A less common model assumption is that immunity is the *product* of the individual cross-protections against previously infecting strains [14, 24]. For the immune history *x*^*′′*^, this yields

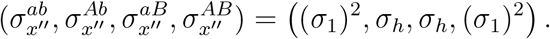

Finally, immunity to each antigen can be treated independently. Here, the strength of immunity is *antigen-specific* and depends upon the total number of antigens against which the host has protection [18–20]. For the immune history *x*^*′′*^, it follows that

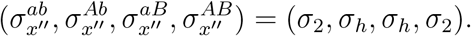

Note that although some models restrict immune memory to only the most recent infections [67, 77], we do not consider this further here, as it can be considered a form of waning immunity.

#### S2.2 Acquisition of immunity

The model of (S7) assumes that after hosts have naturally cleared an infection, they have homologous immunity to the infecting strain, as well as some degree of protection against antigenically similar strains. Such a framework has been referred to as a ‘history-based’ model [13, 34], and is a commonly taken modeling approach [6, 12, 14, 42]. An alternative to history-based models are so-called ‘status-based’ models [13, 27, 34]. Status-based models assume that each host is either completely protected or completely susceptible to a given strain (so immunity reduces susceptibility); there is no ‘partial’ protection at the level of the individual host. Thus, following clearance of an infection (or at time of infection, depending upon when immunity is acquired), each host becomes either fully-protected or fully-sensitive to each strain. The probability of acquiring protection against a particular strain then depends upon the antigenic similarity between that strain and the other strains against which the host has protection [13, 27, 34].

To convert model (S7) into a status-based model, we first restrict immune protections to reducing susceptibility, such that

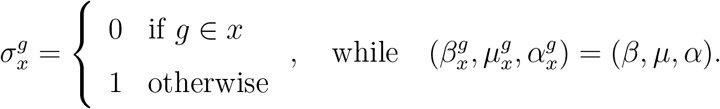

Then, we suppose that upon clearing an infection by strain *g*, a host with immune history *z* becomes a host with immune history *x* with probability 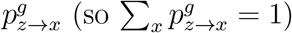, and thus

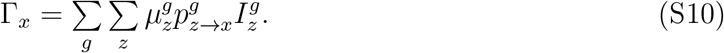

Under these assumptions, model (S7) is a status-based model.

#### S2.3 Co-infection

The model of (S7) assumes that transmission only occurs to uninfected hosts, and thus hosts cannot be simultaneously infected by different strains (i.e., there is no co-infection). This assumption was made primarily for modelling clarity. It is biologically reasonable if, for example, the pathogen is circulating at a low prevalence, infections are acute, and/or infection induces short-term strain-transcending immunity. Yet, these conditions need not apply generally. Hence, co-infection is a common attribute of multistrain models. Most analytically tractable models that include co-infection make all of the following assumptions [6, 12, 15, 41, 54]:

i. Hosts can be simultaneously infected by any number of *different* strains and these strains do not interact. Thus, transmission and clearance rates are unaffected by the presence, or number, of co-infecting strains. Here, the immune environment a strain experiences is set at the time of infection and is unaffected by subsequent infections.
ii. There is no virulence mortality 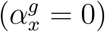 to ensure independence of co-infecting strains [61, 62].
iii. Homologous immunity is sterilizing and life-long (*σ*_*h*_ = 0).

Incorporating these assumptions into our model alters the specification of Γ_*x*_ and the biological interpretation of *S*_*x*_(*t*) in equation (S7). In particular, *S*_*x*_(*t*) now denotes the density of hosts with immune history *x*, irrespective of their current infection status, and

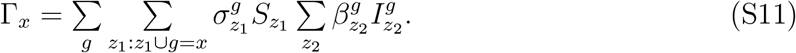

Importantly, because homologous immunity is sterilizing (i.e., *σ*_*h*_ = 0), and acquired at the time of infection, a host that is infected by strain *g*^*′*^ cannot be reinfected by strain *g*^*′*^. As a consequence, and because co-infections are assumed to be non-interacting, it is not necessary to keep track of how many (and which) strains currently infect a given host.

Notably, treating co-infecting strains as non-interacting is biologically unsatisfying, given that a host simultaneously infected with *n* strains is *n*-fold as infectious as a host infected with one strain. Specifically, genetic diversity is unreasonably easily maintained under this assumption. For example, Buckee et al [22] studied the evolution of hypervirulent lineages of *Neisseria meningitidis* using a model in which each strain is defined by its sequence type and level of virulence (which causes host mortality). Since hypervirulence shortens the duration of infection, it should be eliminated from the population by selection. But as hosts can be simultaneously infected by any number of genetically distinct strains, each level of virulence creates ‘new’ genetically distinct strains that can therefore co-infect the same host (even if they share the same sequence type as existing infecting strains), thus maintaining variation in virulence. Incorporating virulence in models of co-infection is not trivial and requires co-infecting strains to interact [61, 62].

### S3 Separation of evolutionary and epidemiological dynamics

The number of equations associated with model (S7) makes the full model challenging to analyze and to interpret. Thus, a variety of clever mathematical approximations have been employed to simplify the dynamics of multistrain models [e.g., 6, 13, 14, 42, 61, 78]. Although these techniques are effective at reducing the (mathematical) dimensionality of the problem, they struggle to provide an evolutionary context for the predictions; for example, they do not provide biological intuition why oscillations are possible in some models but not others [see discussion in 27]. Thus, many important results lack a clear biological interpretation. A promising way to provide greater clarity on the evolutionary process is to separate the variables describing the epidemiological dynamics from those describing the evolutionary dynamics [79, 80]. Doing so allows for the identification and interpretation of the evolutionary forces at play. The evolutionary variables are the allele (i.e., antigen) frequencies and the linkage disequilibrium (LD; i.e., nonrandom associations between alleles) [31]. The epidemiological variables include the total density of infections, *I*(*t*), and the population distribution of immune histories across the uninfected hosts, *S*_*x*_(*t*). Here, we focus on extracting the evolutionary variables for a pathogen genome consisting of two diallelic loci; in Sections S8 and S9 we extend our model and analysis to the study of greater antigenic variation.

**Table S2:**
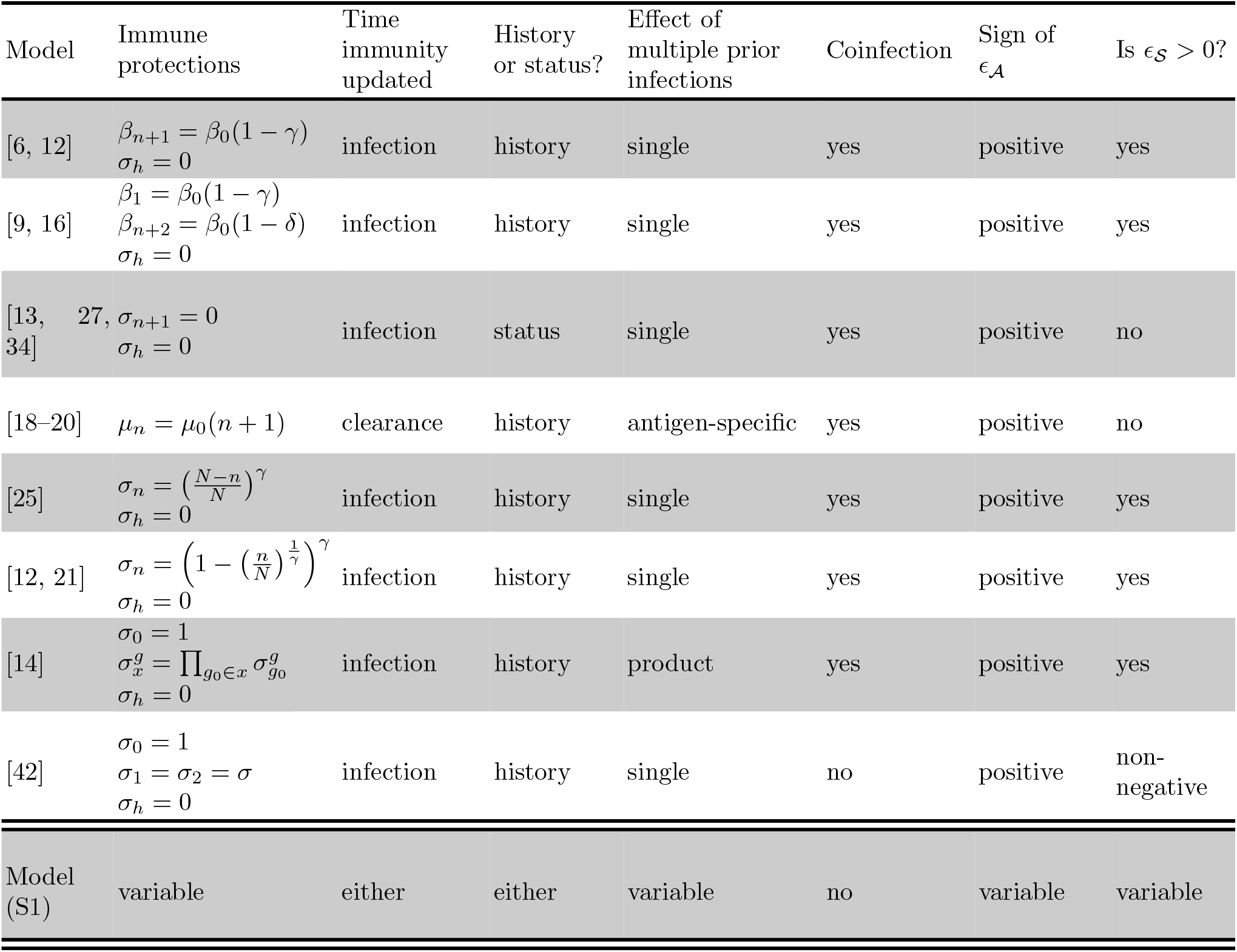
Summary of key models and their relationship to the framework used here. Here, *γ, σ*, and *δ* are positive parameters and *N* is the total number of loci. We define *ϵ*_𝒜_ *≡ f*_0_ *−* 2*f*_1_ + *f*_2_ where *f*_*i*_ *≡ σ*_*i*_*β*_*i*_*/*(*μ*_*i*_ + *α*_*i*_ + *d*), and *ϵ*_𝒮_ *≡ f*_2_ *− f*_h_. Note that both *ϵ*_𝒜_ and *ϵ*_𝒮_ are calculated assuming there are only two diallelic loci. In this case if *σ*_h_ = 0, each of the different ways in which immunity to multiple prior strains combines (single, antigen-specific, and product) is captured by model (S7). There also exist various models whose specification of cross-immunity assumes a one-dimensional antigenic space [e.g., 24, 42], which we do not consider here (the model of [42] considered in the table is the version with four strains).

#### S3.1 The additive selection coefficients and epistasis in fitness

For two diallelic loci, there are four possible pathogen genotypes (strains),

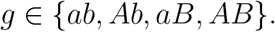

Without loss of generality, we consider strain *ab* to be the reference genotype. Then, the additive selection coefficients of alleles *A* and *B* are

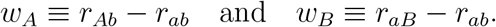

The selection coefficients indicate whether allele *A* (r esp. *B*) is favoured, i. e., *w*_*A*_ *>* 0 (resp. *w*_*B*_ *>* 0), or disfavoured, i.e., *w*_*A*_ *<* 0 (resp. *w*_*B*_ *<* 0), over allele *a*, when *A* (resp. *B*) occurs on the genetic background *b* (resp. *a*). We also account for whether the fitness o f an allele depends upon its genetic background, i.e., whether there is *epistasis in fitness*. Epistasis in fitness is defined as

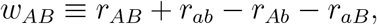

and indicates whether the combination of alleles *A* and *B* is favoured (*w*_*AB*_ *>* 0) or dis-favoured (*w*_*AB*_ *<* 0) over what is expected based upon the additive selection coefficients. Thus, the additive selection coefficients can be thought of as controlling selection on individual alleles, whereas epistasis can be thought of as controlling selection on particular combinations of alleles (in the case of two loci, these combinations are equivalent to genotypes or strains).

Although we can write down the additive selection coefficients and epistasis for the general model, here we assume that immune protections only affect susceptibility. That is because if immune protections only affect susceptibility, the per-capita growth rates of equation (S9) can be written in a particularly simple and illustrative form. In Section S5, we consider other immune protections using an equilibrium analysis.

To write down *w*_*A*_, *w*_*B*_, and *w*_*AB*_, we define

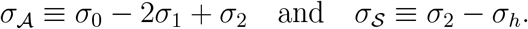

Conceptually, if we consider the function mapping antigenic immunity to the strength of immune protections, *h* : *n* ↦ *σ*_*n*_, then *σ*_𝒜_ is the central difference approximation of d^2^*h/*d*n*^2^, evaluated at *n* = 1. Consequently, *σ*_𝒜_ can be thought of as controlling whether the function relating antigen-specific cross-protection to the number of antigens recognized by the immune system has an accelerating or saturating shape. The function has an accelerating shape if *σ*_𝒜_ *<* 0, and a saturating shape if *σ*_𝒜_ *>* 0 (Fig. 2). On the other hand, *σ*_𝒮_ measures how much more protective homologous immunity is than full antigenic immunity. If *σ*_𝒮_ = 0, full antigenic immunity is equally strong as homologous immunity.

Using these composite parameters, the per-capita growth rate can be written as

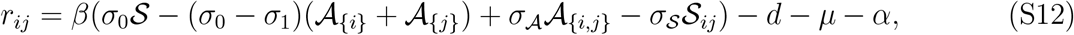

where 𝒜_*{i}*_(*t*) and 𝒜_*{i,j}*_(*t*) are the densities of uninfected hosts that have immunity to *at least* antigen *i* or antigens *i* and *j*, *S*(*t*) = ∑_*x*_ 𝒮_*x*_(*t*) is the total density of uninfected hosts, and 𝒮_*ij*_(*t*) is the density of uninfected hosts that have homologous immunity (prior infection) to *at least* strain *ij*.

Then, the additive selection coefficients can be written as

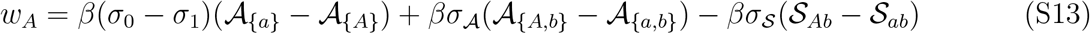

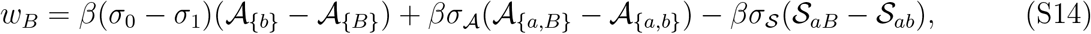

and epistasis in fitness is

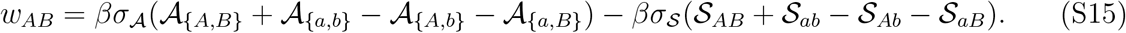

#### S3.2 Relationship between 𝒜_Δ_(*t*) and 𝒮_Δ_(*t*)

Define

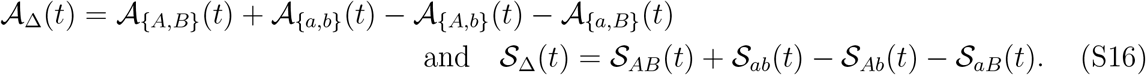

The sign of 𝒜_Δ_(*t*) indicates whether more uninfected hosts have full antigenic immunity to strains belonging to the set *{ab, AB}* (𝒜_Δ_(*t*) *>* 0) or to the set *{Ab, aB}* (𝒜_Δ_(*t*) *<* 0). The sign of 𝒮_Δ_(*t*) indicates whether more uninfected hosts have homologous immunity to strains belonging to the set *{ab, AB}* (𝒮_Δ_(*t*) *>* 0) or to the set *{Ab, aB}* (𝒮_Δ_(*t*) *<* 0). Using these definitions, we can rewrite epistasis in fitness (equation (S15)) as

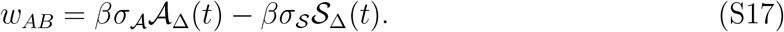

Next, notice that if we expand the sums in 𝒜_Δ_(*t*), we obtain after cancellation

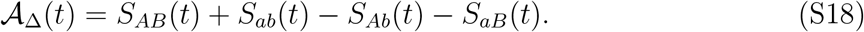

Thus, only hosts with immunity to the antigens defining a single strain contribute to 𝒜_Δ_(*t*). Unlike 𝒜_Δ_(*t*), however, 𝒮_Δ_(*t*) depends upon hosts with immunity to more than one strain, since, after cancellation,

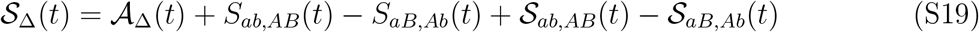

where *S*_*ij,kℓ*_(*t*) denotes the density of uninfected hosts that have homologous immunity to *only* strains *ij* and *kℓ*, and *S*_*ij,kℓ*_(*t*) denotes the density of uninfected hosts that have homologous immunity to *at least* strains *ij* and *kℓ*. A graphical representation of this relationship is provided in Figure S1. As discussed in the main text, in order for 𝒮_Δ_(*t*) and 𝒜_Δ_(*t*) to diverge, hosts must be infected multiple times during their lifespan (and retain memory of these infections).

**Figure S1:**
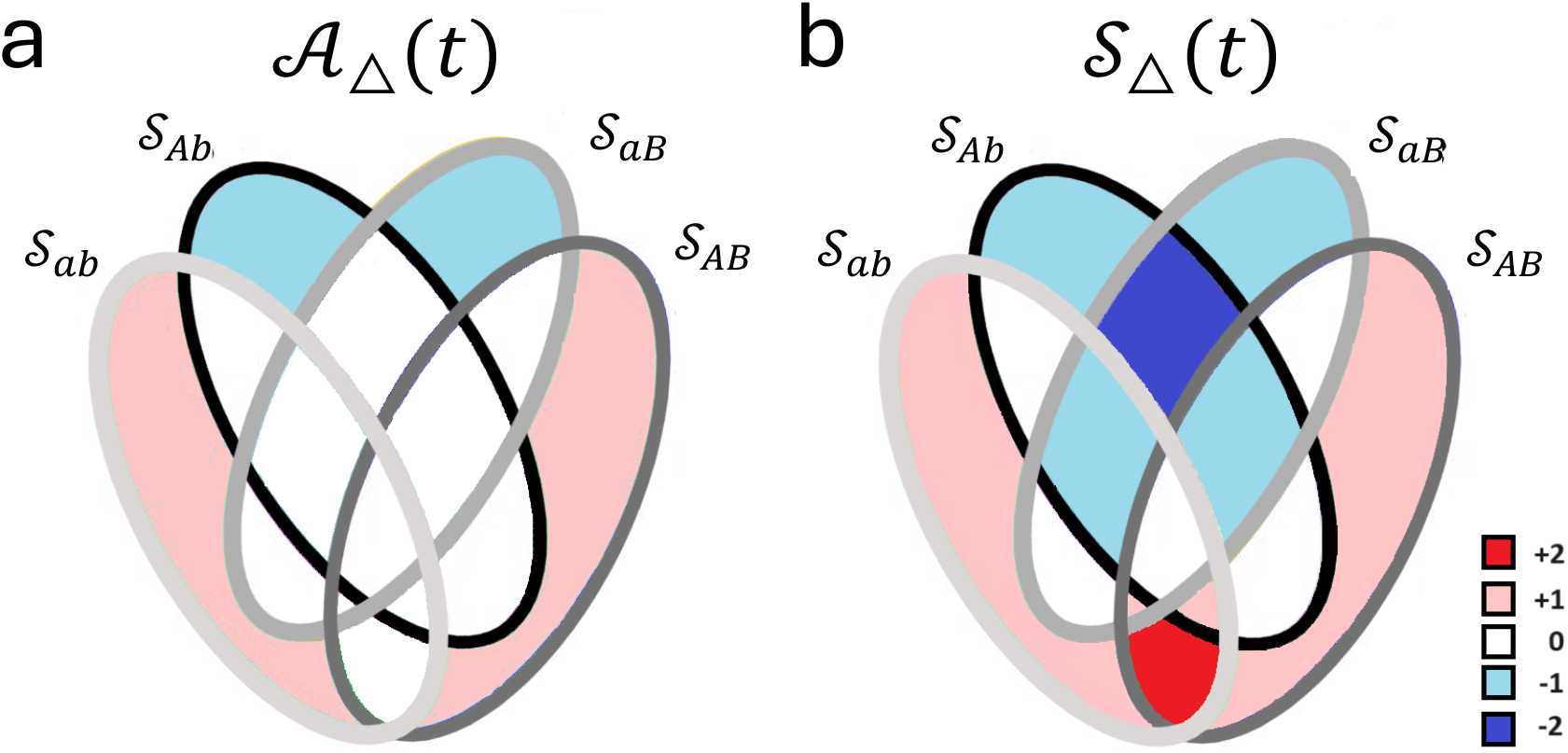
The dynamic immunological landscape. Panel **a** is a Venn diagram of 𝒜_Δ_(*t*), that is, whether more uninfected hosts have full antigenic immunity to strains belonging to the set *{ab, AB}* (i.e., 𝒜_Δ_(*t*) > 0) or to the set *{aB, Ab}* (i.e., 𝒜_Δ_(*t*) < 0). Panel **b** is a Venn diagram of 𝒮_Δ_(*t*), that is, whether more uninfected hosts have homologous immunity to strains belonging to the set *{ab, AB}* (𝒮_Δ_(*t*) > 0) or to the set *{aB, Ab}* (𝒮_Δ_(*t*) < 0). Each ellipse corresponds to a different density of hosts with homologous immunity to *at least* strain *ij*, 𝒮_ij_(*t*). Then, 𝒜_Δ_(*t*) and 𝒮_Δ_(*t*) are computed as the weighted sum of the coloured regions, where the colors indicate the weights. Whereas 𝒜_Δ_(*t*) only includes uninfected hosts with homologous immunity to a single strain, 𝒮_Δ_(*t*) includes uninfected hosts with homologous immunity to multiple strains.

#### S3.3 Equations describing the evolutionary dynamics

To write out the evolutionary dynamics, we first define the evolutionary variables. The allele frequencies are

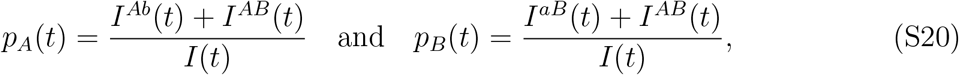

and the linkage disequilibrium between alleles *A* and *B* is

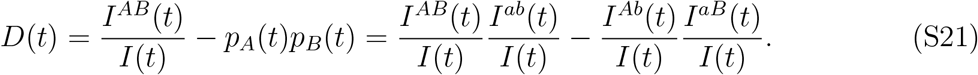

Then, the evolutionary dynamics can be written as

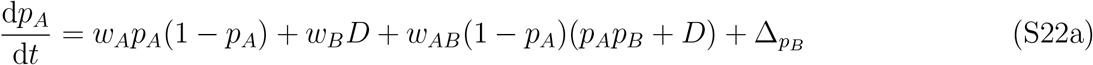

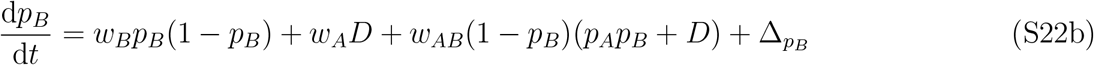

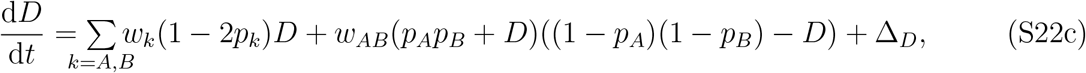

where Δ_*q*_ indicates the effect of recombination on the variable *q*. Note that the general form of these equations follows standard population genetics and is not particular to the epidemiological system we consider here. Epidemiology enters this system through the additive selection coefficients, recombination, and epistasis in fitness, which can vary through time due to the epidemiological dynamics. By writing out the evolutionary dynamics as we have done in (S22), we can identify the evolutionary forces at play and study how they impact the evolutionary process. We can then link each of these forces to the epidemiological model of interest.

#### S3.4 Fast and slow timescales in the evolutionary dynamics

Unbiased recombination does not affect antigen frequencies, Δ_*p B*_ = Δ_*pA*_ = 0 and acts only to remove LD from the population. That is, Δ_*D*_ *≈ −ρD*(*t*), where 0 *< ρ* ≪ 1 is a positive (but small) constant controlling the rate of recombination. Because we assume that there are no intrinsic biases favouring one antigen over the other at a given locus, and because immunity induces strong negative frequency-dependent selection (NFDS) at each locus, the antigen frequencies will quickly reach a quasi-equilibrium state at which each antigen is present at equal frequency within the population, that is, *p*_*k*_(*t*) → 1*/*2 for *k* ∈ *{a, A, b, B}*. Once this quasi-equilibrium state is reached, since *p*_*A*_(*t*) = *p*_*B*_(*t*) = *p*_*b*_(*t*) = *p*_*a*_(*t*), we also obtain *p*_*Ab*_(*t*) = *p*_*aB*_(*t*) and *p*_*AB*_(*t*) = *p*_*ab*_(*t*). That is, the inherent symmetry within the system drives the population to self-organize into two sets of strains, *{ab, AB}* and *{Ab, aB}*. At this point, the members of each set behave identically dynamically. Consequently, strains within each set have the same per-capita growth rates, that is, *r*_*ab*_ = *r*_*AB*_ and *r*_*Ab*_ = *r*_*aB*_. Thus the evolutionary dynamics occur on two timescales. On the fast timescale, strong NFDS operating through the additive selection coefficients will push the antigen and strain frequencies to a quasi-equilibrium state, *p*_*A*_(*t*) *≈* 1*/*2, *p*_*B*_(*t*) *≈* 1*/*2, *p*_*AB*_(*t*) *≈ p*_*ab*_(*t*), and *p*_*Ab*_(*t*) *≈ p*_*aB*_(*t*). At this quasi-equilibrium state, since *r*_*AB*_ = *r*_*ab*_ and *r*_*Ab*_ = *r*_*aB*_, we have

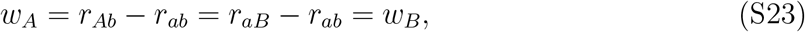

and moreover,

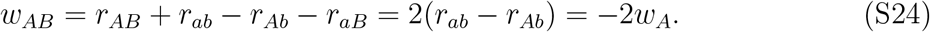

Therefore at quasi-equilibrium we have the relations *w*_*A*_ *≈ w*_*B*_ *≈ −w*_*AB*_*/*2, and so on the slow timescale, the system of equations (S22) reduces to the single equation

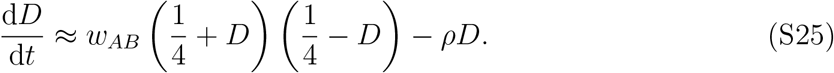

Because LD quantifies whether certain strains are more, or less, abundant than predicted based upon allele frequencies, the presence of LD in the population is necessary for the existence of pathogen strain structure or PSS (see the longer discussion in Section 2 of the main text). Thus, on the slow timescale, equation (S25) captures the evolutionary dynamics of PSS. Moreover, since *p*_*AB*_(*t*) = *p*_*ab*_(*t*) and *p*_*Ab*_(*t*) = *p*_*aB*_(*t*), if *D*(*t*) *>* 0, then the set of strains *{ab, AB}* will be overrepresented in the population, whereas if *D*(*t*) *<* 0, the set of strains *{Ab, aB}* will be overrepresented. Note that in this particular case, an overrepresented strain is also more abundant.

Assuming *ρ* = 0, equation (S25) has four potential equilibria: (1,2) when either set of strains has competitively excluded the other (*D*(*t*) = *±*1*/*4); (3) when one set of strains is overrepresented, but all strains are present (*w*_*AB*_ = 0 for 0 *<* |*D*| *<* 1*/*4); and (4) when there is no structure (*w*_*AB*_ = 0 for *D* = 0). Finally, if the sign of epistasis fluctuates, evolutionary oscillations, here defined to be any sustained dynamical behaviour of LD (including cycles and chaos) may occur. In all cases, however, the dynamical behaviour of equation (S25) hinges upon epistasis. The asymptotic behaviour of LD, arising from numerical simulation of the full epidemiological model of Box 1 is illustrated in Figure 3, and Figure S2 shows representative examples of the dynamical behaviour.

**Figure S2:**
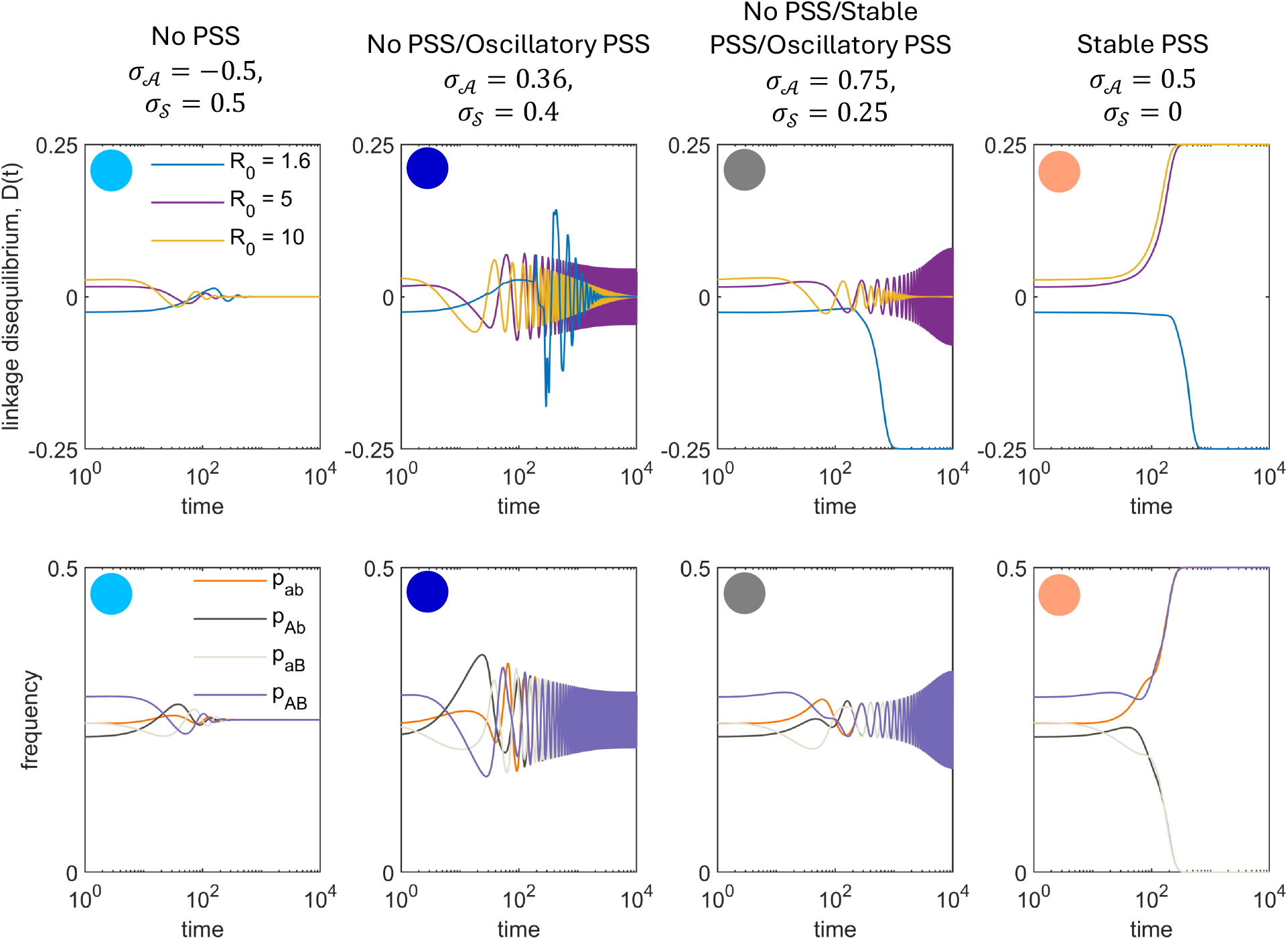
Dynamical behaviour of LD and strain frequencies as *σ*_𝒜_, *σ*_𝒮_, *θ* and *R*_0_ vary. The dynamics of LD depend upon the values of *σ*_𝒜_ and *σ*_𝒮_, and also on the dynamic immunological landscape (controlled by *R*_0_ and *θ*). Each column corresponds to the indicated combination of *σ*_𝒜_ and *σ*_𝒮_. Coloured circles indicate the approximate position of the shown scenario in Figure 3. In the first row, the different trajectories correspond to the indicated value of *R*_0_. The second row shows the strain frequency dynamics for a fixed value of *R*_0_ = 4. The second column assumes *θ* = 0.01, whereas the other columns assume *θ* = 0.05. All simulations assume *b* = *d* = 0.01, *α* = 0, *ρ* = 10^−8^, *σ*_0_ = 1 and *σ*_h_ = 0, whereas *σ*_1_ and *σ*_2_ vary by panel.

Finally, we note that given an observed distribution of allele frequencies, there is a maximum and minimum value of LD. For example, for two diallelic loci

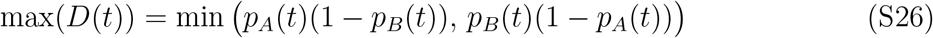

and

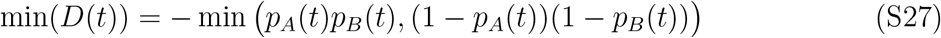

[see 31]. Importantly, in order for pairwise LD to achieve this maximum or minimum, at least one strain must be absent from the population; in the special case when *p*_*A*_(*t*) = *p*_*B*_(*t*), two strains are absent. Similar results hold for higher-order LD among more than two loci [81, 82], and when LD reaches either its maximum or minimum, one or more strains are absent from the population. We will use *D*^*^ to indicate that the specified order of LD attains either its maximum or minimum, that is, one or more strains are competitively excluded.

### S4 Evolutionary dynamics on the slow timescale

In this section, we expand on our discussion in the main text about the evolutionary dynamics, focusing in particular on the role of *σ*_𝒜_ and *σ*_𝒮_. We do so in three sections. First, we attempt to provide additional intuition for our predictions. Second, we show the necessary conditions for stable PSS when *σ*_𝒮_ = 0 is *σ*_𝒜_ *>* 0. Finally, we provide numerical results supporting the claim that when *σ*_𝒮_ = 0, evolutionary oscillations do not occur and the long-term outcome is determined by the sign of *σ*_𝒜_.

#### S4.1 Intuition behind the dynamics

First suppose *σ*_𝒮_ = 0, that is, homologous immunity is no more protective than full antigenic immunity. In this case,

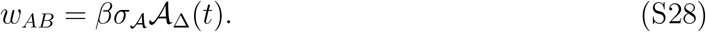

As was demonstrated in Section S3.2, we can write

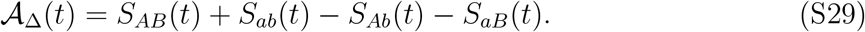

In the absence of waning immunity, 𝒜_Δ_(*t*) indicates whether more uninfected hosts with a single prior infection were infected by one of the strains belonging to one set of strains or the other, i.e., *{ab, AB}* or *{Ab, aB}*. If the set of strains *{ab, AB}* was responsible for more of these infections, 𝒜_Δ_(*t*) *>* 0; if the set of strains *{Ab, aB}* was responsible for more of these infections, 𝒜_Δ_(*t*) *<* 0. Because there is a single epidemiological feedback in (S28), 𝒜_Δ_(*t*), we would not expect epistasis to undergo sustained oscillations, nor for it to change sign at a particular amount of LD. Therefore, we eventually expect LD to reach an equilibrium: either stable PSS, |*D*(*t*)| → 1*/*4, or no PSS, *D*(*t*) → 0. The sign of *σ*_𝒜_ determines which of these outcomes occurs: if *σ*_𝒜_ *>* 0, then stable PSS occurs, whereas if *σ*_𝒜_ *<* 0, all strains coexist at equal frequencies. Finally, if *σ*_𝒜_ = 0, then epistasis is identically zero. Here, the equilibrium amount of LD is contingent on the initial conditions. Note that this prediction for *σ*_𝒜_ = 0 assumes *ρ* = 0; otherwise, for any non-zero *ρ*, if *σ*_𝒜_ = 0, then no PSS is the only stable state. Below we provide the intuition for these predictions.

##### S4.1.1 Sign of *σ*_𝒜_

Suppose we are on the slow timescale, such that *p*_*AB*_(*t*) = *p*_*ab*_(*t*) and *p*_*Ab*_(*t*) = *p*_*aB*_(*t*). Consider a population in which all strains are present at equal frequencies and there is an equal number of uninfected hosts with immunity to each strain, that is, *D*(*t*) = 0 and 𝒜_Δ_(*t*) = 0. Moreover, suppose one set of strains, say *{ab, AB}*, increases in frequency relative to the set of strains *{Ab, aB}*, due to some new infections, at least some of which occur in naive hosts. This results in *D*(*t*) *>* 0, that is, the set of strains *{ab, AB}* is overrepresented. As these infections are cleared, this produces an excess of uninfected hosts with immunity to the overrepresented strains *{ab, AB}*, that is, *S*_*AB*_(*t*)+*S*_*ab*_(*t*) *> S*_*Ab*_(*t*)+*S*_*aB*_(*t*), or 𝒜_Δ_(*t*) *>* 0. This excess of uninfected hosts will have immunity to a single antigen of a randomly chosen strain belonging to the set of underrepresented strains, *{Ab, aB}*, and immunity to either 0 or 2 antigens of a randomly chosen strain belonging to the set of overrepresented strains *{ab, AB}*. The sign of *σ*_𝒜_ then determines three possible cases:

1. If *σ*_𝒜_ *<* 0, then *σ*_1_ *>* (*σ*_0_ + *σ*_2_)*/*2. Here, the excess of uninfected hosts is, on average, more susceptible to infection by a randomly chosen strain belonging to the set of underrepresented strains, *{Ab, aB}*, than a randomly chosen strain belonging to the set of overrepresented strains, *{ab, AB}*. Consequently, the underrepresented strains will cause more infections per capita (i.e., have higher fitness) and so increase in frequency. As a result, the strains will return to an equal frequency, that is, *D*(*t*) → 0.
2. If *σ*_𝒜_ *>* 0, then *σ*_1_ *<* (*σ*_0_ + *σ*_2_)*/*2. Here, the excess of uninfected hosts is more susceptible to infection by a randomly chosen strain belonging to the set of overrepresented strains, *{ab, AB}*, than the set of underrepresented strains, *{Ab, aB}*. Thus, the overrepresented strains will further increase in frequency. This creates a positive feedback loop that eventually drives the set of overrepresented strains to fixation, that is, *D*(*t*) → 1*/*4.
3. Finally, if *σ*_𝒜_ = 0, then *σ*_1_ = (*σ*_0_ + *σ*_2_)*/*2. Here, the excess of uninfected hosts is equally susceptible to infection by any strain. Consequently, neither set of strains will change in frequency. However, if there is any recombination in the population, that is, *ρ >* 0, recombination will systematically remove LD. Recombination will therefore reduce the frequency of the set of overrepresented strains, returning the population to a state at which all strains have equal frequency.

To show this mathematically, note that in the absence of waning immunity and virulence mortality,

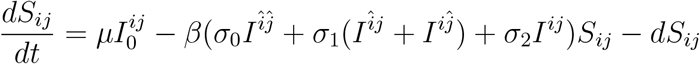

where 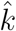 means the allele opposite to *k* at the indicated locus, e.g., â = *A*. Therefore, we can write

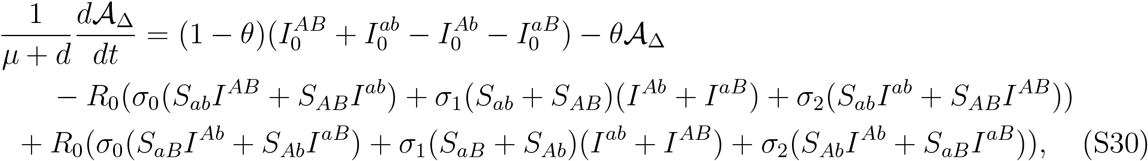

which, on the slow timescale, reduces to

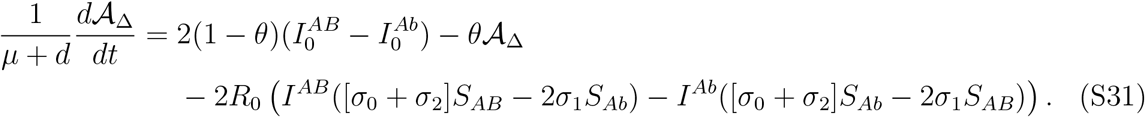

Now, suppose we are at the equilibrium state in which there is no strain structure. In this case, *D*(*t*) = 0 and *S*_*AB*_(*t*) = *S*_*Ab*_(*t*) = *S*_2_(*t*). Imagine that infections by the set of strains *{ab, AB}* slightly increase in density,

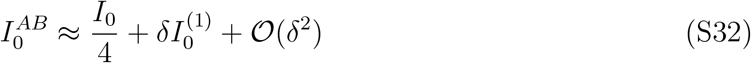

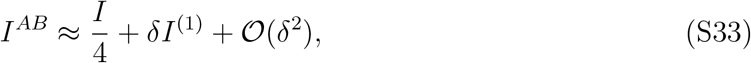

where *I*_0_ is the total density of strains (irrespective of genotype) infecting naive hosts, 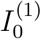 and *I*^(1)^ are the (relative) perturbed values of 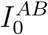 and *I*^*AB*^, respectively, and 0 *< δ* ≪ 1 is a small quantity. In this case,

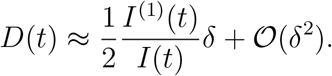

Using these relations, equation (S31) reduces to

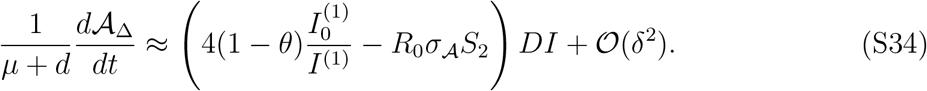

Next, note that *S*_2_ is the number of hosts with immunity to a single strain; in the vicinity of the no strain structure equilibrium, the dynamics of *S*_2_ are given by

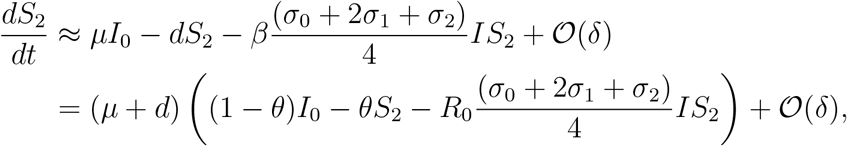

and so

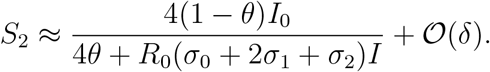

Hence,

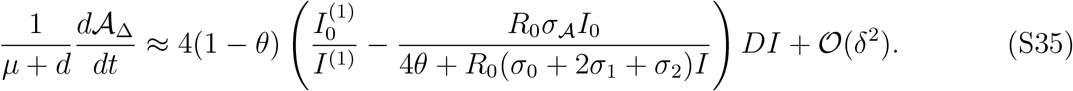

It therefore follows that if

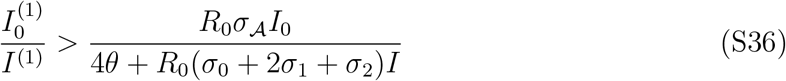

then the increase in strains *{ab, AB}* due to the perturbation results in 𝒜_Δ_(*t*) and *D*(*t*) sharing sign, in which case, if *σ*_𝒜_ *>* 0, *D*(*t*) will increase, whereas if *σ*_𝒜_ *<* 0, *D*(*t*) will decrease. Inequality (S36) always holds if *σ*_𝒜_ *<* 0. If *σ*_𝒜_ *>* 0, inequality (S36) will also hold for biologically ‘reasonable’ perturbations. Here, a biologically reasonable perturbation should be roughly proportional to the existing distribution of infections, that is,

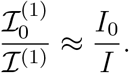

That is because the existing distribution of infections (at equilibrium) indicates the relative likelihood of infection based upon immune history for a given set of parameters. Hence, *I*_0_*/I* measures the relative infection risk of a naive host.

##### S4.1.2 Conflicting evolutionary pressures induced by *σ*_𝒮_ *>* 0

Next, suppose *σ*_𝒮_ *>* 0. Then

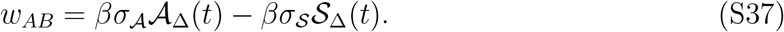

Since 𝒮_Δ_(*t*) consists of uninfected hosts that have been infected only once and uninfected hosts that have been infected multiple times, it can diverge from 𝒜_Δ_(*t*). This can create a second epidemiological feedback on epistasis. In general, this second epidemiological feedback removes LD. To see why, assume that *σ*_𝒜_ = 0. As before, suppose that we start from a state in which all strains are at equal frequency and there is an equal number of uninfected hosts with immunity to each strain, i.e., *D*(*t*) = 0 and 𝒜_Δ_(*t*) = 𝒮_Δ_(*t*) = 0. Then suppose that one set of strains, say *{ab, AB}*, increases in frequency relative to the other through additional infections. This leads to *D*(*t*) *>* 0, that is, the set of strains *{ab, AB}* is overrepresented. As hosts clear these infections, an excess of uninfected hosts with homologous immunity to the strains *{ab, AB}* appears, and so 𝒮_Δ_(*t*) *>* 0. This excess of uninfected hosts has immunity to a single antigen of the underrepresented strains, *{Ab, aB}*, and either no immunity or homologous immunity to the overrepresented strains, *{ab, AB}*. Thus, for this excess of uninfected hosts, the per-capita susceptibility to infection by a randomly chosen member of the set of underrepresented strains is *σ*_1_, and the per-capita susceptibility to infection by a randomly chosen member of the set of overrepresented strains is (*σ*_0_ + *σ*_*h*_)*/*2. Since *σ*_𝒜_ = 0, this implies *σ*_1_ = (*σ*_0_ + *σ*_2_)*/*2, and therefore

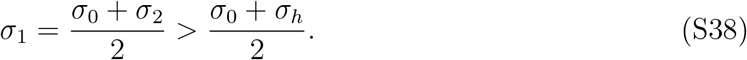

Thus, the excess of uninfected hosts is more susceptible to infection by the underrepresented strains, *{Ab, aB}*, and so the underrepresented strains will increase in frequency. This will return *D*(*t*) → 0. If *σ*_𝒜_ *<* 0 in addition to *σ*_𝒮_ *>* 0, this implies *σ*_1_ *>* (*σ*_0_ + *σ*_2_)*/*2, and so inequality (S38) will be more pronounced. Thus, the per-capita advantage of the underrepresented strains increases. Consequently, we would again expect *D*(*t*) → 0. However, if *σ*_𝒜_ *>* 0, inequality (S38) will not hold. In this case, the two epidemiological feedbacks become opposed. This can either produce evolutionary oscillations or result in a stable nonzero value of LD that is not equal to *D*^*^ (i.e., at equilibrium, *w*_*AB*_ = 0 and 0 *<* |*D*(*t*)| *<* 1*/*4).

Due to the high dimensionality of the model, analytical results are difficult to obtain. However, in the next section we improve the mathematical prediction of the stability of the PSS equilibrium when *σ*_𝒮_ = 0.

#### S4.2 Epistasis and LD must share sign for local asymptotic stability when *σ*_𝒮_ = 0

Taking the derivative of the differential equation for LD, 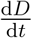, given in system (S22), with respect to any of the population variables (arbitrarily indicated as *X*) under the assumption that recombination is negligible (*ρ ≈* 0), we obtain:

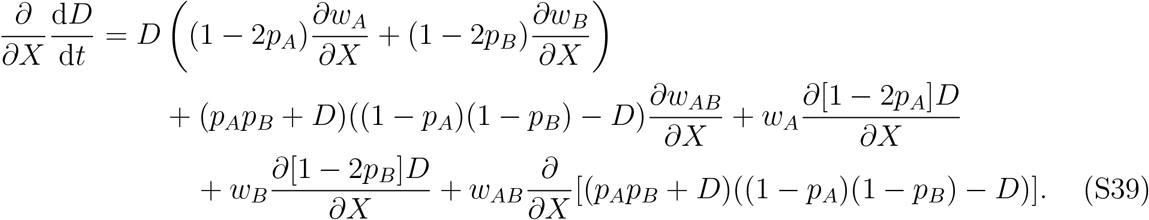

We evaluate (S39) at *p*_*A*_ = *p*_*B*_ = 1*/*2 and *D* = 1*/*4 or *D* = *−*1*/*4; these are the stable PSS equilibria that occur when *σ*_𝒮_ = 0. Since *r*_*AB*_ = *r*_*ab*_ and *r*_*Ab*_ = *r*_*aB*_, and so *w*_*AB*_ = *−*2*w*_*A*_ = *−*2*w*_*B*_, equation (S39) reduces to

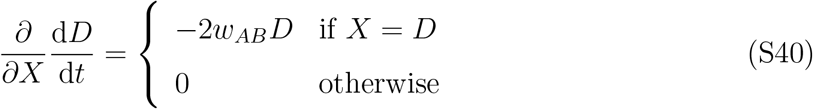

Here, the Jacobian of the full system that combines both the evolutionary and epidemiological variables, evaluated at either of the stable PSS equilibria, becomes block triangular, with one of the ‘blocks’ on the diagonal given by equation (S40) (when *X* = *D*). Thus, one of the eigenvalues is *−w*_*AB*_*/*2 (if *D* = 1*/*4) or *w*_*AB*_*/*2 (if *D* = *−*1*/*4). It follows that a necessary condition for PSS to be locally asymptotically stable when *σ*_*S*_ = 0 is that the sign of epistasis matches the sign of LD.

Now, on the slow timescale, the sign of *D*(*t*) indicates whether there are more hosts currently infected by members of the set of strains *{AB, ab}* (if *D*(*t*) *>* 0) or by members of the set of strains *{Ab, aB}* (if *D*(*t*) *<* 0). Therefore *D*(*t*) and 𝒜_Δ_(*t*) share sign whenever more uninfected hosts have full antigenic immunity to the set of overrepresented strains. Indeed, as |*D*(*t*)| → 1*/*4, only one set of strains will be present in the population. Therefore, if hosts gain or lose immunity to a set of antigens that define an infecting (or previously infecting) strain, at the stable PSS equilibrium

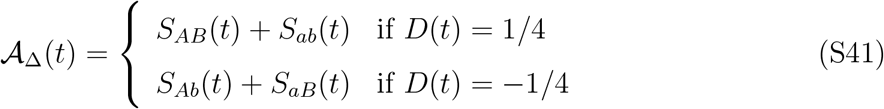

Applying this result to equation (S40), it follows that when *σ*_𝒜_ *>* 0, epistasis and LD share sign at the PSS equilibrium, otherwise they do not. Thus, a *necessary* condition for local asymptotic stability of the PSS equilibrium when *σ*_𝒮_ = 0 is that *σ*_𝒜_ *>* 0. The high dimensionality of the system impeded the derivation of the necessary and sufficient conditions. In the next section, we therefore numerically explore the predictions.

#### S4.3 Numerical simulations of key predictions

In the previous sections, we have provided intuition for our key predictions: (i) if *σ*_𝒮_ = 0 and *σ*_𝒜_ *>* 0, stable PSS occurs (we also proved this is the necessary condition); (ii) if *σ*_𝒮_ = 0 and *σ*_𝒜_ *<* 0, no PSS occurs; (iii) evolutionary oscillations can only occur if both *σ*_𝒮_ *>* 0 and *σ*_𝒜_ *>* 0. Here, we provide numerical simulations to confirm these predictions. In the main text, we show (numerically) that evolutionary oscillations are possible for *σ*_𝒮_ *>* 0 and *σ*_𝒜_ *>* 0 (see Fig. 3 and Fig. 4); furthermore, we show that when *σ*_𝒮_ = 0, the sign of *σ*_𝒜_ predicts the (stable) outcome of either no PSS (*σ*_𝒜_ *<* 0) or stable PSS (*σ*_𝒜_ *>* 0). Here, we expand the simulations for the case when *σ*_𝒮_ = 0 to also show the dynamics of 𝒜 _Δ_(*t*) in relation to *D*(*t*).

For our simulations, we used the model specified by equations (S7), and assumed that: (i) immunity is lifelong and acquired at time of clearance, (ii) immune protections only affect susceptibility to infection (so 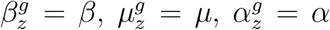), and (iii) we fixed *b* = *d* = 0.01, *σ*_0_ = 1, *σ*_2_ = *σ*_*h*_ = 0, *α* = 0, and *ρ* = 10^*−*8^. We then looped over 40 evenly spaced values of *R*_0_ ∈ [1.3, 20], 40 evenly spaced values of *θ* ∈ [5 *×* 10^*−*5^, 0.5], and 20 evenly spaced values of *σ*_1_ ∈ [0, 1]. For each parameter combination, we generated a random initial condition at *t* = 0, subject to the constraint that the total population density is 1. Thus, the initial condition is not on the slow timescale. We then numerically integrated the system of ODEs until an equilibrium is reached; we determined this numerically by integrating forward in time until

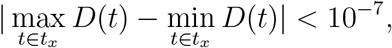

where *t*_*x*_ is the last 10^4^ time units. Figure S3 shows the outcome of these 3.2*×*10^4^ simulations. Three notable implications arise from the simulations. First, 𝒜 _Δ_(*t*) and *D*(*t*) share sign (Fig. S3**a, b**), except during the fast timescale transient dynamics (these fast timescale dynamics are the horizontal trajectories in panels **a**,**b**; here, 𝒜 _Δ_(*t*) is rapidly changing while *D*(*t*) is not). Second, in all 3.2*×*10^4^ simulations, the sign of *σ*_𝒜_ predicts the long-term outcome (Fig. S3**c**): when *σ*_𝒜_ *>* 0, |*D*(*t*)| → 1*/*4, whereas when *σ*_𝒜_ *<* 0, |*D*(*t*)| → 0. Finally, sustained evolutionary oscillations were not observed in any of the simulations, in agreement with our prediction that these can only occur if both *σ*_𝒜_ *>* 0 and *σ*_𝒮_ *>* 0.

### S5 Different types of immune protections

Next, we consider a more general case in which we assume that immune protections may affect transmissibility, clearance, and virulence mortality, but that homologous immunity is equivalent to full antigenic immunity. We obtain an expression for epistasis in fitness, which is more complicated than when immunity only affects susceptibility. That is because if immunity affects transmissibility, clearance, and/or virulence-mortality, the fitness (percapita growth rate) of an infection will depend upon the immune environment the infection is found in. As a consequence, the per-capita growth rate of strain *ij* depends upon the frequency of strain *ij* infections found in each particular environment. Therefore, we focus upon deriving epistasis in fitness at equilibrium. In this case, there is one epidemiological feedback, which is whether more hosts have antigenic immunity to members of one set of strains or the other (e.g., *{ab, Ab}* or *{Ab, aB}*). Hence, if recombination is weak, there is no reason to expect epistasis to become zero and/or change sign at some non-zero amount of LD. Consequently, we expect that if PSS occurs, it will be the largest in magnitude possible given the allele frequencies, that is, *D*(*t*) → *D*^*^. Specifically, if *p*_*A*_(*t*) = *p*_*B*_(*t*) = 1*/*2, and if PSS occurs, then |*D*(*t*)| → 1*/*4 (this means two strains will be excluded from the population). We will focus upon these two equilibria here.

**Figure S3:**
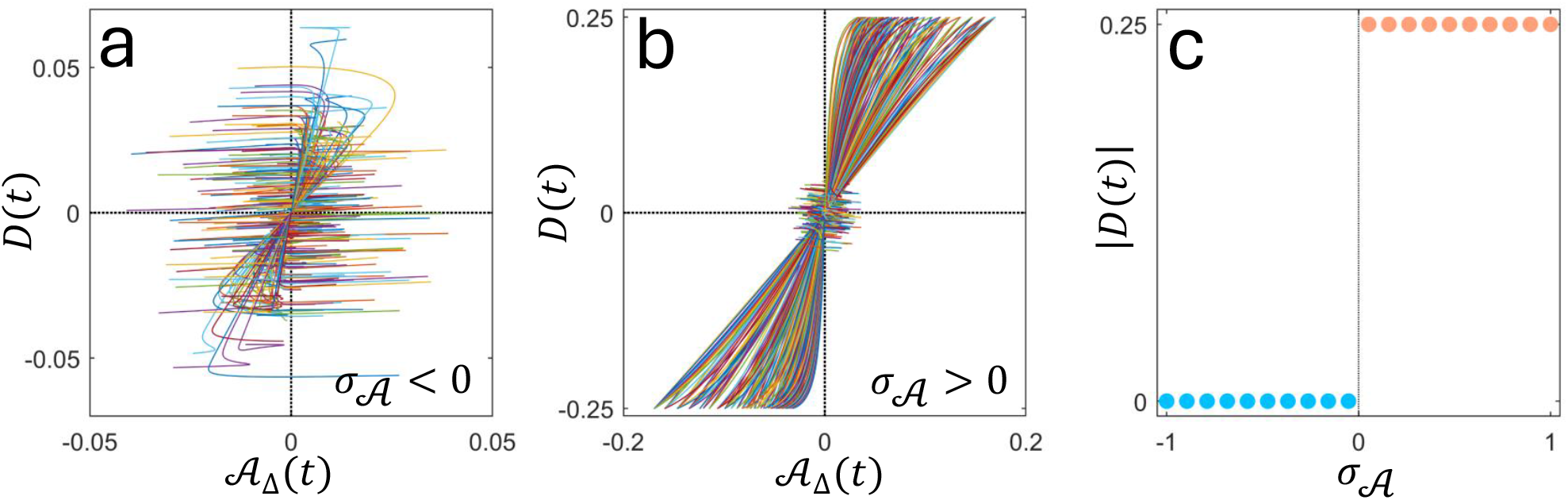
Relationship between LD and 𝒜 _Δ_(*t*) for different values of *R*_0_ and *θ*. Numerical simulations show that 𝒜 _Δ_(*t*) and *D*(*t*) generally share sign, regardless of whether *σ*_𝒜_ > 0 (panel **a**) or *σ*_𝒜_ < 0 (panel **b**); any divergence is transient, occurring on the fast timescale (horizontal trajectories in panels **a**,**b**). In the long term, the evolutionary outcome is determined by the sign of *σ*_𝒜_: if *σ*_𝒜_ < 0, *D*(*t*) *→* 0, whereas if *σ*_𝒜_ > 0, |*D*(*t*)| *→* 1*/*4 (panel **c**). Shown are the results of 3.2 × 10^4^ simulations, in which we looped over 40 evenly spaced values of *R*_0_ ∈ [1.3, 20], 40 evenly spaced (on log-scale) values of *θ ∈* [5 × 10^−5^, 0.5] and 20 evenly spaced values of *σ*_1_ ∈ [0, 1]. All simulations assume *b* = *d* = 0.01, *ρ* = 10^−8^, *σ*_0_ = 1, *σ*_2_ = *σ*_h_ = 0, and *α* = 0.

To calculate epistasis at equilibrium we define

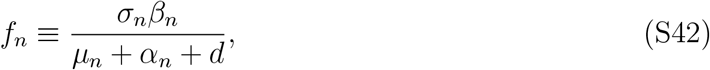

which is proportional to the effective reproductive number of an infection in a host with immunity to *n* antigens of the infecting strain. This is because, an infection is established in a host with immunity to *n* of its antigens with probability *σ*_*n*_; such an infection has an expected lifetime of (*μ*_*n*_ + *α*_*n*_ + *d*)^*−*1^, during which it will produce new infections at a rate proportional to *β*_*n*_.

Next, at equilibrium, the reproductive number of strain *ij* [e.g., see 83], which we denote 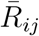, is

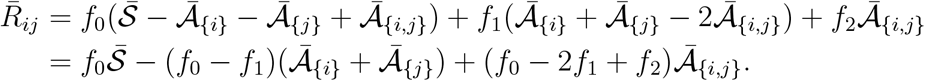

Now, the per-capita growth rate, *r*_*ij*_, is a measure of fitness in continuous time, whereas the reproductive number,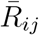, is a measure of fitness in discrete time [83]. This can be explicitly shown using, for example, the Next Generation Theorem [84, 85], a technique which converts a continuous-time system into a discrete time system [83]. Whereas in continuous time, epistasis in fitness is defined as

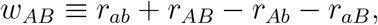

in discrete time, epistasis in fitness becomes

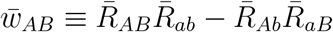

[see e.g., 86, 87]. Without loss of generality, we focus upon the equilibrium in which the set of overrepresented strains is *{ab, AB}*. At this equilibrium, 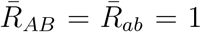 moreover, 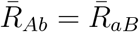 Therefore, When (_*pA, pB*,_*D*) = (1/2, 1/2, 1/4),

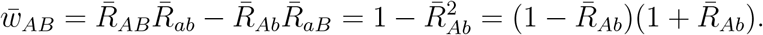

Since 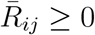,

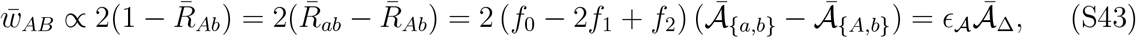

where proportionality is up to a positive constant, ϵ_𝒜_= *f*_0_ − 2*f*_1_ + *f*_2_, and 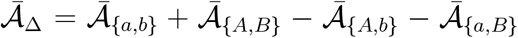 (note that 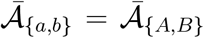 and 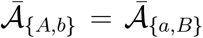 at equilibrium). Hence, if *ϵ* _𝒜_ *>* 0, epistasis and LD will share sign.

There are four important consequences of the inclusion of different types of immune protections.

1. First, *ϵ*_𝒜_ is a decreasing function of *σ*_1_, *β*_1_, and *α*_1_, and is an increasing function of *μ*_1_. Thus, increasing the strength of immune protections targeting transmission (smaller *β*_1_), susceptibility to infection (smaller *σ*_1_), and clearance rate (larger *μ*_1_) promotes PSS by making it more likely *ϵ*_𝒜_ is positive. On the other hand, increasing the strength of immune protections targeting virulence mortality (smaller *α*_1_) inhibits PSS by making it more likely *ϵ*_𝒜_ is negative.
2. Second, if immunity only reduces virulence mortality, then genetic polymorphisms cannot be stably maintained. This can be seen by supposing only one strain is present, say *AB*, and that the population is at its endemic equilibrium. If, in addition, homologous immunity is as protective as full antigenic immunity and there is no waning immunity, uninfected hosts can be either naive or have immunity to the antigens *AB*. Denote the equilibrium density of naive hosts as 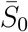, and the equilibrium density of uninfected hosts with immunity to antigens *AB* as 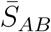. Then strain *Ab* and/or strain *aB* can increase in frequency when rare if they have a higher basic reproductive number than strain *AB*, that is,

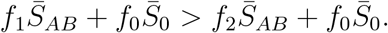 Likewise, strain *ab* can increase in frequency when rare if

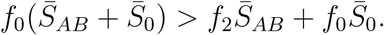 If immunity only reduces virulence mortality, that is, *α*_2_ *< α*_1_ *< α*_0_, then *f*_0_ *< f*_1_ *< f*_2_ and so neither inequality is satisfied. Thus if one strain is dominant in the population, it can out compete all the other strains. Consequently, if immunity only acts to reduce virulence mortality, it cannot maintain genetic polymorphisms.
3. Third, suppose only one type of immune protection is operating. In particular, suppose immunity either reduces susceptibility to infection, reduces transmissibility, or increases clearance rate. Furthermore, suppose that whatever type of immune protection is present, its strength additively increases with the number of antigens that a host has been exposed to, that is,

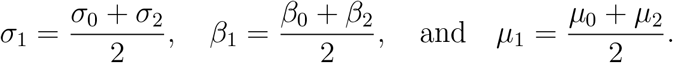 If immunity affects only transmissibility or susceptibility to infection, then *ϵ*_𝒜_ = 0. Thus PSS can only occur for transmissibility or susceptibility if immune protections are saturating, that is, the increase in immunity is more than additive (this is also shown in Figure 2). However, if immunity affects only clearance rate, then an additive increase in immune protections yields

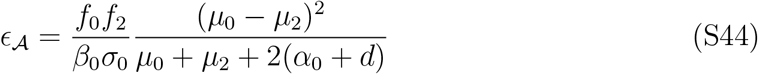 As this is positive, immunity affecting clearance rate is more effective at producing PSS (Fig. S4). Besides, we would generally expect the opposite prediction for immune protections that reduce virulence mortality, provided there is another type of immune protection maintaining genetic polymorphisms. For example, suppose the strength of immune protections additively increases, and affects both susceptibility and virulence mortality. Further, assume that *f*_0_ *> f*_1_ and *f*_0_ *> f*_2_, that is, genetic polymorphisms at both loci are stably maintained. Then

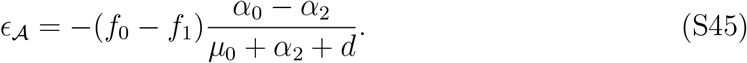 Since *f*_0_ *> f*_1_ and *α*_0_ *> α*_2_, it follows that *ϵ*_𝒜_ *<* 0, that is, no strain structure is favoured when immune protections against virulence mortality are additively increasing. Thus immunity affecting virulence mortality is less effective at producing strain structure (Fig. S4). Indeed, suppose immunity affects only clearance rate and we solve for the value of *μ*_1_, say 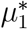, at which *ϵ*_𝒜_ = 0. Thus if 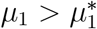 PSS occurs. Solving the relation *θ* = *d/*(*μ*_0_ +*α*_0_ +*d*) for *d*, and using this in 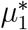, we obtain

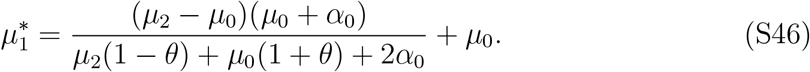 This is an increasing function of *θ* and *α*_0_. Therefore, PSS is increasingly likely (smaller 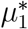) when infections become more acute (*θ* → 0) and infections become less virulent (*α*_0_ → 0 for fixed *θ*), as claimed in the main text.
4. Finally, the derivation of equation (S43) assumes that homologous immunity is equivalent to full antigenic immunity. The analogous quantity to *σ*_𝒮_ is *ϵ*_𝒮_ ≡ *f*_2_ *− f*_*h*_. We conjecture that *ϵ*_𝒮_ *>* 0 is a necessary condition for oscillatory PSS.

### S6 Variation in immune protection between alleles and and/or strains

The key assumption facilitating the separation of timescales in Section S3.4 is that the alleles at each locus, and the combinations of alleles, are of equal intrinsic fitness, i.e., they induce identical immune protections and do not differ with respect to any other life-history characteristic (e.g., transmissibility [13, 42]). Thus, all hosts with immunity to (for example) one antigen of a strain have the same level of protection, irrespective of the identity of the antigen. As a result, allele frequencies rapidly converge to 1*/*2. After that, the LD, which determines the amount of strain structure, evolves on the slow timescale.

**Figure S4:**
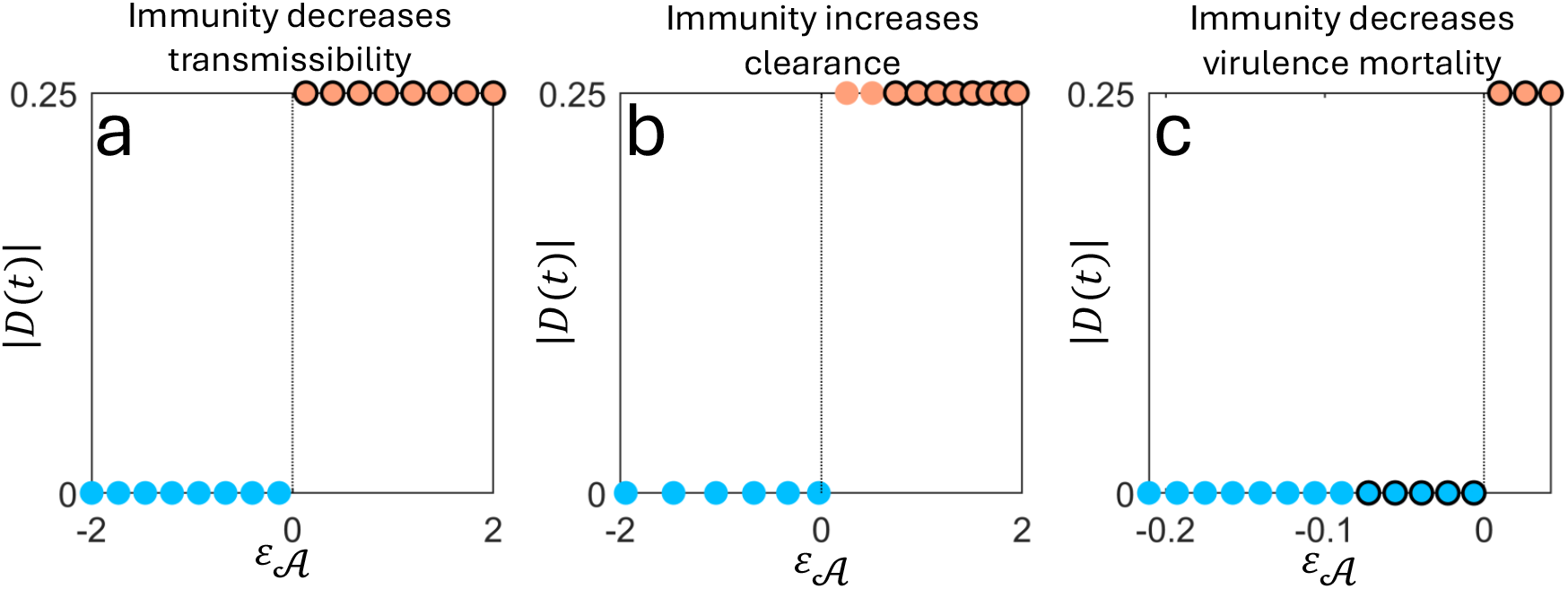
The ability of different types of immune protections to produce PSS can be predicted by the sign of *ϵ*_𝒜_. In panel **a**, immunity only reduces transmissibility (i.e., *σ*_n_ = *σ*_0_, *μ*_n_ = *μ*_0_, and *α*_n_ = *α*_0_), and we vary *β*_1_ over 16 evenly-spaced values between *β*_0_ and *β*_2_ = *β*_0_*/*2 inclusive. Here, *ϵ*_𝒜_ = 0 when *β*_1_ = (*β*_0_+*β*_2_)*/*2, and so an equal number of circles are to either side of the dashed black line (circles with black edges indicate *β*_1_ > (*β*_0_ + *β*_2_)*/*2). In panel **b**, we assume immunity only effects clearance rate (i.e., *σ*_n_ = *σ*_0_, *β*_n_ = *β*_0_, and *α*_n_ = *α*_0_), and vary *μ*_1_ over 16 evenly-spaced values between *μ*_0_ and *μ*_2_ = 2*μ*_0_ inclusive. Because immune protections affecting clearance rate are more likely to produce PSS, *ϵ*_𝒜_ = 0 for *μ*_1_ < (*μ*_0_ + *μ*_2_)*/*2, and so more circles are to the right of the dashed black line than the left (circles with black edges indicate *μ*_1_ > (*μ*_0_ + *μ*_2_)*/*2). Finally, in panel **c**, immunity only reduces virulence mortality and susceptibility (i.e., *β*_n_ = *β*_0_ and *μ*_n_ = *μ*_0_). Here, we assume that immunity additively reduces susceptibility, *σ*_1_ = (*σ*_0_ + *σ*_2_)*/*2, and vary *α*_1_ over 16 evenly-spaced values between *α*_2_ = 0 and *α*_0_ inclusive. Because immune protections affecting virulence mortality are less likely to produce PSS, *ϵ*_𝒜_ = 0 for *α*_1_ > (*α*_0_ + *α*_2_)*/*2. Thus, more circles are to the left of the dashed black line than to the right (circles with black edges indicate *α*_1_ < (*α*_0_ + *α*_2_)*/*2). Unless otherwise noted, all simulations assume *μ*_0_ = 9.49, *β*_0_ = 40, *α*_0_ = 0.5, *b* = *d* = 0.01, *σ*_0_ = 1, and *ρ* = 10^−9^; thus *R*_0_ = 4, *θ* = 10^−3^, and the probability a naive host dies from an infection, *α*_0_*/*(*μ*_0_ + *α*_0_ + *d*), is 0.05.

The symmetry between alleles (and strains) could be disrupted in at least two ways:

(i) if there is an unequal number of alleles at the different loci, preventing all alleles from coexisting at equal frequencies, or (ii) if the different alleles and/or combinations of alleles induce different levels of immune protection. In Section S8, we explore what happens when the number of alleles at each loci is not identical. Here, we examine how variation in the immune protections against each allele, and each combination of alleles, affects our predictions. We investigate two key predictions: (1) whether homologous immunity being more protective than full antigenic immunity is necessary for sustained evolutionary oscillations, and (2) whether, if homologous immunity is as protective as full antigenic immunity and so oscillations are not possible, then *σ*_𝒮_ *>* 0 implies that LD stably persists at either its maximum (equation (S26)) or minimum (equation (S27)) value, whereas if *σ*_𝒮_ *<* 0 no strain structure occurs.

To test these predictions, we define 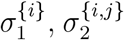, and 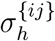, as the reduction in susceptibility to an infection by strain *ij* of an uninfected host with immunity to antigen *i* but not *j*, immunity to both antigens *i* and *j*, and homologous immunity, respectively. We assume that full antigenic immunity is as protective as homologous immunity for each combination of antigens, that is, 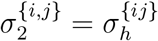. Then the per-capita growth rate of strain *ij* is

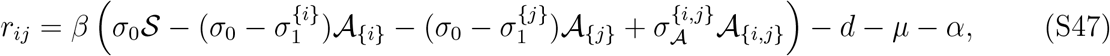

where

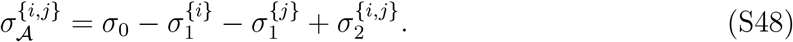

Consequently, the additive selection coefficients and epistasis are

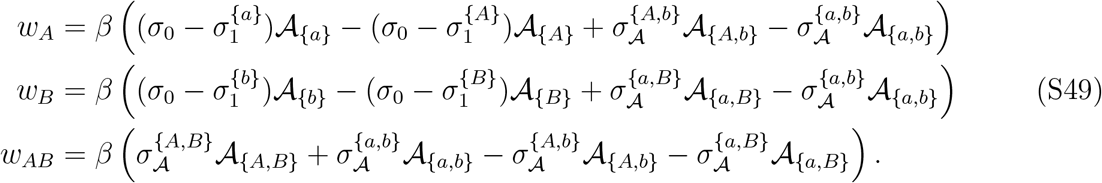

Because the amount of LD possible depends upon the allele frequencies, we test our predictions using a normalized version of LD [88], *D*^*′*^, defined as

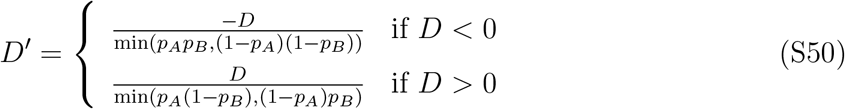

Here, if *D*^*′*^ = 0, there is no PSS. If *D*^*′*^ = 1, there is the maximum possible amount of PSS, given allele frequencies. Further, we test our predictions for the sign of *σ* _𝒜_ using the sign of the value of *σ* _𝒜_ averaged across allelic combinations:

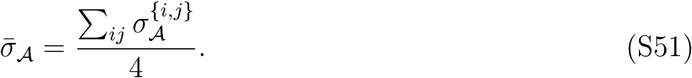

Here, epistasis is no longer the product of a composite parameter, *σ*_𝒮_, and a composite variable 𝒜_Δ_(*t*), which makes analytic progress difficult. Therefore, we rely on simulations. To do so, we let the protection against alleles *i* and *j* be

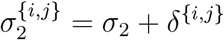

where *δ*^{*i,j*}^ ∼ *𝒩* (0, *v*_2_) is a normally-distributed random variable with mean 0 and variance *v*_2_, and we let the protection against the alleles *k* ∈ *{a, b}* and *K* ∈ *{A, B}* be

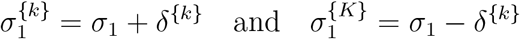

where *δ*^*{k}*^ ∼ *𝒩* (0, *v*_1_) is a normally-distributed random variable with mean 0 and variance *v*_1_. We adjust the immune protection of allele *k* and *K* by *±* the same amount so that fitness variation between alleles at the same locus does not cancel on average.

If there is no variation in protection against combinations of alleles, that is,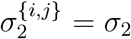, then we can measure the total deviation from symmetry for a randomly generated set of 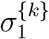 as

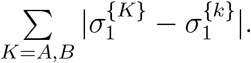

The logic here is that when 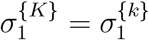, there is symmetry between alleles; as the difference grows, so too will the deviation. Likewise, if there is no variation in protection against single alleles, that is, 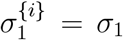, then we can measure the total deviation from symmetry for a randomly generated set of 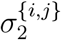 using strain *ab* as reference,

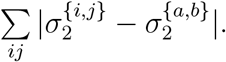

Figures S5 and S6 show the outcome of numerical simulations using randomly chosen values of *R*_0_, *θ*, 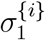, and 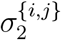. Each simulation was run until an equilibrium was reached, which was determined by the condition

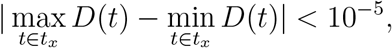

where *t*_*x*_ is the last 10^4^ time units. Two notable implications emerge from the simulations. First, for all simulations, the system eventually converges to an equilibrium; sustained evolutionary oscillations were not possible. This suggests that our prediction that the necessary condition for evolutionary oscillations is a divergence between homologous immunity and full antigenic is robust to variation in immune protections against alleles and strains. Note, however, due to the variation between strains in intrinsic fitness, stable PSS no longer needs to correspond to *D*^*′*^ = 1; it is possible that *w*_*AB*_ = 0 for some intermediate value of *D*^*′*^. Second, the sign of 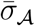 becomes an increasingly worse predictor of whether PSS occurs as the variation in immune protections against alleles and strains increases. This is particularly true when (1) 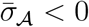 as compared to 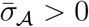 or (2) 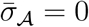. In case (1), individual variation in the fitness of alleles and strains creates additional epistatic pressure favouring individual strains, which is capable of building up LD. In case (2), even if 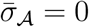, this need not mean that epistasis is zero.

## S7 The link between strain frequencies and the immunological landscape

As discussed in the main text, various epidemiological assumptions can disrupt the correlation between which set of strains is overrepresented in the population, and which set of strains has more hosts with immunity to it (so *D*(*t*) and 𝒜_Δ_(*t*), 𝒮_Δ_(*t*) do not share sign).

Here we explore two ways in which this disruption can occur; in both we assume for simplicity that full antigenic immunity is equivalent to homologous immunity, *σ*_*h*_ = *σ*_2_. In the first example, suppose uninfected hosts lose immunity to each antigen at a per-capita rate of *ω*. For example, an uninfected host with immunity to antigens {*a, A, b*} becomes immune to antigens {*a, A*} at a per-capita rate *ω*. In the second example, suppose that following clearance of infection, hosts gain immunity to a single, randomly selected antigen with probability *p*, and both antigens with probability 1 − *p*.

**Figure S5:**
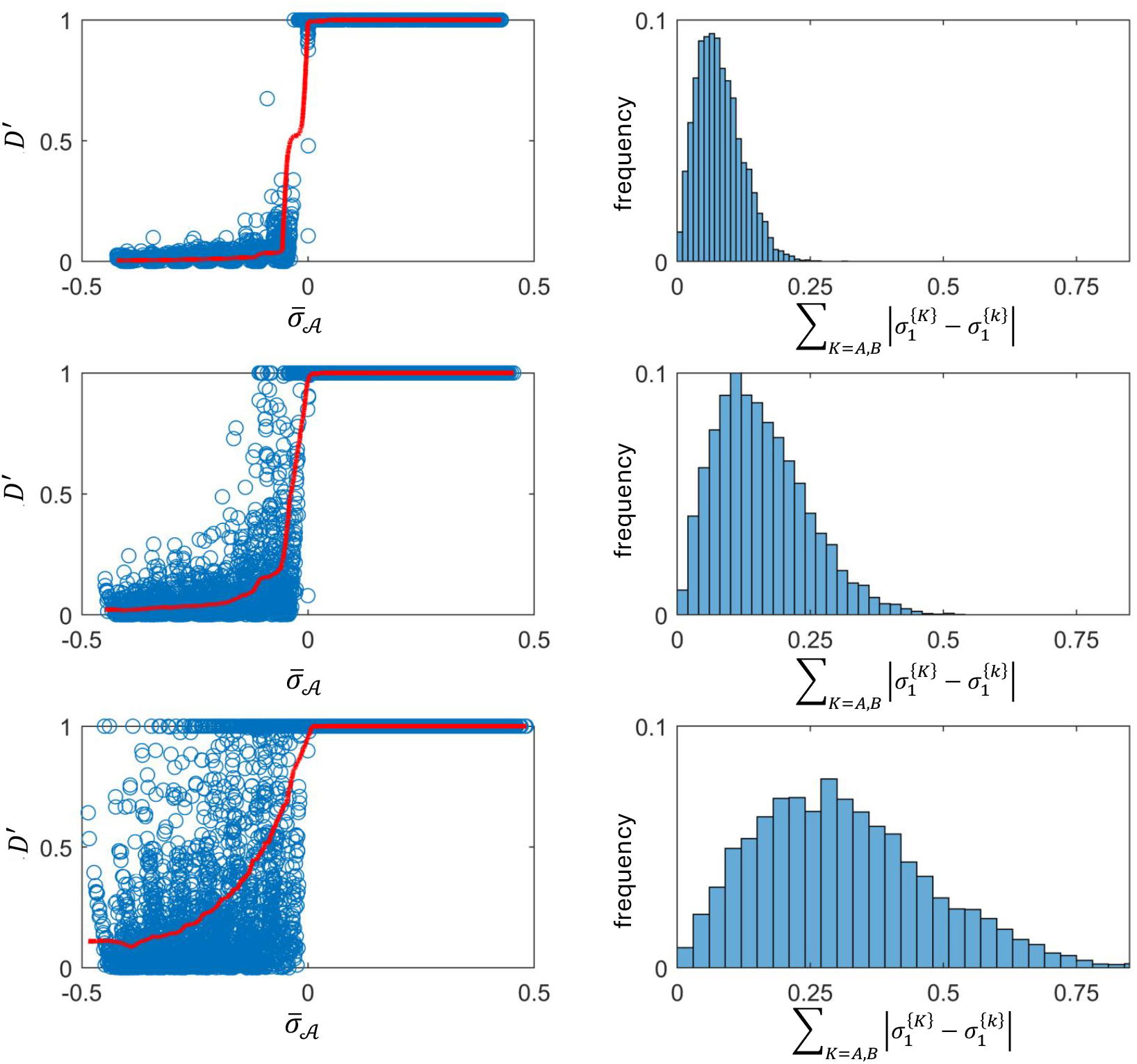
Variation in immune protections against alleles and the predictive ability of *σ*_𝒮_. Here, we vary the value of *σ*_1_ for each allele, specifically, 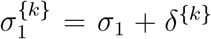 and 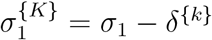, where *δ*^{*k*}^ *∼* 𝒩(0, *v*_1_), that is, *δ*^{*k*}^ is a normally-distributed random variable with mean 0 and variance *v*_1_. For each row we vary the value of *v*_1_: for the first row, *v*_1_ = 0.025; for the second row, *v*_1_ = 0.05; for the third row, *v*_1_ = 0.1. We then loop over 15 evenly-spaced values of *σ*_1_ ∈ [0.4, 0.8]. For each value of *σ*_1_, we perform 400 simulations (so each panel is 6 *×* 10^3^ simulations), where in each simulation we randomly set allelic fitness variation (i.e., the values of *δ*^{*k*}^), as well as the values of *θ* and *R*_0_; these are uniform random variables in the intervals *θ* ∈ [10^−4^, 0.5] and *R*_0_ ∈ [1.5, 18]. The first column shows how the sign of 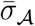 affects the equilibrium value of *D*^*′*^. The second column shows the distribution of fitness variation used in the simulations. The red line is the rolling average of *D*^*′*^ using a window of size 400. All simulations assume *σ*_0_ = 1, *σ*_2_ = 0.2, *b* = *d* = 0.01, *α* = 0 and *ρ* = 10^−8^.

Since homologous immunity is of the same strength as full antigenic immunity, suppose the immune history *x* for these examples only tracks the unique antigens against which a host has immunity (e.g., in the previous formulation, *x* = {*ab, Ab*}, would become *x* = {*a, A, b*}. The two assumptions above leave the equations for the 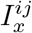 unchanged, but the net rate at which immune history *x* is gained and lost is

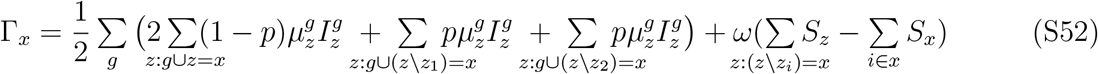

where (*z\z*_*i*_) indicates the genotype *z* without the *i*-th locus; e.g., if *z* = *ab*, then (*z\z*_1_) = *b*.

**Figure S6:**
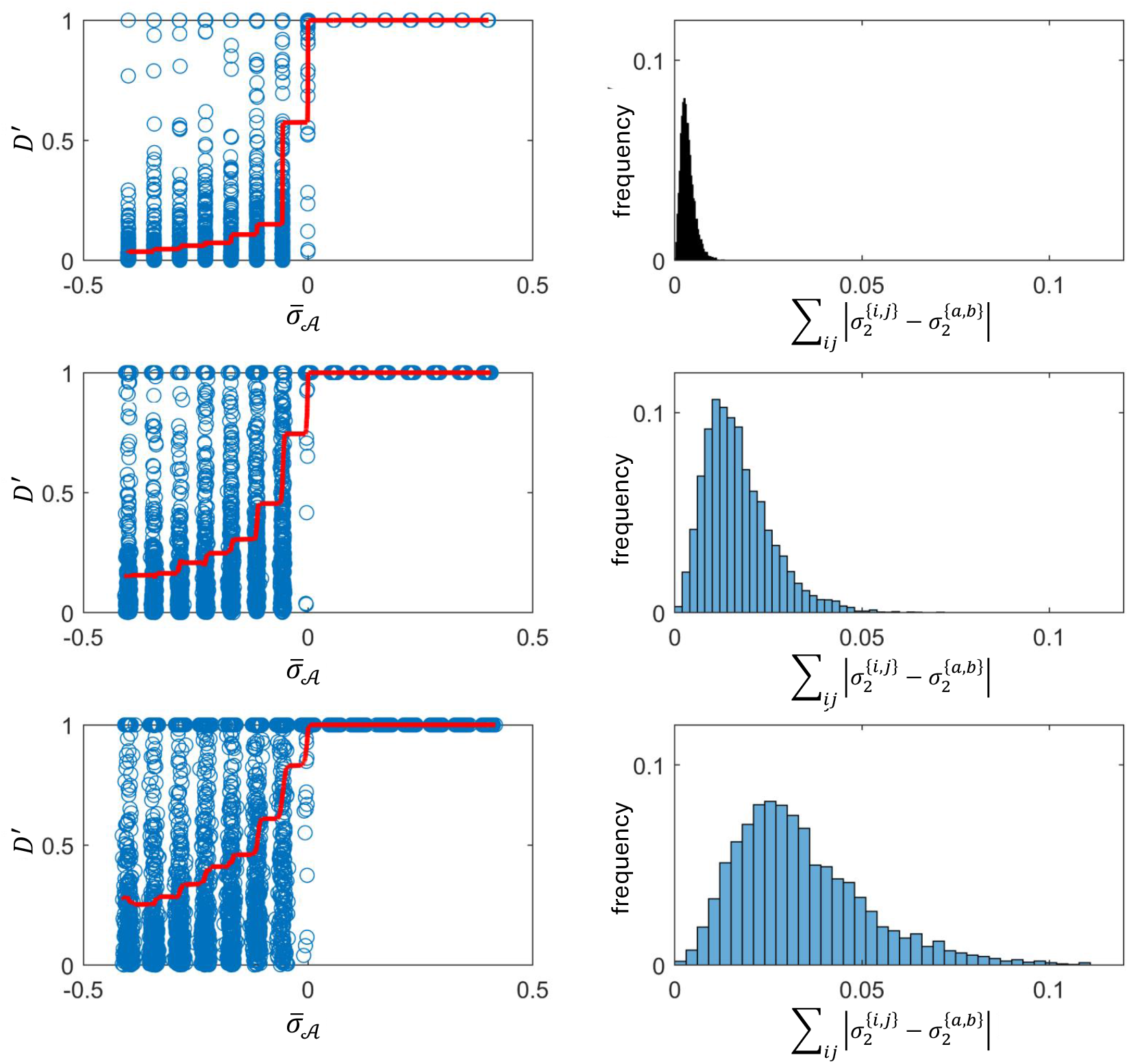
Variation in immune protections against combinations of alleles and the predictive ability of *σ*_𝒮_. Here we vary the value of *σ*_2_ for each allele, specifically, 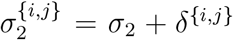, where *δ*^{*i,j*}^ *∼ 𝒩* (0, *v*_2_). For each row we vary the value of *v*_2_: for the first row, *v*_2_ = 0.001; for the second row, *v*_2_ = 0.005; for the third row, *v*_2_ = 0.01. We then loop over 15 evenly-spaced values of *σ*_1_ ∈ [0.4, 0.8]. For each value of *σ*_1_, we perform 400 simulations (so each panel is 6 *×* 10^3^ simulations). For each simulation we randomly set the fitness variation for combinations of alleles (i.e., the values of *δ*^{*i,j*}^), as well as the values of *θ* and *R*_0_; these are uniform random variables in the intervals *θ* ∈ [10^−4^, 0.5] and *R*_0_ ∈ [1.5, 18]. The first column shows how the sign of 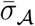 affects the equilibrium value of *D*^*′*^, and the second column shows the distribution of fitness variation used in the simulations. The red line is the rolling average of *D*^*′*^ using a window of size 400. All simulations assume *σ*_0_ = 1, *σ*_2_ = 0.2, *b* = *d* = 0.01, *α* = 0 and *ρ* = 10^−8^.

When we simulate the dynamics (Fig. S7 and Fig. S8), we observe two differences from the previous predictions.

**Figure S7:**
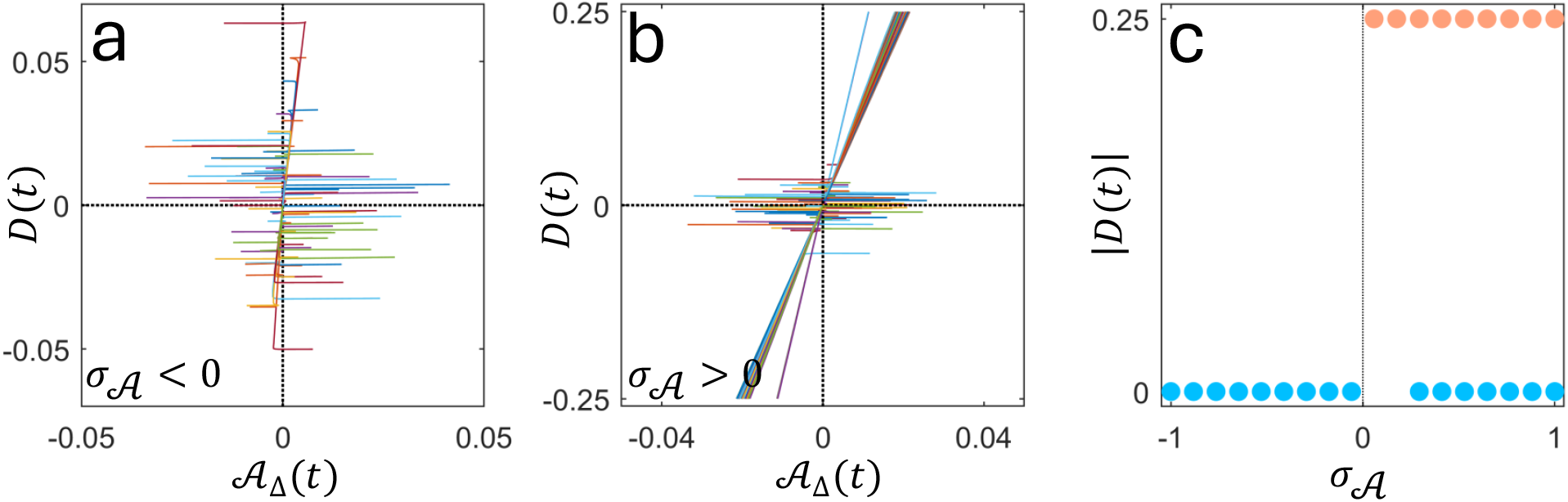
Disrupting the link between strain frequencies and the immunological landscape through the loss of immunity. In this example, host lose immunity to different antigens at independent rates *ω* = 0.2. This can disrupt the expected coupling between *D*(*t*) and 𝒜_Δ_(*t*) (panel **a**,**b**), reversing the predictions about the effects of the sign of *σ*_𝒜_ for the formation of PSS or no PSS (panel **c**), depending upon the value of *θ* and *R*_0_. Each circle in panel **c** represents 10^2^ simulations in which we loop over 10 evenly spaced values of *R*_0_ ∈ [1.5, 10] and 10 evenly spaced values (on log-scale) of *θ* ∈ [10^−4^, 0.39]. All simulations assume *b* = *d* = 0.01, *ρ* = 10^−8^, *σ*_0_ = 1, *σ*_2_ = *σ*_*h*_ = 0, and *α* = 0.

1. If immunity wanes in an antigen-by-antigen fashion (i.e., *ω >* 0), then strain structure may not form when *σ*_𝒜_ becomes large (Fig. S7). The logic here is as follows. When *σ*_𝒜_ is large, *σ*_1_, *σ*_2_ → 0. Consequently, hosts with immunity to one strain (say *AB*) can be infected by, at most, the other strain in the set of strains {*ab, AB*}, that is, strain *ab*. When this occurs, this produces hosts with immunity to all four antigens and thus full protection from infection. In order for these hosts to then become susceptible again, they have to lose immunity to one antigen at each locus. Since immunity to each antigen is lost randomly, such hosts are equally likely to produce hosts that are susceptible to members of one set of strains or the other, irrespective of which set of strains is overrepresented. However, as these hosts become susceptible to infection, the set of overrepresented strains will infect and deplete its available hosts faster. This creates an excess of hosts with immunity to the set of underrepresented strains, meaning that 𝒜_Δ_(*t*) can have the opposite sign to LD. Since *σ*_𝒜_ → *σ*_0_ *>* 0, epistasis in fitness will then also have the opposite sign to population LD. Thus, strain structure may not form, even if *σ*_𝒜_ is large. Note that in Figure S7, these predictions are sensitive to the epidemiological parameters *θ* and *R*_0_, because these parameters shape the magnitude of the impact of a fixed rate of waning immunity (in all simulations in Figure S7, *ω* = 0.2).
2. Suppose that following infection, hosts acquire immunity to both antigens of a strain with probability 1 −*p*, or only a single antigen with probability *p* (each antigen is chosen with probability 1*/*2). If *p* is sufficiently large, strain structure occurs only when *σ*_𝒜_ *<* 0 (Fig. S8).

The logic here is as follows. Suppose that the set of strains {*ab, AB*} is overrepresented in the population (*D*(*t*) *>* 0). Here, we would expect naive hosts to be more likely to become infected by either strains *ab* or *AB*. Following infection by strain *AB* (say), the host is then most likely to become infected by strain *ab* (since it is more abundant). However, each infection can provide immunity to at most a single antigen (since *p* → 1). Thus, when the two randomly selected antigens are at different loci, following a secondary infection, the host will then have immunity to either the antigens *A, b* or *a, B* from the set of underrepresented strains, {*Ab, aB*}. Consequently, 𝒜_Δ_(*t*) will have the opposite sign to *D*(*t*) and more hosts will have immunity to the set of underrepresented strains than to the set of overrepresented strains. If *σ*_𝒜_ *<* 0, this provides an advantage to the set of overrepresented strains, such that PSS can occur.

**Figure S8:**
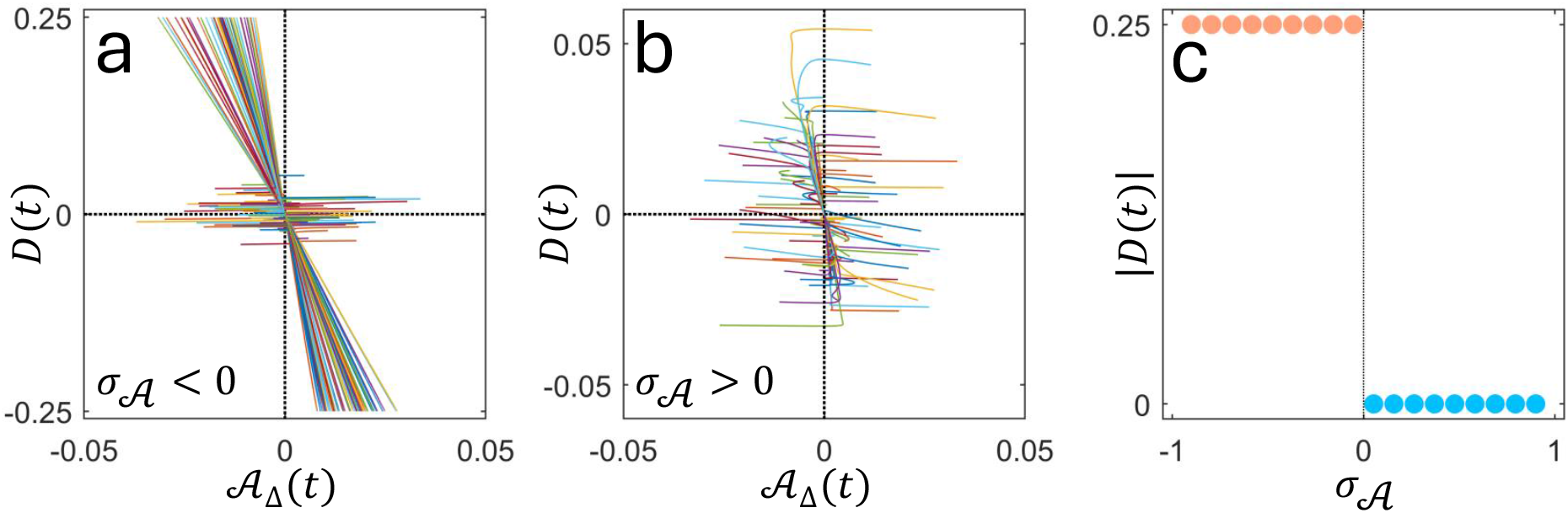
Disrupting the link between strain frequencies and the immunological landscape through the gain of immunity. In this example, following infection, hosts acquire immunity to a single, randomly chosen antigen (i.e., *p* = 1). This disrupts the expected coupling between *D*(*t*) and 𝒜_Δ_(*t*) (panel **a**,**b**), reversing the predictions about the effects of the sign of *σ*_𝒜_ for the formation of PSS or no PSS (panel **c**). Each circle in panel **c** represents 10^2^ simulations in which we loop over 10 evenly spaced values of *R*_0_ ∈ [1.5, 10] and 10 evenly spaced values (on log-scale) of *θ* ∈ [10^−4^, 0.39]. All simulations assume *b* = *d* = 0.01, *ρ* = 10^−8^, *σ*_0_ = 1, *σ*_2_ = *σ*_*h*_ = 0, and *α* = 0.

Note that these predictions are sensitive to how immune protections are modelled. In particular, when immunity reduces susceptibility, a host will typically only gain immunity to the infecting strain when it is successfully infected [15]. If, instead, immune protections affect, for example, transmissibility, even an ‘unsuccessful’ infection (i.e., an infection that is incapable of forward transmission) will update the host’s immunity. These differences are not expected to have a large effect, at least in models with simple epidemiological assumptions [15]. However, as the underlying epidemiological model becomes more complex, the evolutionary dynamics can become more sensitive to the type of immune protection, as the above examples illustrate.

### S8 Variation in allelic diversity across loci

The model featured in Section S3 assumes that there is an equal number of alleles at each locus. This symmetry allowed for the separation of fast and slow timescales, as all alleles rapidly converged to the same frequency, at which point we could trace the evolution of LD without needing to account for further allele frequency change. Here, we consider what happens if at one locus there are two alleles *i* ∈ {*A*_1_, *A*_2_}, and at the other there are three, *j* ∈ {*B*_1_, *B*_2_, *B*_3_}. We obtain six possible strains,

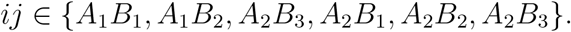

Without loss of generality, we choose strain *A*_1_*B*_1_ as the reference strain. Here, the evolutionary variables are the frequency of alleles *A*_2_ and *B*_*j*_ at time *t*, which are

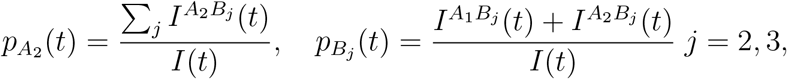

and the pairwise LD between alleles *A*_2_ and *B*_*j*_,

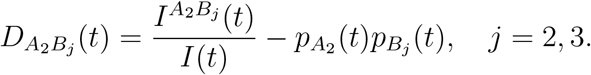

Next, the three additive selection coefficients are defined as

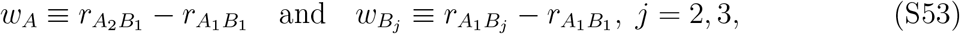

and the two terms specifying epistasis in fitness are

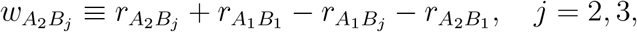

that is,

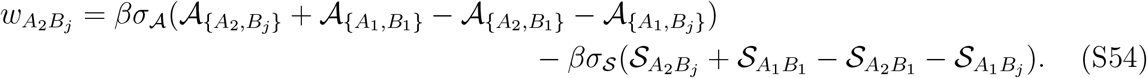

Here, as before, 𝒜_{*i*}_ and 𝒜_{*i,j*}_ denote the density of all uninfected hosts with immunity to at least the antigen *i* and antigens *i* and *j*, respectively, and 𝒮_*ij*_ denotes the density of uninfected hosts with homologous immunity to at least strain *ij*.

It is then straightforward to write out the equations describing the evolutionary dynamics. Here we focus on interpreting the simulated dynamics. The key is to recognize that additive selection favours the frequency of each allele at a given locus to be equal, that is, 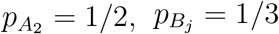, whereas epistatic selection either favours LD to increase or decrease in magnitude.

This depends upon whether *D*_*ij*_(*t*) and *w*_*ij*_ are of the same or opposite signs. In the two-allele, two-locus model of Section S3, allele frequencies could be equal *and* LD could assume any value (depending upon epistatic selection). As a consequence, two timescales emerged: a fast timescale on which allele frequencies were driven to equilibrium (*p*_*k*_ → 1*/*2), and a slow timescale on which LD evolved (see equation (S25)). When there are more alleles at one locus than the other, this need not be true. In this case, competition between epistatic and allelic selection can induce complex dynamics.

To illustrate this, we first consider the case in which *σ*_𝒮_ = 0. As was true for the model of Section S3, there is a single epidemiological feedback on epistasis. Thus, oscillatory evolutionary dynamics are not possible and the population will converge to an evolutionary equilibrium. If *σ*_𝒜_ *<* 0, then epistasis favours linkage equilibrium, 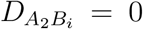, whereas additive selection favours 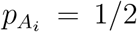 and 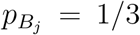. Since these can be simultaneously achieved, there is no conflict between epistasis and additive selection (Fig. S9). However, when *σ*_𝒜_ *>* 0, epistasis favours pairwise LD evolving towards *D*^*^. However, this cannot be achieved for all three {*A*_2_, *B*_*j*_} pairs if the population is in a state favoured by additive selection (i.e., 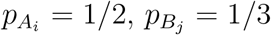). This creates a conflict between epistasis and additive selection, and so the evolutionary outcome depends upon their relative strengths. If *σ*_*A*_ is positive but small, epistasis will be relatively weak compared to additive selection. This will cause the allele frequencies to diverge from their expectation, but 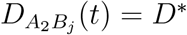 for two of the three {*A*_2_, *B*_*j*_} pairs; the third pair will be in linkage equilibrium (so two genotypes will have been eliminated from the population). As *σ*_*𝒜*_ increases, the allele frequencies diverge further from their allele-frequency based optimum, that is, the distance (measured here using the Euclidean norm)

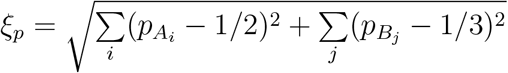

becomes increasingly large (Fig. S9), while the pairwise LD becomes closer to *D*^*^, that is,

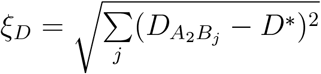

becomes increasingly small (Fig. S9). Eventually, when *σ*_𝒜_ is sufficiently large, 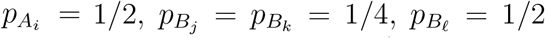 for *j* ≠ *k* ≠ *ℓ* = 1, 2, 3, and 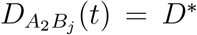 for each pairwise combination (so three genotypes will have been eliminated from the population) (Fig. S9). Thus, the sign of *σ*_𝒜_ still determines whether PSS forms or not, but now its strength additionally affects the amount of PSS observed. Importantly, however, even if *σ*_𝒜_ is maximized, the PSS observed will not be equivalent to the DSS of Gupta et al [6, 12].

Second, suppose that *σ*_𝒮_ *>* 0. As was the case for two diallelic loci, evolutionary oscillations are possible. But there is an important difference: because there is no longer a fast and a slow timescale, the allele frequencies will *not* reach an equilibrium. Both LD and the allele frequencies will oscillate (Fig. S10).

### S9 More than two loci

What happens as the number of loci increases? To illustrate this, we suppose there are three diallelic loci, {*a, A*}, {*b, B*} and {*c, C*}. Without loss of generality, choose strain *abc* as the reference strain. Here, the evolutionary dynamics consist of the dynamics of the allele frequencies, *p*_*k*_(*t*), *k* ∈ {*A, B, C*}, e.g.,

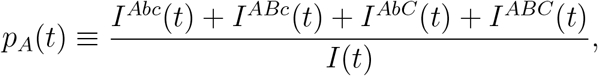

three pairwise LD terms, *D*_*ij*_(*t*), *ij* ∈ {*AB, AC, BC*}, e.g.,

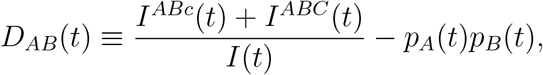

and triple LD,

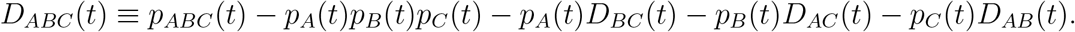

**Figure S9:**
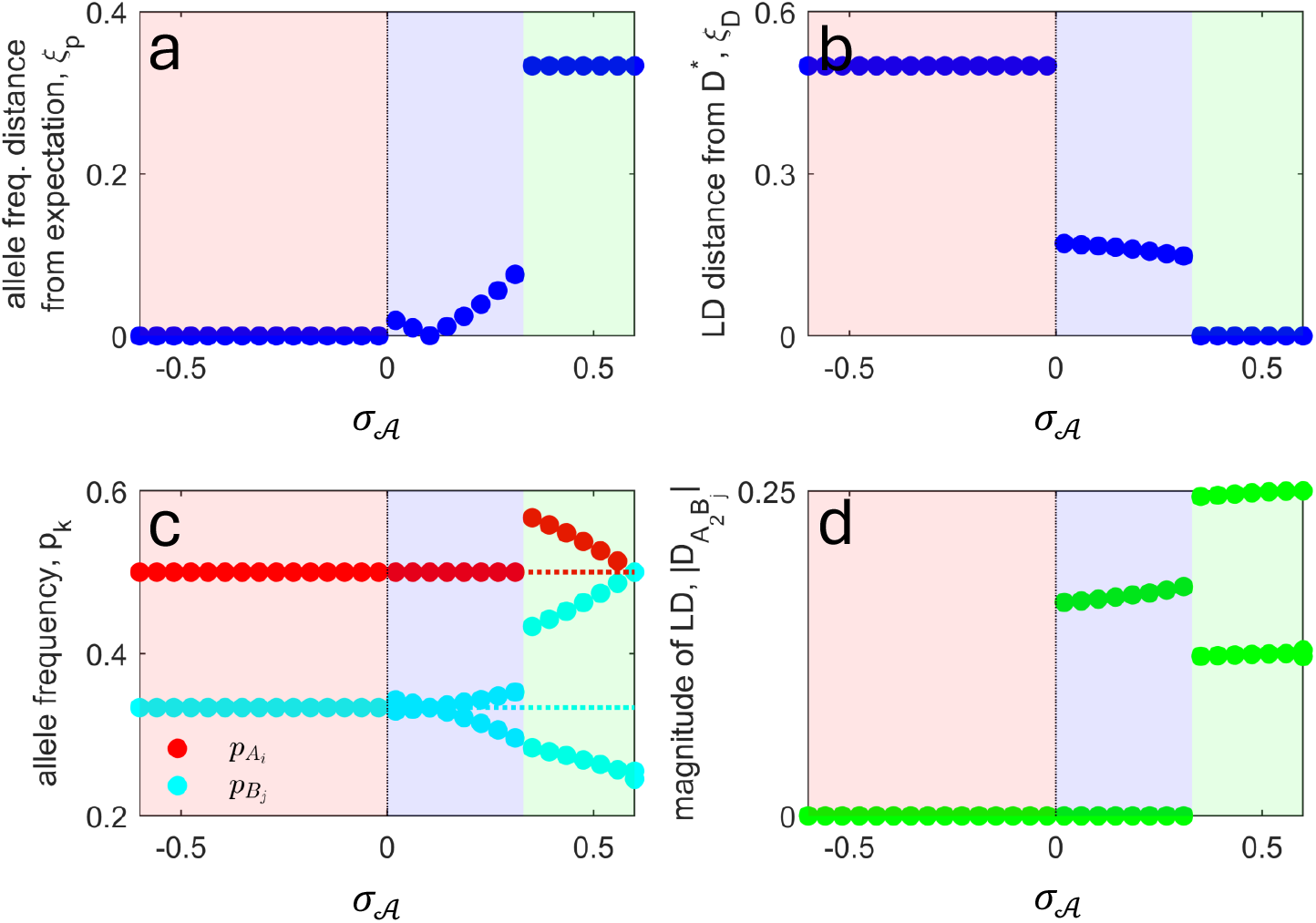
Consequences of variation in allelic diversity across loci. An unequal number of alleles across loci can produce conflict between selection on alleles (additive selection coefficients) and on strains (epistasis) because allele frequencies cannot be at their expected values 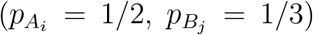 *and* the LD between each {*A*_2_, *B*_*j*_} pair satisfy 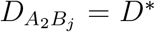. If *σ*_𝒜_ *<* 0 (red shaded region), there is no conflict because epistasis does not favour strain structure. If *σ*_𝒜_ *>* 0 but weak (blue shaded region), although PSS is favoured, selection on alleles is stronger than on strains and so there is no LD between one of the {*A*_2_, *B*_*j*_} pairs. As *σ*_𝒜_ increases further (green shaded region), so does the strength of epistasis; this pushes all three 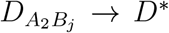 (panel **b**) and allele frequencies away from their expectation (panel **a**). Parameter values: *β* = 1, *μ* = 0.4, *b* = *d* = 0.01, *ρ* = 10^−8^, *σ*_0_ = 1, *σ*_2_ = *σ*_*h*_ = 0.4, and *σ*_1_ ∈ [*σ*_0_, *σ*_2_].

**Figure S10:**
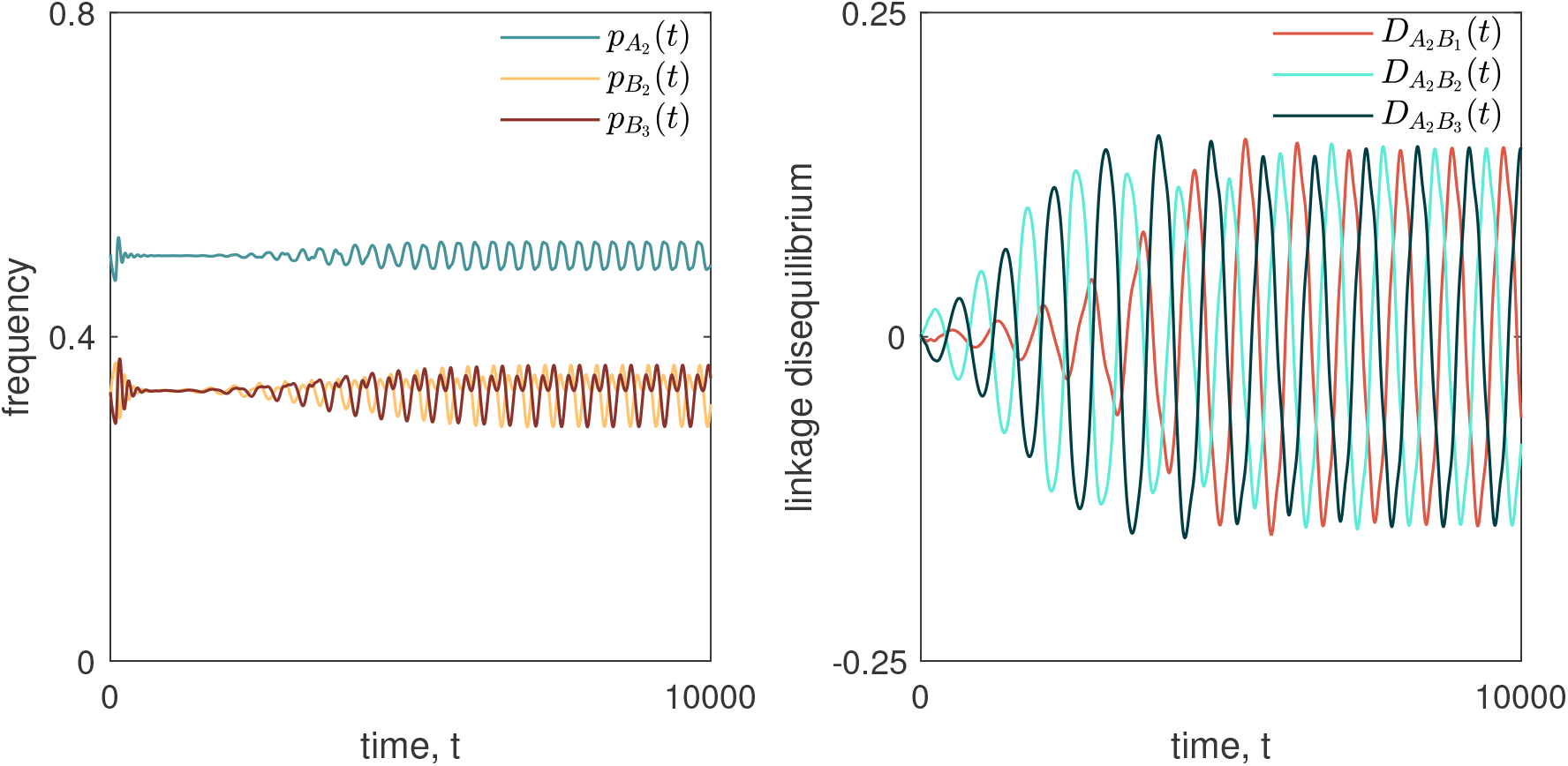
Conflict between selection on alleles and strains generates complex evolutionary oscillations. With variation in allelic diversity across loci, there is no longer a separation between fast and slow timescales. This is because allele frequencies cannot be at their expectation *and* all pairwise LD terms reach *D*^*^. Consequently, when evolutionary oscillations occur, these involve both allele frequencies and LD. Parameter values: *β* = 1, *μ* = 0.4, *α* = 0, *b* = *d* = 0.01, *ρ* = 0, *σ*_0_ = 1, *σ*_1_ = 0.3, *σ*_2_ = 0.15, and *σ*_*h*_ = 0.

Limiting our attention to immune protections that reduce susceptibility, from the model of Section S1, the per-capita growth rate of strain *ijk* is

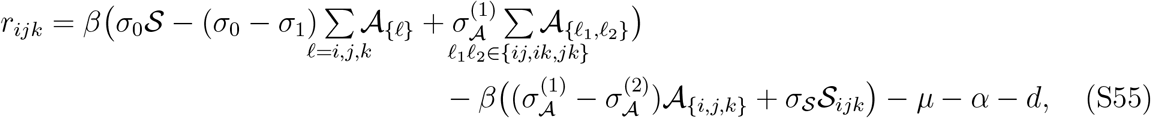

where *σ*_𝒮_ *≡ σ*_3_ − *σ*_*h*_, and

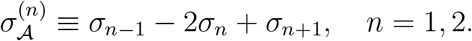

Thus, 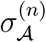 measures whether having antigenic immunity to *n* antigens of the infecting strain provides more, 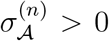, or less, 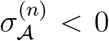, protection than would be expected based upon the protection offered by having immunity to *n* − 1 versus *n* + 1 antigens.

Using these per-capita growth rates, the additive selection coefficient for allele *A* (and similarly for alleles *B* and *C*) is

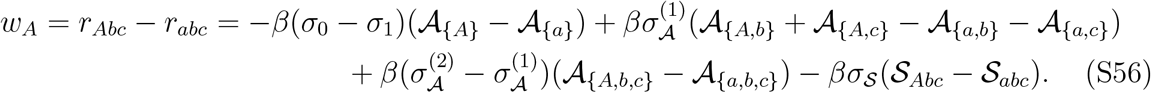

Pairwise epistasis for alleles *ij* ∈ {*AB, AC, BC*} is defined as before when using the allele at the third locus as the reference allele, *ℓ* ∈ {*a, b, c*}; for example,

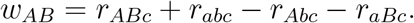

Using our per-capita growth rates, this can be written as

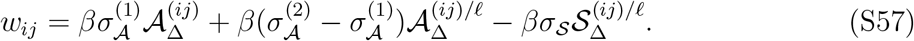

*w*_*ij*_ includes the following composite variables:

- 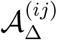 indicates whether more uninfected hosts have immunity to the antigen combinations *ij* and/or *îĵ* (i.e., 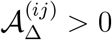), or immunity to the antigen combinations *îj* and/or *iĵ* (i.e., 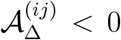). Note that this is irrespective of genetic background, and that î indicates the other allele at the indicated locus, e.g., if *i* = *A*, then *î* = *a*. For example,

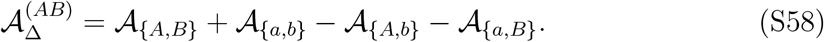
- 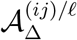 indicates whether more uninfected hosts with immunity to antigen *ℓ* also have immunity to the antigen combinations *ij* and/or îĵ (i.e., 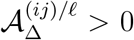), or immunity to the antigen combinations î*j* and/or *i*ĵ (i.e., 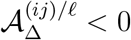). For example,

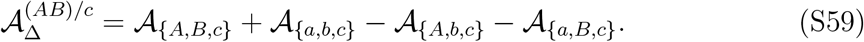
- 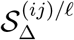 indicates whether more uninfected hosts have homologous immunity to members of the set of strains {*ijℓ*, îĵ*ℓ*} (i.e., 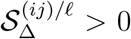), or homologous immunity to members of the set of strains {*i*ĵ*ℓ*, î*jℓ*} (i.e., 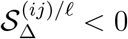). For example,

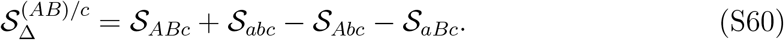

Finally, when there are three loci, we need to account for triple epistasis in fitness, denoted *w*_*ABC*_. Since by definition, *r*_*ABC*_ = *r*_*abc*_ + ∑_*k*_*w*_*k*_ + ∑_*ij*_ *w*_*ij*_ + *w*_*ABC*_, this is

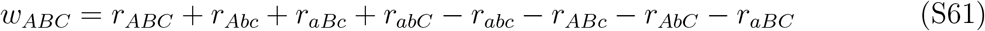

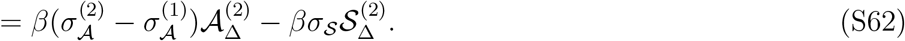

*w*_*ABC*_ includes the following composite variables:

- 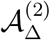 indicates if more hosts have full antigenic immunity to strains belonging to the set {*ABC, Abc, aBc, abC*} (i.e., 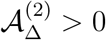), or strains belonging to the set {*abc, ABc, AbC, aBC*} (i.e., 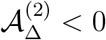). That is,

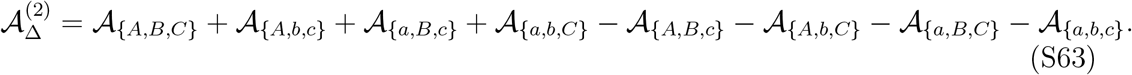
- 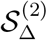 indicates if more hosts have homologous immunity to strains belonging to the set {*ABC, Abc, aBc, abC*} (i.e., 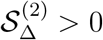), or strains belonging to the set {*abc, ABc, AbC, aBC*} (i.e., 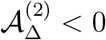). That is,

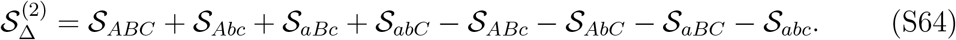

Note that the sets of strains {*ABC, Abc, aBc, abC*} and {*abc, ABc, AbC, aBC*} are characterized by each member having no more than one antigen in common with the other members.

This allows us to write down the equations describing the evolutionary dynamics. In what follows, we use these equations to extract general qualitative predictions. Note that our emphasis is on biological interpretability, so we do not attempt to rigorously characterize the dynamics.

#### S9.1 Fast and slow timescales, and different types of strain structure

We first observe that in the three-diallelic-locus model, there is also a fast timescale on which *p*_*k*_(*t*) → 1*/*2, and a slow timescale on which pairwise and triple-LD (strain structure) evolve. After the fast transient dynamics, on the slow timescale pairwise LD becomes

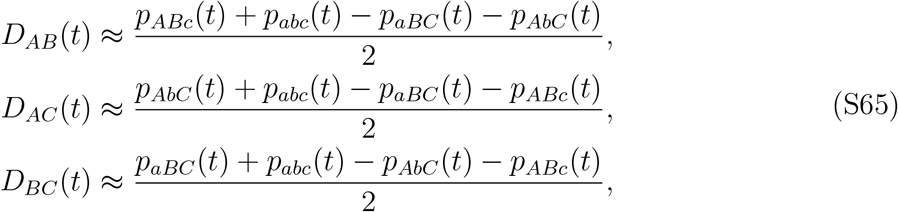

and triple LD becomes

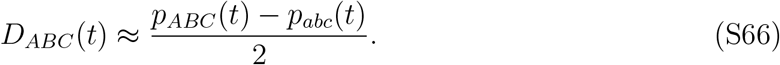

Inspection of these quantities allows us to identify two distinct ways in which strain structure can emerge:

1. When a set of strains whose members share no more than one allele with any other is overrepresented, that is, either set

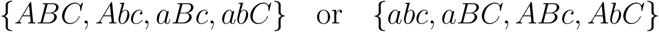

is overrepresented. This type of strain structure is driven by the build-up and maintenance of triple LD, and the removal of pairwise LD. Here, when strain structure is maximized (one set is competitively excluded), *D*_*ABC*_ (*t*) = *±*1*/*8 while *D*_*ij*_ (*t*) = 0. If we suppose there is no pairwise LD (*D*_*ij*_ (*t*) = 0), then, on the slow timescale

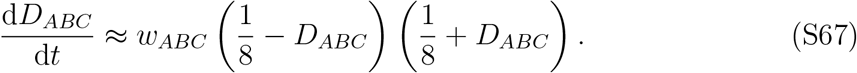 By inspection of equation (S67) we observe that in order for triple LD to build up and be maintained, triple LD (*D*_*ABC*_) and triple epistasis (*w*_*ABC*_) must share sign.
2. When one of the four sets of strains whose members have no alleles in common is over-represented, that is, one of the sets

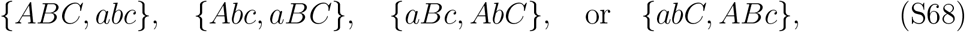

is overrepresented. This type of strain structure is driven by the build-up and maintenance of pairwise LD, and the removal of triple LD. Here, when strain structure is maximized (i.e., three of the sets of strains in (S68) are competitively excluded), then *D*_*ij*_(*t*) = *±*1*/*4, and *D*_*ABC*_ (*t*) = 0.

If we suppose that there is no triple LD (*D*_*ABC*_ (*t*) = 0), then on the slow timescale

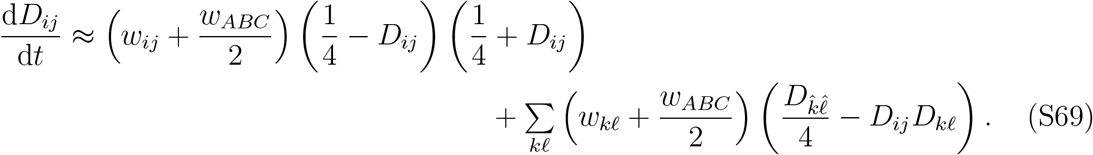

From equation (S69) we observe that it is no longer necessary that *w*_*ij*_ and *D*_*ij*_ must share sign for pairwise LD to be maintained. Instead, *w*_*ij*_ + *w*_*ABC*_ */*2 and *D*_*ij*_ need to share sign. Note that

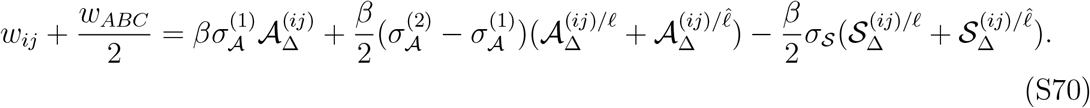

#### S9.2 Shape of the function controlling cross-protection

Next, suppose *σ*_𝒮_ = 0. Here, we study how the 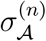 affect the evolutionary dynamics. Recall that 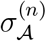 measures whether the rate of increase of cross-protection is locally saturating 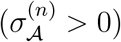 or accelerating 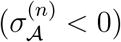 at *n*.

Two curves divide up the qualitatively different evolutionary behaviour in the (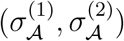) space. In what follows, we determine these two curves; the results are presented in Figure 5 in the main text. Throughout, we assume that the set of overrepresented strains also has more hosts with antigenic immunity to its members, that is, 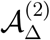 and *D*_*ABC*_ share sign, and 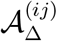 and *D*_*ij*_ share sign. We will not consider ways in which these relationships could break down as in Section S7.

##### Curve 1

*w*_*ABC*_ = 0. This occurs when 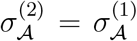, that is, the rate of increase of cross-protection is the same regardless of the number of antigens the host has immunity to. This partitions the parameter space between when triple LD can be maintained and when it cannot. In particular, since triple LD requires the signs of *w*_*ABC*_ and *D*_*ABC*_ to match, it can only be maintained if 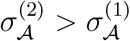.

##### Curve 2

*w*_*ij*_ + *w*_*ABC*_ */*2 = 0. From equation (S69), this curve specifies when pairwise LD can or cannot form. Since this curve depends upon the epidemiological dynamics in addition to the parameters 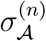, it is not straightforward to precisely specify the curve in (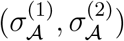) space. However, note that in equation (S70), we can write 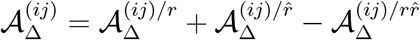, where

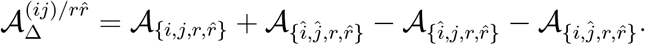

Each of these terms requires hosts to be infected by different pathogen strains. Hence, if we suppose that the frequency of hosts being infected multiple times is low compared to hosts being infected once, e.g., 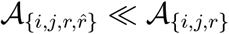 then 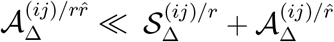. Here, equation (S70) becomes

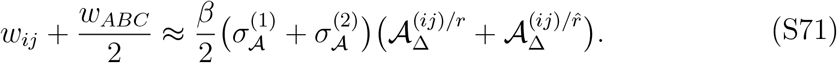

It follows that 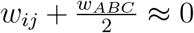 when 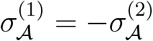, or *σ*_0_ − *σ*_1_ − *σ*_2_ + *σ*_3_ = 0. Since the maintenance of pairwise LD requires the signs of *w*_*ij*_ + *w*_*ABC*_ */*2 and *D*_*ij*_ to match, invoking the approximation of (S71), pairwise LD can be maintained if 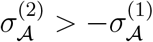, otherwise it cannot.

From consideration of these curves and Figure 5, notice that if both

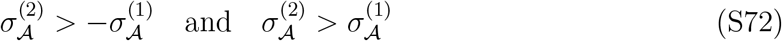

are satisfied, it is possible that pairwise LD can be maintained in the absence of triple LD. Moreover, triple LD can be maintained in the absence of pairwise LD. This creates the possibility of an evolutionary bistability: depending upon the initial conditions, the population can stably exhibit different types of structure, that is, either |*D*_*ABC*_ | is maximized and *D*_*ij*_ = 0, |*D*_*ij*_| is maximized and *D*_*ABC*_ = 0.

From our analysis above and Figures 5 and S11, increasing 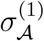 and/or 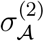, produces strain structure, but depending on which of these increases more rapidly, different types of structure can occur. Large 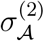 relative to 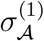 tends to favour triple LD, resulting in a population with four strains present, which share no more than one antigen with any other. If instead 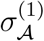 is large relative to 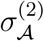, this favours pairwise LD, resulting in a population in which only two strains are present that share no antigens. The intuition behind this is as follows: if 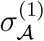 is positive and larger than 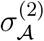, immune protections are much more effective against a single antigen than expected, and so the minimum amount of strain diversity that preserves antigenic variation is favoured. If 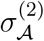 is instead large, then immune protections are more effective when strains share two antigens and relatively ineffective against strains sharing a single antigen. Consequently, greater strain (genotype) diversity is expected to be maintained in the population.

#### S9.3 *σ*_𝒮_ *>* 0 promotes oscillations and maintains diversity

When *σ*_𝒮_ *>* 0, this has a similar effect for three loci than for two. In particular, *σ*_𝒮_ *>* 0 increases the likelihood that *w*_*ABC*_ and *w*_*ij*_ + *w*_*ABC*_ */*2 do not share sign with *D*_*ABC*_ and *D*_*ij*_, respectively. Thus, *σ*_𝒮_ *>* 0 breaks up strain structure and promotes genotypic diversity. Likewise, by introducing a second epidemiological feedback through hosts with homologous immunity (hosts belonging to either 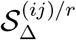(for *w*_*ij*_) or 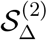 (for *w*_*ABC*_)), *σ*_*S*_ *>* 0 can produce evolutionary oscillations with any level of strain structure, during which the set of overrepresented strains varies (Fig. S12). Simulations suggest that these oscillations do not occur between different types of structure in the deterministic system, even when bistability is possible. However, in the presence of demographic stochasticity, it seems likely that this can occur, which would further complicate the dynamics.

**Figure S11:**
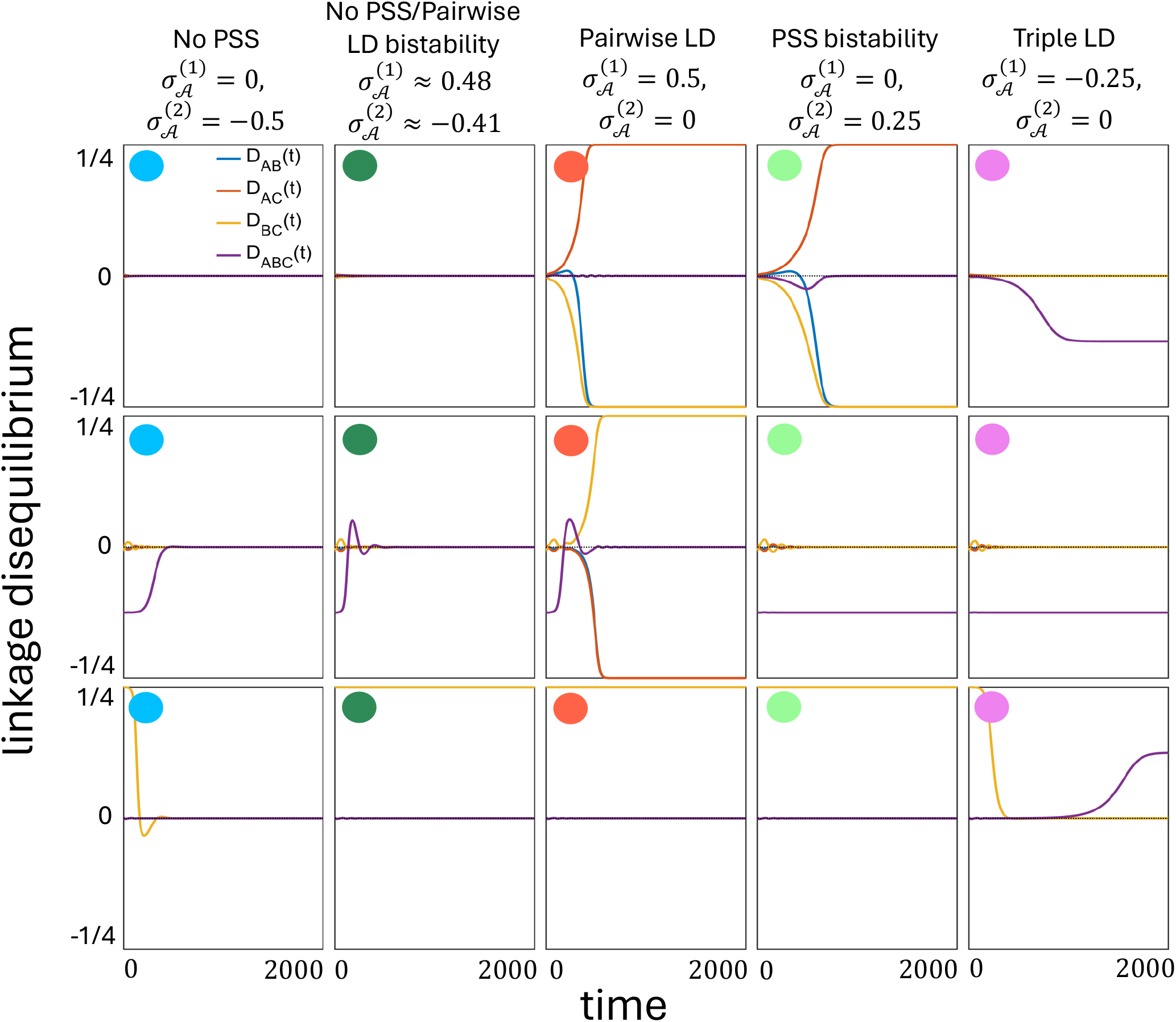
Dynamical behaviour of LD as 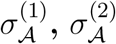, and the initial conditions vary. The dynamics of pairwise- and triple LD (*D*_*ij*_(*t*) and *D*_*ABC*_ (*t*)) depend upon the values of 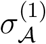 and 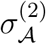 (different columns) and the initial conditions (different rows). Each column corresponds to the indicated values of 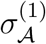 and 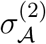, with the coloured circle indicating the approximate position of this scenario in Figure 5. Each row corresponds to a different initial condition. In the first row, all simulations start near a state of no PSS; in the second row, all simulations start near a state of triple LD PSS; in the third row, all simulations start near a state of pairwise LD PSS. All simulations assume *b* = *d* = 0.01, *β* = 1, *μ* = 0.4, *ρ* = 10^−6^, *σ*_0_ = 1 and *σ*_3_ = *σ*_*h*_ = 0; *σ*_1_ and *σ*_2_ vary as indicated by column.

**Figure S12:**
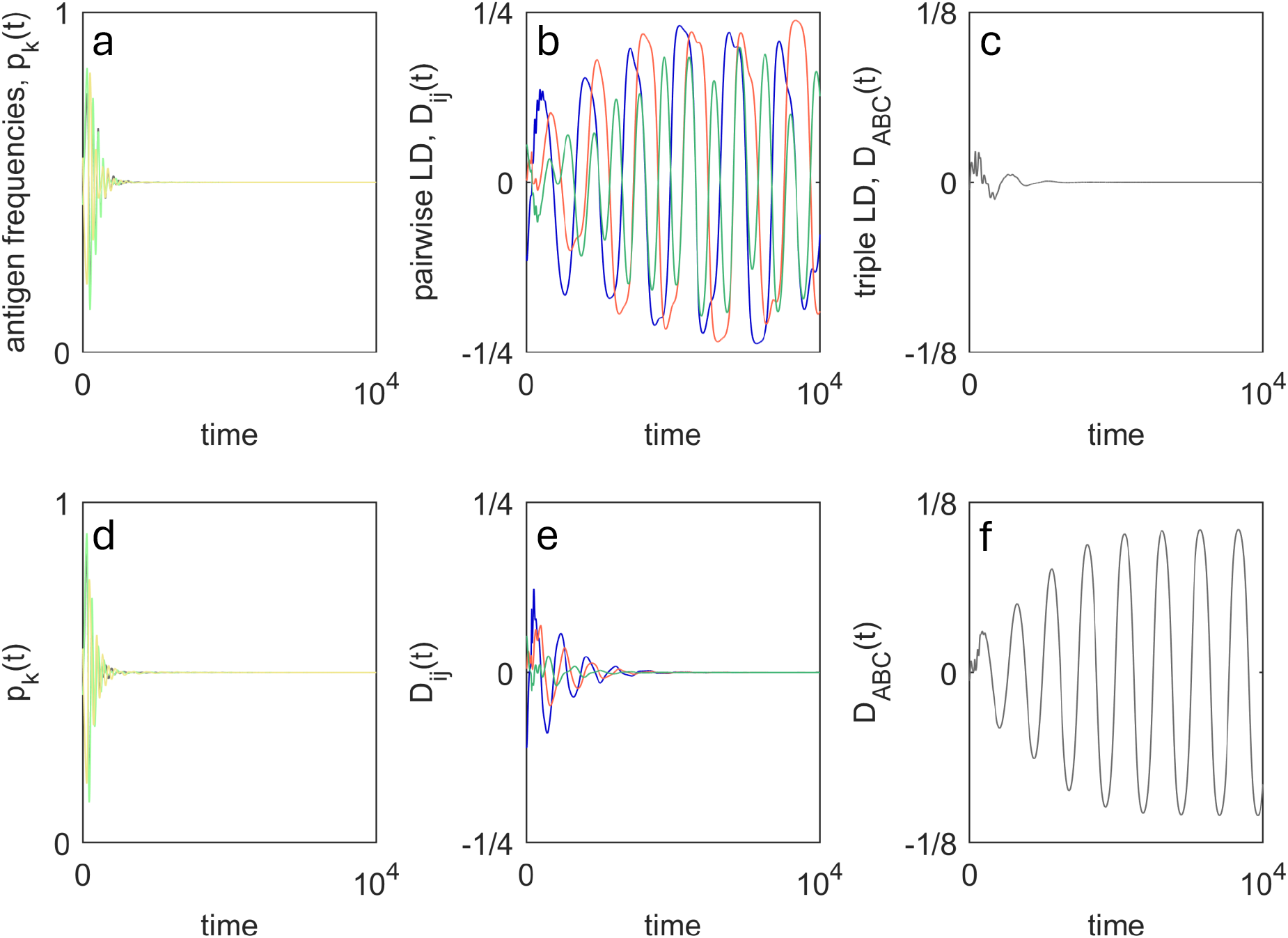
Evolutionary oscillations for more than two loci. For more than two loci there is more than one possible order of LD, and consequently, there are different ‘types’ of PSS. As a result, evolutionary oscillations can occur for any order of LD. All simulations assume *b* = *d* = 0.005, *β* = 1, *μ* = 0.5, *ρ* = 10^−8^, *σ*_0_ = 1 and *σ*_*h*_ = 0.05. In panels **a**-**c**, *σ*_1_ = 0.6, *σ*_2_ = 0.25, *σ*_3_ = 0.2; in panels **d**-**f**, *σ*_1_ = 0.7, *σ*_2_ = 0.3, *σ*_3_ = 0.2.

